# Simple synaptic modulations implement diverse novelty computations

**DOI:** 10.1101/2023.08.16.553635

**Authors:** Kyle Aitken, Luke Campagnola, Marina Garrett, Shawn Olsen, Stefan Mihalas

## Abstract

Since environments are constantly in flux, the brain’s ability to identify novel stimuli that fall outside its own internal representation of the world is crucial for an organism’s survival. Within the mammalian neocortex, inhibitory microcircuits are proposed to regulate activity in an experience-dependent manner and different inhibitory neuron subtypes exhibit distinct novelty responses. Discerning the function of diverse neural circuits and their modulation by experience can be daunting unless one has a biologically plausible mechanism to detect and learn from novel experiences that is both understandable and flexible. Here we introduce a learning mechanism, *familiarity modulated synapses* (FMSs), through which a network response that encodes novelty emerges from unsupervised multiplicative synaptic modifications depending only on the presynaptic or both the pre- and postsynaptic activity. FMSs stand apart from other familiarity mechanisms in their simplicity: they operate under continual learning, do not require specialized architecture, and can distinguish novelty rapidly without requiring feedback. Implementing FMSs within an experimentally-constrained model of a visual cortical circuit, we demonstrate the generalizability of FMSs by reproducing three distinct novelty effects recently observed in experiments: absolute, contextual (or oddball), and omission novelty. Additionally, our model reproduces functional diversity within cell subpopulations, leading to experimentally testable predictions about connectivity and synaptic dynamics that can produce both population-level novelty responses and heterogeneous individual neuron signals. Altogether, our findings demonstrate how simple plasticity mechanisms within the cortical circuit structure can give rise to qualitatively distinct novelty responses. The flexibility of FMSs opens the door to computationally and theoretically investigating how distinct synapse modulations can lead to a variety of experience-dependent responses in a simple, understandable, and biologically plausible setup.

## 1 Introduction

Brains of complex organisms contain internal representations of the world that are shaped by stimuli they have become familiar with over time. Since their environment can change rapidly, an organism’s survival can be dependent upon its ability to quickly identify novel stimuli. Indeed, over decades of study, effects of stimulus novelty have been found throughout the brain and are known to occur over many timescales [1–5] These effects vary from internal changes such as promoting learning and memory to behavioral adjustments including changes to perception, attention, and exploration [2–4]. Across sensory modalities and species, novel stimuli are generally associated with an increased response relative to their familiar counterparts [2, 3]. Such novelty-responses (or their inverse, familiarity-responses) have been observed in cortical, subcortical, and neuromodulatory areas of the brain at both an individual cell level [6–9] and across macroscopic cell populations [10–12]. Additionally, studies have distinguished responses to distinct types of novelty. For example, *absolute* novelty, when an organism is exposed to a previously unobserved stimulus [10, 13], is distinguished from *contextual* (or oddball) novelty, where a previously observed stimulus is novel only in the context of recently observed stimuli that may also occur from the *omission* of an expected stimulus [14–16].

The mammalian neocortex is believed to play an especially important role in modeling the world around us and thus how it responds to these various types of novel stimuli is of great interest [3]. Within the cortex, what is believed to be a general purpose disinhibitory circuit is repeated across different brain regions and species, and many recent experimental studies have elucidated the properties of the cells within this circuit [17–20]. Specifically, the structure of this cortical circuit is defined by connectivity between somatostatin (SST) and vasoactive/intestinal peptide (VIP) expressing inhibitory interneurons as well as pyramidal excitatory neurons [21]. This circuit is thought to facilitate novelty responses through mutual inhibition between the VIP and SST populations that provides a disinhibitory pathway from VIP to excitatory cells [22]. Recent experimental studies have found that novelty responses vary significantly across these distinct cell populations [23, 24]. These studies suggest that the enhanced response of VIP cells to novel stimuli suppresses the SST population’s response, releasing the local excitatory population from inhibition and leading to an increased excitatory novelty response.

Although broad cell classes are a useful simplification to understand the function of the cortical circuit, each class can be further divided into subclasses or types that differ in gene expression patterns, synaptic connectivity, electrical properties, and morphology [19, 25–28]. Indeed, within the excitatory, SST, and VIP cell populations, subpopulations that have distinct feature-coding across familiar and novel stimuli have been recently identified [23, 24]. Given these recent results, an open question is what biological mechanisms might allow populations to have such diversity in experience-dependent coding, and how this coding diversity relates to changes in the population’s macroscopic response to novel stimuli.

Since the observation of the brain’s ability to rapidly detect novel stimuli, computational models have been used to investigate how the brain might distinguish familiar representations and evoke distinct responses to unfamiliar stimuli [16, 29, 30]. Many of these models rely on modifications of synaptic connections to encode stimuli. For example, Hopfield networks can encode familiar stimuli via lateral connections and are capable of recalling said stimuli using recurrent activity [31]. However, many of these computational models require carefully placed synaptic connections to encode distinct memories [32, 33] or strict training and testing phases that do not reflect an organism’s natural behavior [32, 34], both of which limit their ability to be implemented into more general models. Additionally, some models rely on complex non-local credit assignment mechanisms that are biologically unrealistic to develop their novelty-responses [35, 36].

In this work, we introduce a mechanism that implements simple plasticity rules via synaptic modulations and is capable of adapting to stimuli through biologically-realistic local, unsupervised learning. Broadly, it relies on modulating the synapses that play a role in producing the output responses of familiar stimuli, and as such we refer to the mechanism as *familiarity modulated synapses (FMSs)*. A strength of FMSs is their simplicity and thus generality; we show FMSs can broadly represent various synaptic plasticity effects that occur over different timescales. We focused on parameterizing the FMSs such that they represent biologically realistic plasticity mechanisms such as long-term potentiation/depression (LTP/D) [37, 38] or short-term synaptic plasticity (STSP) [39]. FMSs can be implemented on a set of excitatory or inhibitory synapses feeding from one cell population to another whose strengths and connections are randomly drawn, meaning it requires essentially no specialized architecture and is thus straightforward to implement into more complex neural network models. The mechanism also requires no specific training regimen, simply becoming adapted to stimuli it has seen in recent history under continuous learning, similar to how biological organisms learn. We first establish properties of the FMSs in the simplest possible feedforward setting. Afterwards, we incorporate several distinct FMS mechanisms into a model of the visual cortical circuit, with connectivity properties constrained from multi-patch synaptic physiology studies [20], relative cell counts from in situ hybridization experiments [17, 18], and additional cell properties from electrophysiology recordings [19]. We demonstrate the generalizability of the FMSs by modelling three distinct novelty effects: absolute [10, 13], contextual (oddball) [14, 15], and omission novelty [16]. Although each of these novelty effects has been studied in isolation, recent studies in the visual cortical circuit of mice investigate how distinct cell populations respond to all three types of novelty [23, 24]. The flexibility of FMSs allows for us to simultaneously capture the three novelty effects within our experimentally-constrained model of the cortical circuit, while also reproducing the diverse subpopulation coding seen in the same experiments [24].

### Related works

Many existing models of novelty detection rely on modifications of synaptic connections in order to encode familiar stimuli, but often require specialized connection architectures in order to encode distinct memories [32, 33], do not operate under a continual learning setting [22, 32, 34, 40, 41], or rely on complex non-local credit assignment [35, 36, 42], all of which the FMSs avoid. Refs. [40, 41] consider how a firing-rate dependent learning rule, directly derived from passive and dimming-detection experiments, can match time-averaged and time-dependent responses. Feedforward adaptation as a means of repetition suppression has been previously studied previously [22, 29, 35, 42–44] and is advantageous because it does not require convergence to a steady state or feedback-dependent activity to distinguish stimuli [45]. Novelty responses on an image change detection task were reproduced using STSP-like synaptic modulations [29, 42]. The specific form of the synaptic modulations used in this work are an unsupervised version of those described in Refs. [35, 36] that originated in the learning-how-to-learn machine learning literature [46].

Many other computational models of the visual cortical circuit have been built to understand the individual cell population effects of disinhibition and how the circuit’s activity might change over learning [18, 22, 47–54]. While many models of cortical circuits treat inhibitory interneurons as a unitary population [47, 54], more recent models have incorporated the diversity of interneuron populations, including the VIP-SST-Excitatory disinhibitory circuit [18, 22, 48–53]. Ref. [52] studies and models the VIP-SST-Exc. disinhibitory circuit in L2/3 of mice in the setting of visual context modulation and finds contextual modulation is unlikely to be inherited from L4 and thus may rely on local circuitry. A computational model of the cortical circuit constrained by electrophysiological studies that incorporates population diversity and inhibitory plasticity was recently used to study prediction errors in Ref. [48, 53]. Although they also investigate how connectivity influences the development of neuron subpopulations, the training/testing stimulus sequences are different from the ones we investigate here.

### Setup: familiarity modulated synapses

In this work we consider networks of firing-rate, point-like excitatory and inhibitory neurons that can influence one another through synapses that we represent using weight matrices. Let **W** represent a set of *fixed* synapses that connects a presynaptic population of neurons to a postsynaptic population, with firing-rates at time *t* represented by the vectors 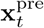 and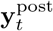, respectively (Fig. 1a, left). For example, the postsynaptic population’s firing-rates may be related to the presynaptic population’s activity via 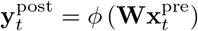 where *ϕ* (·) ≥ 0 is a non-linear function that accounts for the postsynaptic neurons’ properties such as their firing threshold and maximum firing rate. We take **W** to be sparse and, for simplicity, take the nonzero weights to be drawn from a normal distribution. Furthermore, the sign of the nonzero elements of **W** are fixed by the cell-type of the presynaptic population: excitatory neurons only have positive-weight synapses so that they increase postsynaptic potentials and inhibitory neurons only have negative-weight synapses.

**Figure 1:**
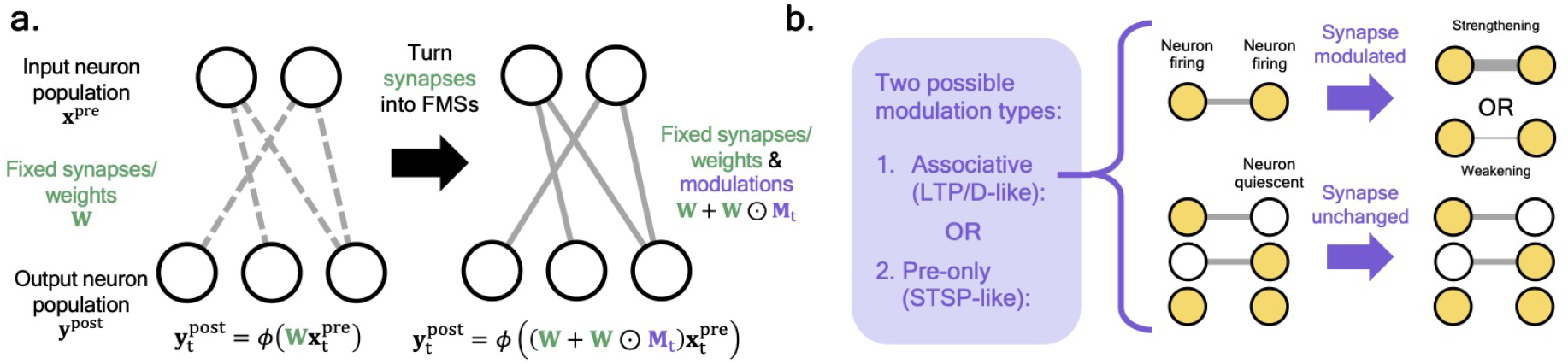
Familiarity modulated synapses. **(a)** On the left, an exemple feedforward firing-rate network, where a population of (firing-rate) input neurons, **x**^pre^, influences a population of output neurons, **y**^post^, through a set of fixed synaptic connections, **W**. On the right, the fixed synaptic connections are modified to become familiarity modulated synapses (FMSs), i.e. **W** → **W** + **W** ⊙ **M**_*t*_, allowing each synapse’s strength to be modulated over time via the matrix **M**_*t*_. **(b)** The two types of modulations we consider in this work: (1) associative and (2) pre-only dependent. See Eq. (2) for explicit expressions. For an associative update rule, examples of how the behavior of neurons influences the way their synapses are modulated (see Fig. S1a for equivalent pre-only diagram). In short, the modulations will either strengthen (*η >* 0) or weaken (*η <* 0) the neuron connections if both the pre- and postsynaptic neuron are firing and a synaptic connection already exists between said neurons.

We modify the fixed weights to be *familiarity modulated synapses* (FMSs) by taking

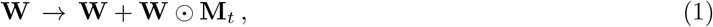

where **M**_*t*_ represents time-dependent modulations to the synapses represented by **W** and ‘⊙’ is the elementwise product. In our exemplar network, the relation between pre- and postsynaptic activity would be 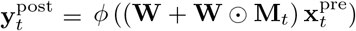 (Fig. 1a, right). We will investigate two distinct modulation mechanisms throughout this work that determine how **M**_*t*_ evolves in time,

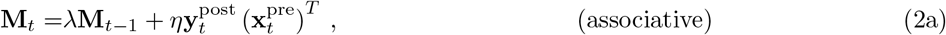

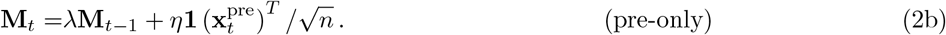

Both rules are completely unsupervised and modulate based on only information locally available to the synapse. The *associative* update, Eq. (2a), is the more general modulation rule dependent upon both the post- and presynaptic neuron firing rates at time *t* (Fig. 1b). The parameter 0 *< λ <* 1 controls how quickly the modulations return to their baseline values, while |*η*| determines the size of the updates. Importantly, the sign of *η* controls the sign of **M** and thus whether synapses are strengthened or weakened by the modulations, i.e. if their magnitude increases or decreases, respectively. The *pre-only* modulation update expression, Eq. (2b), is only dependent on the presynaptic firing rate, in which case the dependence on 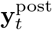 is replaced with **1**, the all 1’s vector, and normalized by the square root of the number of output neurons *n*.

Throughout this work, all **W** are fixed and thus the total synapse strength is only modified through the **M**_*t*_ term. “Training” will refer to the time period where a network is exposed to certain stimuli and its synapses are modified *solely via the unsupervised FMSs* described above. Crucially, we do not allow the modulations to change whether a synapse is excitatory or inhibitory, i.e. if *W*_*ij*_ ≥ 0 then *W*_*ij*_ + *W*_*ij*_*M*_*t,ij*_ ≥ 0 for all time. For simplicity, we also do not allow for new synapses to form, i.e. a synapse that doesn’t exist at initialization cannot be modulated.

Biologically, we envision the modulations as various mechanisms leading to changes in the synapses that occur over varied timescales and biological mechanisms. The associative mechanism, Eq. (2a), could broadly represent long timescale synaptic changes resulting from LTP/D mechanisms. Long term potentiation or depression of said synapses can be implemented by changing the sign of the learning rate, *η*. Meanwhile, the modulations that are only presynapse-dependent, Eq. (2b), could represent faster modulation mechanisms such as STSP. With these biological mechanisms in mind, we limit the size of the modulations such that they do not exceed synaptic changes that have been observed in experiment (see Methods for additional details).

## 2 Results

### 2.1 A simple, unsupervised, feedforward novelty-detector

To explore some basic properties of the FMSs, we first investigate their effect in a simple feedforward network that we show develops distinct responses to stimuli it has been exposed to before, what we refer to as *familiar* stimuli throughout this work. Many of the results we establish in the simple network with a single FMS mechanism generalize to the visual cortical circuit model we discuss afterwards in Sec. 2.2 with several distinct FMS mechanisms.

We represent the neuronal encodings of stimuli using distinct sparse random binary vectors (Fig. 2a, Methods). Prior to training, we draw two sets of 8 stimuli from this distribution. During training, the stimuli from what becomes the familiar set will be exposed to the network while its weights undergo unsupervised updates via an FMS mechanism. After training, we will compare the network’s response to the familiar set and the other set that was held out during training, what we refer to as the *novel* set. Noise is added to all input stimuli throughout this work (Methods).

**Figure 2:**
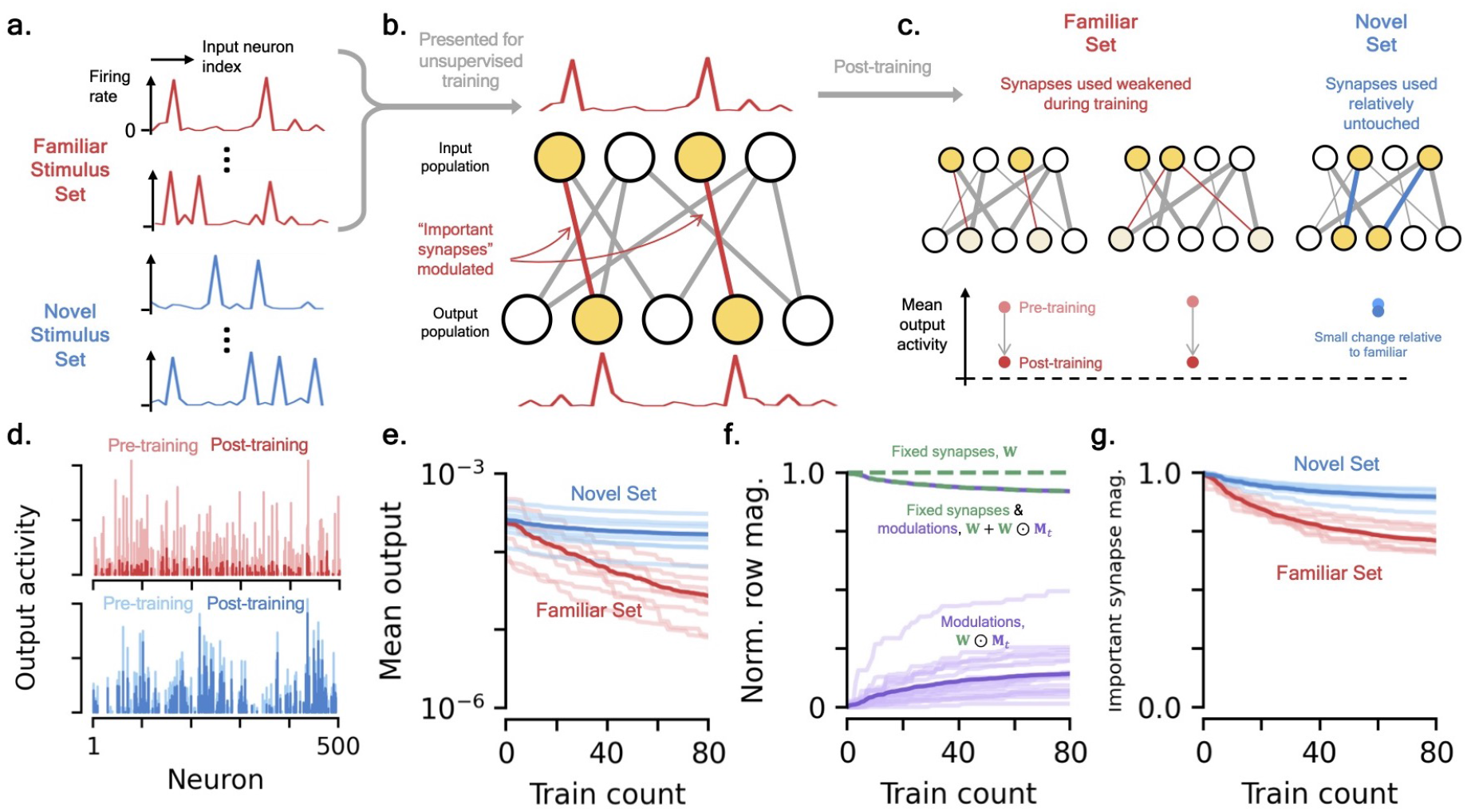
Familiar modulated synapses in a simple network. **[a-c]** *Schematic of network behavior and exposure to familiar set*. **(a)** The familiar (red) and novel (blue) sets of stimuli that excite the input neuron population are drawn from the same distribution, random sparse binary vectors with added noise. **(b)** We consider a simple two-layer network with FMSs connecting an excitatory input population to an output neuron population. At each time step a randomly chosen familiar stimulus excites the input population and, through the modulated synapses, causes the output population to fire in some pattern. For the example considered in this figure, the associative modulations weaken any synapses that connects a pre- and postsynaptic neuron that both fired for the given familiar stimulus, e.g. an effect that could arise from LTD. **(c)** After training, many of the network’s synapses have been modulated, changing its output behavior. The familiar set’s mean output activity is reduced relative to their pre-training activity. The post-training mean output activity of the novel set is relatively unchanged. **[d-g]** *Results from example network and training. In this example, there are* 8 *familiar and* 8 *novel stimuli. Each familiar stimulus has been input into the network* 10 *times (shuffled order) for* 80 *training steps total*. **(d)** Example raw output response activity for a familiar (red) and novel (blue) stimulus pre- and post-training. **(e)** Change in mean output activity of the familiar and novel sets over training. Mean output activity across each stimulus set (dark) and individual stimuli (light) shown. **(f)** Normalized mean row magnitude of the modulation term, **W** ⊙ **M**_*t*_ (purple), the unmodulated weight matrix, **W** (green), and total synaptic strength (green and purple) over training. Mean (dark) and individual rows (light) shown. **(g)** Change in important synapse magnitude for familiar and novel inputs as a function of training time (Methods).

The simple network consists of only two populations of neurons, an excitatory input population and an arbitrary output population,^1^ that are sparsely connected by synapses represented by the weight matrix **W** and with non-linearity *ϕ* (*·*) providing the output population activity (Fig. 2b). For brevity, we will refer to this network as the *familiar modulated synapse network* (FMSN). Before training, the synapse strengths are randomly initialized, but they are subject to modulations via an FMS mechanism, represented by the matrix **M**_*t*_. The two modulation types of Eq. (2) and the possibility of strengthening or weakening synapses (i.e. the sign of *η*) gives four qualitatively distinct FMSs. For the example we explicitly consider here, we take the FMS’s modulations to be associative and weakening, meaning a synapse/weight is weakened if both its pre- and postsynaptic neuron are firing, e.g. it is LTD-like (Fig. 2c). This corresponds to updates via Eq. (2a) with *η <* 0. Equivalent plots for the pre-only rule, e.g. STSP-like, and synapses that are strengthened by the modulations, e.g. LTP-like, are provided in the SM (Fig. S1). We will later return to how these choices affect the results presented here.

#### The FMSN develops distinct responses to familiar and novel stimuli

We use a training schedule where the FMSN is sequentially passed stimuli from the familiar training set several times in a random order. That is, at each time step, a stimulus is randomly drawn from the familiar set, noise is added to it, and it is input into the network. After each pass through the network, the FMSs are updated according to Eq. (2a).

For the example considered here, each familiar stimulus is presented to the network 10 times, for a total of 80 training steps. Post-training, we observe that the familiar output activity is significantly suppressed relative to its pre-training activity (Fig. 2d). Comparatively, the novel output activity changes little from the modulations, and so post-training its activity is large relative to the familiar set.^2^ We can understand how the network’s response changes during training by comparing the output activity of the familiar and novel sets had we stopped training after a certain number of familiar stimulus exposures. Over the course of training, we see the network’s response to all 8 familiar stimuli quickly weakens while its response to the 8 stimuli of the novel set remains relatively unchanged (Fig. 2e). This happens concurrently with a growth in the size of the synaptic modulations and, since the modulations in this example are weakening, a smaller total synaptic magnitude (Fig. 2f). Eventually, the changes to the network stabilize as additional examples continue to be presented. The reduction of output activity for the familiar stimuli occurs concurrently with a sparser response to the familiar stimuli over time as well as decreased decodability of stimulus identity, consistent with experimental results of familiarization (Methods, Figs. S2[a-c]) [23, 24].

#### Distinct ‘important synapses’ lead to distinct responses

What about the pattern of synapse modulation is causing this significant change in response for stimuli in the familiar set? Although almost all of synapses undergo some modulation during training (a byproduct of the noise added to inputs), only a small percentage are modulated significantly (Fig. S2d). Intuitively, a reason for the distinct output behavior could be that different synapses have large contributions to the output activity for members of the familiar and novel sets, so changing a subset of them only affects certain stimuli (Fig. 2c). For a given stimulus, we define its *important synapses* as those synapses that would be modulated according to Eq. (2a) from passing the stimulus through the network, before any training has occurred (Methods). With this definition, for the setup we consider here, each (nonzero) synapse has an approximately 2.5% chance of being an important synapse for a given stimulus. Prior to training, we can check that the important synapses of distinct stimuli have little overlap: a familiar and novel stimulus share on average only 0.14% of their important synapses. We can then track how the update rule of Eq. (2a) affects the important synapses of the familiar and novel sets differently. The total strength of the important synapses of the familiar set changes drastically, while those of the novel set remain relatively unchanged because of the small overlap of important synapses (Fig. 2e). It is the greater weakening of important synapses associated to the familiar stimuli, often bringing the neurons’ activity below firing thresholds, that leads to their significantly smaller responses relative to the novel stimuli.

The idea of targeted synaptic modulations as a means of encoding familiarity has been known for quite some time, most famously in Hopfield networks [31]. In the SM, we argue the FMSN can be approximately viewed as a feedforward Hopfield network, i.e. the weight modulations that encode the memory of the familiar inputs are on feedforward synapses and not lateral connections. A stimulus forward pass through the FMSN is similar to measuring its energy in the equivalent Hopfield network. Thus, familiar stimuli having a low mean response is similar to them being low-energy states.

#### Synapse modulations change responses to stimuli in the subspace spanned by familiar stimuli

Since we draw the familiar and novel stimuli from the same distribution, between-stimulus correlations are relatively uniform across all stimuli. How would the FMSN respond to a stimulus that is more correlated with a familiar stimulus than the novel stimuli? More generally, one may consider what characteristics of stimuli determine how much they are suppressed by the learned modulations.

In the SM, we argue that the approximate **M** learned over the FMSN training causes any stimulus that lies in the subspace spanned by the familiar set to have a decreased response relative to its pre-training magnitude.^3^ This includes the familiar stimuli themselves but also their linear combinations (Fig. 3a). Furthermore, since any stimulus can be decomposed into parts that lie within and perpendicular to said subspace, the less any stimulus lies within this familiar subspace the less its response will be suppressed by modulations (Figs. 3a, S3a). In other words, the more a stimulus is correlated with the familiar inputs, the more its response will be suppressed in the FMSN. Part of the success of the FMSN we investigate here relies on the fact that the familiar subspace is small relative to the full space of possible stimuli. Stimuli randomly drawn from the distribution that are not exposed to the network, e.g. the novel inputs, are likely to lie approximately perpendicular to this subspace and thus have their response relatively unchanged by training.

**Figure 3:**
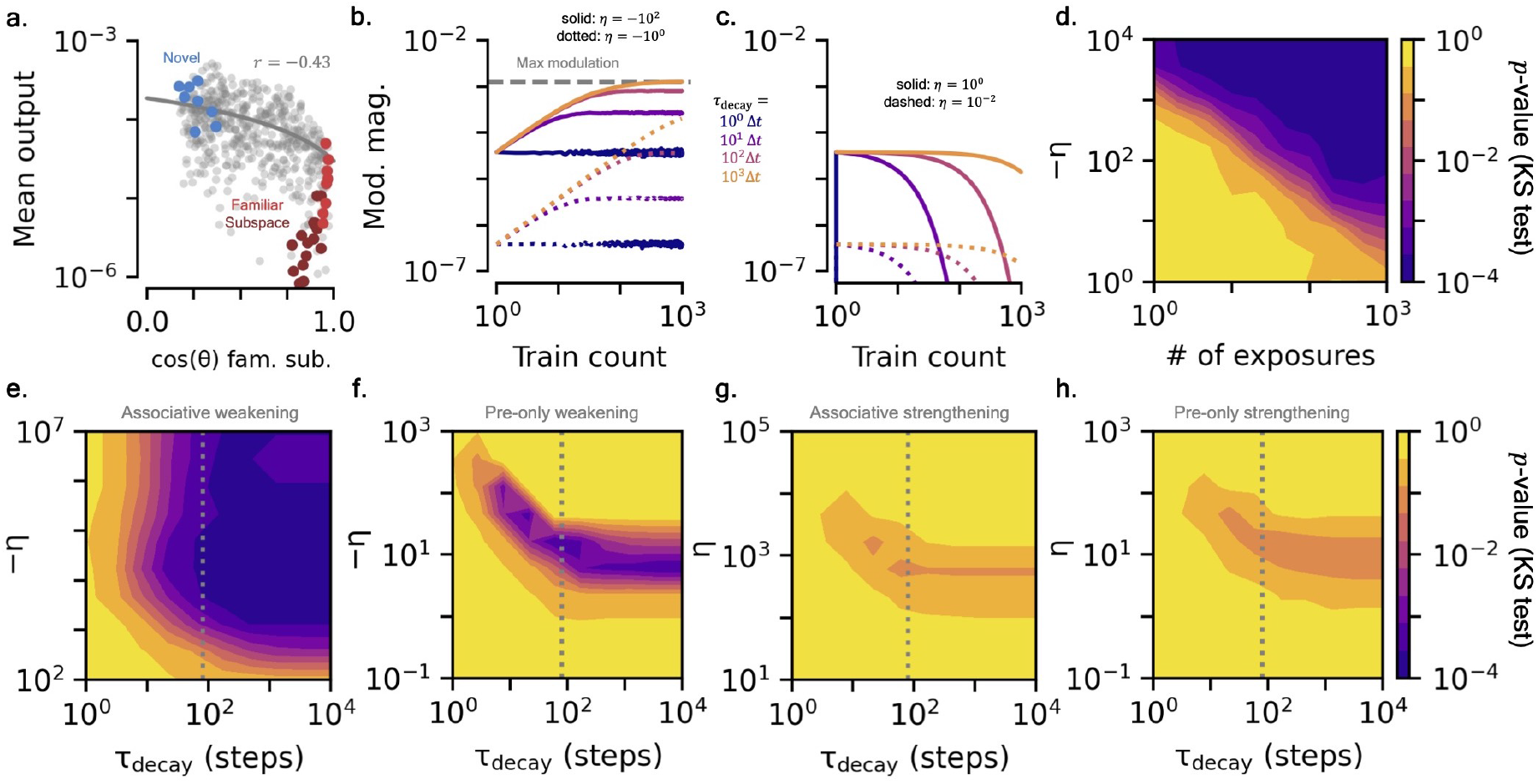
Additional properties of familiarity modulated synapses. **(a)** Cosine distance of stimuli to the subspace spanned by the familiar stimuli (‘familiar subspace’) versus mean output from the FMSN. Grey dots show sparse random binary vectors (Methods). Familiar stimuli (light red), their linear combinations (dark red), and the novel stimuli (blue) are highlighted. Grey line shows linear regression fit. **(b)** Growth of modulation magnitude while being repeatedly exposed to a single familiar stimulus as a function of *τ*_decay_ = 1*/*(1 − *λ*), in units of time steps, and *η*. Dashed grey line shows maximum modulation strength imposed by biological constraints (Methods). **(c)** Decay of modulation magnitude after a single familiar stimulus exposure as function of *λ* and *η*. **(d)** Ability to distinguish output magnitude distributions of familiar and novel sets (KS-test *p*-value) as a function of learning rate, *η*, and the number of times each familiar stimulus has been exposed. **[e-h]** *KS-test to distinguish post-training output magnitude distributions of familiar and novel sets for the four types of modulations as a function of τ*_*decay*_ *and η*. **(e)** FMSN with associative weakening modulations. The grey vertical line shows the timescale of the task, 80 time steps. **(f)** Same as (e), for pre-only weakening modulations. **(g)** Associative strengthening modulations. **(h)** Pre-only strengthening modulations.

#### Learning and decay rates strongly influence magnitude of modulation effects

For training, we have assumed that one stimulus is presented at each time step and time steps are separated by some Δ*t* that could be a characteristic timescale of the input stimulus sequence. Of course, biological effects such as STSP and LTP/D can affect synapses over significantly different timescales. How can the FMSs be adjusted to account for such effects? To investigate this, it is useful to recast the FMS’s decay rate, *λ*, as *decay timescale, τ*_decay_ = Δ*t/*(1− *λ*). Modifying *τ*_decay_ affects the time to saturation of the modulations, allowing one to tune both the number of stimuli and time it takes to see the modulations stabilize as well as their steady-state magnitude (Fig. 3b). Varying the size of the FMS’s other parameter, the learning rate *η*, affects the size of the modulations and thus the speed and magnitude of the FMSN’s change in response. For large enough *η*, the modulations encounter the biological bounds, which limit their growth in size (Fig. 3b). Relatedly, how long a given input influences the modulations, or, how long the FMSN “remembers” a past stimulus, is also affected by the decay timescale and learning rate (Fig. 3c). A single familiar input can influence responses for only a few time steps or thousands, a fact that will play an important role later on when we model novelty effects of significantly different timescales.

The modulation learning rate can also influence how many exposures to the familiar set are needed in order for the network to develop distinct responses relative to the novel set. The larger the modulations, the greater the change to the FMSN from a single input stimulus, leading to distinct responses in a fewer number of stimulus presentations (Fig. 3d). Notably, in the setup we consider here, distinct responses can develop after *just one* exposure to each familiar stimulus. Although large learning rates can lead to quicker response changes, when one has noisy input stimuli, a large learning rate causes the modulations to also fit the noise. Indeed, for fixed training time, there exists optimal learning rates for distinguishing the familiar and novel sets that balance this trade-off between modulations that quickly capture the stimulus signal but not the noise (Fig. 3e).

#### What FMSN properties lead to significant differences in familiar and novel responses?

So far, we have specifically considered the case of an FMS that has associative updates that weaken the network’s excitatory synapses. Of course, this covers a small subset of biological mechanisms – there are synapse modulations that strengthen connections, are only presynaptic dependent, and/or act on inhibitory synapses. The FMSs of Eq. (2) are general enough to model all these cases.

Much of what we discussed above also holds for the presynapse-only update mechanism of Eq. (2b) that also weakens the excitatory synapses of the FMSN (Fig. S1). However, because of its lack of postsynaptic dependence to pinpoint which synapses to update, the pre-only weakening mechanism is much more susceptible to noise. Too large of a learning rate can overfit the noise and quickly cause all inputs to be suppressed (Fig. 3f). Surprisingly, we observe that distinguishing the familiar and novel outputs using modulations that *strengthen* the excitatory connections of the FMSN is significantly less effective for both associative and pre-only dependent FMSs (Figs. 3g,h). Note that the strengthening of excitatory synapses enhances the response of familiar stimuli relative to their pre-training magnitudes (Fig. S1b). We investigate what causes the differences between the strengthening and weakening FMSs in more detail in the SM (Figs. S3[b-j]). In short, we find two major contributions to the relatively poorer performance of the strengthening mechanisms: (1) tighter modulation bounds for strengthening imposed by experiment and (2) neurons’ non-linear behavior that causes firing to cutoff below certain potentials and saturate at higher potentials, built into *ϕ*(*·*). The latter of these effects can be partially overcome by considering an FMS that strengthens *inhibitory* synapses [22]. Stronger inhibition causes the output neurons’ responses to get smaller, a similar effect as the weakening of excitation we found to be the most effective above (SM, Fig. S3[b-j]).

There are many other properties of the FMSN that can be explored that we only briefly touch upon here. For example, allowing modulations to further weaken or strengthen synapses beyond the bounds imposed by associating these modulations with LTP/D and STSP leads to even larger differences between the FMSN’s response to familiar and novel stimuli (Figs. S2e,f). Increasing the noise makes it harder for the FMS mechanism to isolate the signal, making it more difficult to distinguish novel and familiar responses (Fig. S2g). However, effects from noise can be overcome by exposing the network to the familiar stimuli more times, giving it more observations to isolate the signal. Increasing both the number of input and output neurons also increases the distinguishability between the familiar and novel sets (Fig. S2h). Though we leave a full investigation of FMS capacity for future work, we also see the FMSN is capable of becoming familiar with much more than 8 stimuli while still having a distinct response to novel stimuli (Figs. S2k,l). Lastly, we can use the FMSN to predict the most efficient coding of the sparse binary input vectors for distinguishing the familiar and novel sets. Lower sparsity reduces the variance in neuronal responses, and thus makes it easier to distinguish familiar and novel inputs, but also increases the similarity of any two stimuli because each one has more nonzero components. Thus, optimal sparsity is not too high or low (Fig. S2j).

### 2.2 Cortical microcircuit novelty response in a stimulus change task

We now implement the FMSs in a visual cortical circuit model to capture three distinct novelty responses recently observed in mice while they perform an image change detection task [23, 24]. We note that the primary purpose of this model is to demonstrate the flexibility of FMSs and their ability to simultaneously produce three novelty effects observed in the VIP population recordings [23, 24], but do not attempt to constrain this as the *only* types of plasticity that could lead to the experimentally observed results.

#### Review of image change detection task and measurement

The stimuli used in the experimental task consist of a set of 8 familiar training images and a held out set of 8 novel images (Fig. 4a). The task consists of image presentations from these sets at quick, regular intervals that are separated by a grey screen (Fig. 4b). The same image is presented several times in a row before switching to another image within the set and mice are rewarded for responding to the image change by licking a water spout. During this time, neuronal responses from the visual cortex are recorded in hour-long sessions using two-photon calcium imaging. Mice are trained on what becomes a familiar set of eight images and their neuronal responses are recorded in a ‘familiar’ imaging session after achieving a performance threshold (Fig. 4c). Shortly after, neuronal responses are also gathered over multiple sessions when the mice are exposed to the same task using the novel set of eight images. The mice’s initial exposure and exposure after at least one session to this novel set of images are referred to as the ‘novel’ and ‘novel-plus’ imaging sessions. Additionally, only during the imaging sessions, image omissions can occur, i.e. grey screen is displayed in place of a single image presentation (Fig. 4b).

**Figure 4:**
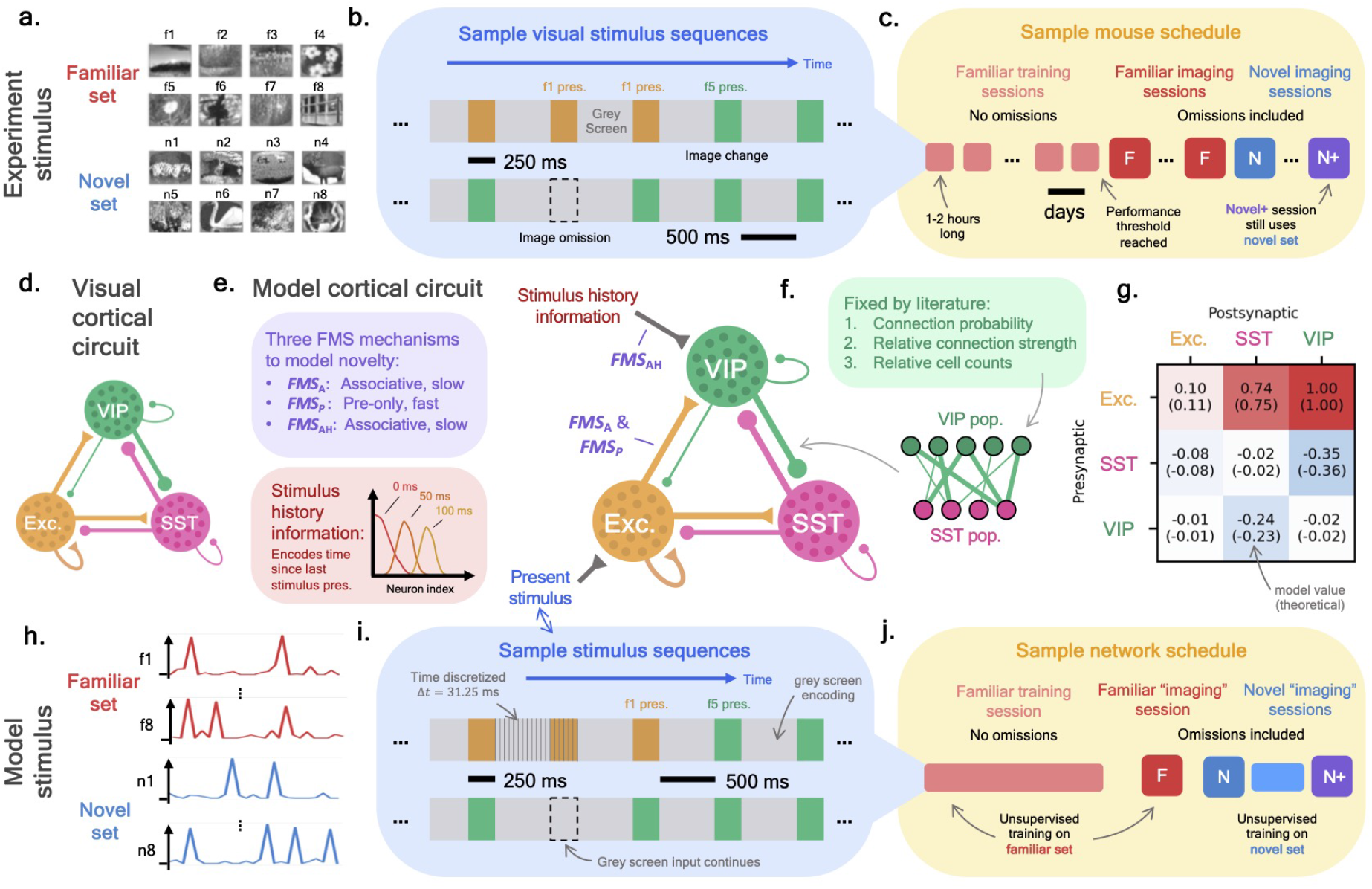
Image change task and visual cortical circuit: experiment and model setup. **[a-c]** *Experimental stimulus details*. **(a)** Example familiar and novel image sets (reproduced, with permissions from Ref. [24]). **(b)** Sample stimulus sequences showing an image change (top) and omission (bottom). **(c)** Typical training/imaging schedule. Boxes represent sessions that occur on different days, each lasting an hour or two. **(d)** Diagram of a subset of the visual cortical circuit showing the cell populations that were recorded in experiment. **(e)** Diagram of the cortical circuit model we study in this work. The SST, VIP, and Exc. circles each represent populations of neurons connected by weights fixed from experimental data [17, 18, 20]. Three FMS mechanisms (purple); *FMS*_A_, *FMS*_P_, and *FMS*_AH_; are added to the network to model novelty responses. At each time step, the network receives inputs representing an encoding of the ‘present stimulus’ being shown (blue) as well as ‘stimulus history’ information (red) in the form of an encoding of the time since the last image presentation (Methods). **(f)** Many features of the cortical circuit model are fixed by experimental literature [17, 18, 20]. **(g)** Mean inter-population connection strengths. Values from an exemplar model (top) and analytically computed values (bottom) are shown (Methods). **[h-j]** *Model stimulus details*. **(h)** Exemplar familiar and novel stimuli sets, drawn from a sparse random binary vector distribution. **(i)** Sample present stimulus sequences showing a stimulus change (top) and omission (bottom, Methods). **(j)** Model training scheduling consisting of training session on familiar stimuli, familiar ‘imaging’ session, and then novel/novel-plus ‘imaging’ sessions. At all points of training and imaging, all FMSs are continuously updated via their unsupervised rules as stimuli are passed through the network.

The response to various novelty effects are recorded across several transgenic lines to capture excitatory, SST, and VIP population responses in the visual cortex. These cell populations form the cortical microcircuit discussed in the introduction whose connection probabilities and strengths have been carefully studied (Fig. 4d). Experimental analyses show that the effects of novelty give rise to significantly different responses in these three populations [23, 24], which we discuss in more detail below.

#### Cortical microcircuit model

Given that we have observed the FMS mechanism yields distinct responses to familiar and novel stimuli, we built a model of the cortical microcircuit to study if it can develop the several experimentally observed novelty responses when exposed to stimulus sequences similar to that of experiment. Our firing-rate model consists of three groups of neurons, representing the SST, VIP, and excitatory neuron populations (Fig. 4e).^4^ The excitatory population receives inputs representing the bottom-up encoding of the raw stimulus sequence while the VIP population receives inputs representing top-down information about the history of the sequence (specifically the ‘timing’ of recent stimuli and not image identity, see below for additional details). We estimate connection properties between populations by aggregating results from several recent experimental studies. Relative cell counts are estimated from in situ hybridization experiments [17, 18] (Methods). An estimation of inter-population connection probabilities comes from multi-patch synaptic physiology [20] and the relative strength of neuron connections between populations is estimated from the same study, supplemented with additional cell dynamical properties from electrophysiology recordings [19] (Figs. 4f, g; Methods). In particular, fits of measured postsynaptic potentials are used to estimate unmodulated, individual synapse strengths as a function of the pre- and postsynaptic cell type (Fig. S4). Due to the unprecedented detail of recent experiments [17–20], coupled with necessary corrections from the experimental to the in-vivo setting, we believe the ‘skeleton’ of the cortical circuit model represents one of the most accurate estimates of this system to date.

We allow the connections in our microcircuit model to change by introducing several FMS mechanisms into the synapses connecting the various populations of the network. Since it is observed that the VIP cells drastically change their response across all three types of novelty in the experiment [24], in this work we focus on adding FMS mechanisms to capture their specific novelty responses. The purpose of focusing only on the VIP response is to demonstrate how several FMS mechanisms may collectively model distinct novelty responses within a single population. We leave a complete modelling of the distinct cell type responses and related plasticity mechanisms for future work. To capture the VIP novelty responses, we add three separate FMS mechanisms to the synapses onto the VIP neurons: *FMS*_A_, *FMS*_P_, and *FMS*_AH_ (Fig. 4e).

1. ***FMS***_**A**_ **(Associative, Exc**. → **VIP)** is added on the synapses going from the excitatory to the VIP cells. Its learning and decay rate (*η* and *λ*) are tuned to learn and retain familiarity over a timescale of hours to days. Since it operates on a slow timescale and is pre- and postsynaptic dependent, *FMS*_A_ could model LTD-like effects on said synapses.
2. ***FMS***_**P**_ **(Pre-only, Exc**. → **VIP)** is *also* added to the set of synapses between the excitatory and VIP populations, but unlike *FMS*_A_ it is tuned to learn and forget on a timescale of seconds. The fast timescale over which it operates and its presynaptic dependence makes *FMS*_P_ a natural model for STSP-like effects on the synapses.
3. ***FMS***_**AH**_ **(Associative, Stimulus history** → **VIP)** is added to the synapses feeding into the VIP population from the stimulus history input neurons (see below). Its learning and decay rate are tuned to operate over long timescales, similar to the LTD-like *FMS*_A_.

Motivations for adding these particular modulations within the circuit are discussed below.

#### Model ‘image’ change stimulus

As we saw in the FMSN, modulations are entirely driven by the stimuli being passed to the network, so we reproduce the pattern of stimuli from the image change detection experiment. To represent neuronal encodings of the images used in the experiment, we again use random sparse binary vectors as the distinct stimuli (Fig. 4h). An ‘image’ presentation is represented by a stimulus encoding being passed to the network for several time steps along with time-varying noise (Fig. 4i). The image presentation is followed by a proportional number of grey screen time steps, where the network receives only noisy input (Fig. 4i). This pattern repeats with a similar distribution of image change times used in experiment (Fig. S5a). Stimulus omissions are represented by additional grey screen time steps (Fig. S5b). We assume the excitatory population receives this bottom-up *present stimulus* input and drives the other populations (Fig. 4e). Additionally, we assume the microcircuit receives top-down inputs representing information about the recent history of the stimulus (Methods, Fig. S5c). In particular, the *stimulus history* input is an encoding of the time since the last stimulus presentation, with encodings of similar times more correlated than disparate times.^5^ This information is passed directly to the VIP cells, which are known to receive feedback inputs from higher cortical areas [8, 20]. Finally, time-correlated noise is injected into all neuron populations to represent activity from sources neglected in this model, e.g. activity from behavior (Methods, Fig. S5d).

Similar to the training schedule used in experiment, we first expose the network to the familiar stimuli over a long training session, then gather cell responses to the task using the familiar set in what we continue to call an ‘imaging’ session. Immediately afterwards, we gather responses to the stimulus change task using the novel stimulus set, and, after additional exposure to the novel image set, gather the novel-plus responses (Fig. 4j, Methods). The familiar, novel, and novel-plus imaging session stimulus sequences are statistically identical. Importantly, the neuronal response presented here are gathered in a continuous learning setting, i.e. the network continues to modulate its weights via *FMS*_A_, *FMS*_P_, and *FMS*_AH_ at all steps of training and imaging. We scan over three parameters, the learning rates for all three FMS mechanisms, to determine modulation rates that best match experimental observations (Methods, Fig. S5o). We emphasize that, other than minor adjustments to the network at initialization to ensure realistic responses, the cortical circuit model only undergoes unsupervised adjustments via the various FMS mechanisms from exposure to stimulus sequences that closely match the stimuli on which the mice were trained (Methods).

For the purposes of comparing our model to experiment, we first focus on three distinct novelty responses that our model captures seen in mean VIP population responses of the experimental data [24]: (1) absolute, (2) contextual, and (3) omission novelty (see Fig. S6 for SST and Exc.).

##### 1. Absolute novelty: familiar modulation occurs despite irregular stimulus sequence

The change between the familiar and novel image sets represents *absolute novelty* – up until the novel imaging session the mice have never observed the set of images now used in the image change task. In both experiment and our model, the VIP cells respond weakly to image presentations in the session that uses the familiar set relative to image presentations in the session that uses the novel set (Fig. 5a). As we confirm below, for the cortical circuit model, the change in response is caused by *FMS*_A_, the slow-learning FMS mechanism on the excitatory to VIP synapses. *FMS*_A_ functions almost identically to the FMSN discussed earlier: over training, exposure to the familiar stimuli causes the network to develop a suppressed response to them relative to the novel stimuli (Fig. 5b). The stimulus sequence here is quite different from that of the FMSN: a single stimulus is repeatedly input to the network and is often separated by noisy grey screen. Additionally, the postsynaptic population of *FMS*_A_, the VIP cells, receive input from several additional sources such as the SST population and the recurrent VIP connections. Nevertheless, over the long familiar training period, *FMS*_A_ gradually modulates the important synapses of the familiar set more than the novel set, leading to a distinct response across sessions (Fig. 5b). Notably, the additional synaptic inputs and noise make modulating only those synapses important for the familiar set more difficult, leading to a fair amount of suppression to novel inputs as well. However, after these changes stabilize, we still observe distinct responses to the familiar and novel stimuli. Just like the FMSN, the change in stimulus response occurs concurrently with a growth in the modulations of *FMS*_A_ over training and, since the modulation are once again weakening, an overall decrease in the strength of the synapses connecting the excitatory population to the VIP population (Fig. 5c).

**Figure 5:**
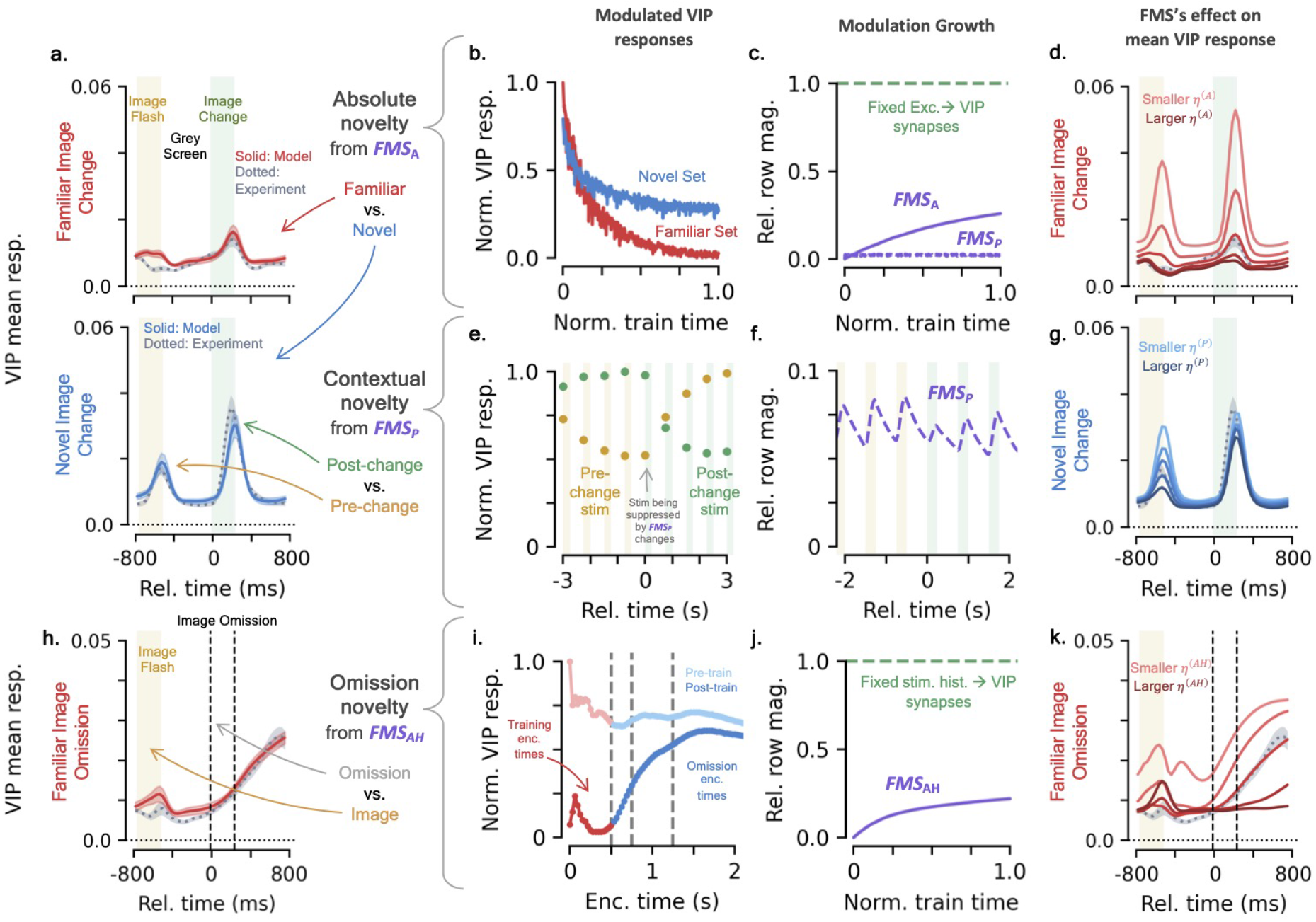
FMSs implement three distinct novelty effects in a cortical circuit model. **(a)** Mean VIP responses to image changes of the cortical circuit model (solid colored line) and experiment (grey dotted line) [24]. Top shows mean response in the familiar imaging session and bottom in the novel imaging session. Green-shaded background represents time where changed image is presented, yellow-shaded is pre-change image. **[b-d]** *Absolute novelty and FMS*_A_. **(b)** Exemplar mean VIP image change responses to the familiar (red) and novel (blue) sets over training (Methods). **(c)** Change in *FMS*_A_ (purple solid) and *FMS*_P_ (purple dashed) modulations over training. Green dotted line shows fixed portion of excitatory to VIP synapses. **(d)** Same as top of (a), for different *FMS*_A_ learning rates (Methods). **[e-g]** *Contextual novelty and FMS*_P_. **(e)** Normalized VIP responses around an image change event. Yellow is pre-change image, green is changed image. **(f)** Same as (c), but zoomed in to show change in *FMS*_P_ over an example image change. **(g)** Same as bottom of (a), for different *FMS*_P_ learning rates. **(h)** Same as (a), for mean VIP response to image omission in a familiar session. Area between vertical dotted lines represents times where image would normally be presented. **[i-k]** *Omission novelty and FMS*_AH_. **(i)** Change in VIP response to various encoded times over training. Red encoded times are seen during training, blue are times when omissions are present. **(j)** Change in *FMS*_AH_ (purple) modulations over training. Green dotted line shows fixed portion of stimulus history to VIP synapses. **(k)** Same as (h), for different *FMS*_AH_ learning rates.

To confirm that it is the modulations from *FMS*_A_ that cause the large difference in VIP response across the familiar and novel sessions, we can isolate its effect by training identical microcircuit models with different *FMS*_A_ learning rates. Indeed we see that, as we decrease (or increase) *FMS*_A_’s learning rate and thus its modulation magnitude, the response of the VIP population in the familiar sessions grows (or shrinks) as the overall strength of the excitatory to VIP synapses changes (Fig. 5d, S7a). Once again there is a trade off between modulations with too large of a learning rate that suppresses all responses and those with too small of a learning rate that suppresses none.

##### 2. Contextual novelty: fast familiarization and forgetting captures local oddball effects

In the experimental task, image changes represent *contextual novelty* – since images are repeatedly presented at least 10 times, when the image identity changes it represents a local oddball and is contextually novel. In the novel session, we observe an increased response of VIP cells to image changes relative to the pre-change image in both our model and the experimental data (Fig. 5a, bottom). Although smaller, the effect is also present in the familiar session (Fig. 5a, top). Notably, this is a very different effect than the absolute novelty we discussed above; it only takes seconds for mice to establish an image as their baseline and this information is quickly updated to the current image being presented [24]. To model a novelty effect that learns and forgets quickly, *FMS*_P_ is introduced. For *FMS*_P_, the presynaptic-dependent modulations make the most recent stimulus presentations become familiar, leading to smaller VIP response on repeats of the same stimulus. An image after a change is novel to *FMS*_P_, meaning the VIP response is larger because it is not familiarity suppressed (Fig. 5e). After the change occurs, *FMS*_P_ begins suppressing the important synapses of the current stimulus, while those that were important for the pre-change image are gradually released from suppression. This rapid turnover is reflected in the significantly quicker growth and decay of the *FMS*_P_ modulations relative to those of *FMS*_A_ (Fig. 5f). Finally, as we did for *FMS*_A_, we see that varying the learning rate of *FMS*_P_ isolates its effects on the cortical circuit response. A weaker learning rate changes the relative heights of the pre-change and post-change VIP responses because the image that has been presented several times in a row is less suppressed by modulations (Fig. 5g, S7b).

The operation of both *FMS*_A_ and *FMS*_P_ on the excitatory to VIP synapses demonstrates a remarkable property of the FMSs: distinct types of modulations, e.g. slow and fast, can function on the *same* set of synapses simultaneously. This matches biology, where effects of LTP/D and STSP can affect the same set of synapses. The distinct FMSs can encode different types of novelty present in stimuli, allowing them to model the various novelty responses that are observed in certain neuronal populations. This allows for a single synaptic population to affect the postsynaptic population’s response in a way that compounds the various novelty effects. For example, the largest responses of the VIP cells occur when both absolute and contextual novelty occurs, i.e. a novel image change, which is a result of the minimal suppression from *FMS*_A_ and *FMS*_P_ simultaneously.

##### 3. Omission novelty: a decrease in familiar correlation causes omission ramping

Due to the temporal structure of the task during training, images are expected to be separated by 500 ms of grey screen. *Omission novelty* occurs when, instead of an image, additional grey screen is displayed, representing a global oddball. We observe a ramping response in the VIP cells of both the model and experiment when images are omitted (Fig. 5h). In the model, this is a result of *FMS*_AH_, the FMS mechanism on the stimulus history input synapses to the VIP cells. Recall that the stimulus history signal is a neuronal encoding of the time since the last stimulus presentation, where the encodings of similar times are more correlated than disparate times (Fig. 5e, Methods). Over training, *FMS*_AH_ becomes familiar with encoded times-since-last-image that occur with no omissions, suppressing the corresponding VIP responses to the stimulus history signal. When omissions occur, longer-time encodings are passed to the network and the familiarity suppression is lost, leading to an increased response in VIP cells. The ramping occurs because the longer-time encodings have a less correlated representation to the familiar short-time encodings. That is, similar to what was seen in the FMSN, the network has formed a familiar subspace of the short-time encodings and the longer-time encodings that gradually get further from this subspace cause a gradual increase in the VIP response (Fig. 5i). The encoded-times that are novel but still close to the encoded times that are familiar, e.g. 510 ms, have their outputs quite suppressed. The longer encoded-times, e.g. 1000 ms, have outputs barely suppressed at all. As with the other FMSs we’ve investigated, this change in response occurs concurrently with a gradual growth in the modulations on *FMS*_AH_’s synapses over training (Fig. 5j). Additionally, the size and time of onset of the VIP ramping can be changed by adjusting the magnitude of *FMS*_AH_’s learning rate (Fig. 5k, S7c).

The omission novelty responses occurring concurrently with the absolute and contextual demonstrates the ability to have multiple inputs into the same postsynaptic cell population with distinct synaptic dynamics. *FMS*_A_ and *FMS*_P_ operate on the excitatory to VIP synapses, while *FMS*_AH_ acts on the stimulus history to VIP synapses and all three can produce their corresponding novelty effect in the VIP cells when the corresponding stimulus occurs.

###### Novel images become familiar with exposure over time

Although the novel image set is initially unfamiliar to the mice and evokes distinct novelty-related responses across cell populations, over many exposures one would expect the images to gradually become familiar to the mice. Indeed, the enhanced VIP response to the novel images persists throughout the entire novel imaging session, but gradually disappears as the mice become accustom to the novel set over many sessions of exposure [23, 24]. Since our model is evaluated in a continuous learning setting, the FMS mechanisms are actively modulating the network’s response, allowing it to also adapt to the novel stimulus set over time in the same way it adapted to the familiar set during training. Hence, our model also exhibits a gradual change in response to the novel image set over sessions, eventually returning to a suppressed VIP response to novel set images in the novel-plus imaging session (Fig. S6a, top right; Methods). In the experiment, even after being exposed to the novel set of images, the familiar set of images still evoke a response consistent with them being familiar stimuli [24]. The *FMS*_A_ modulations decay slowly enough that the modulatory effects of both the familiar and novel images can persist simultaneously (Fig. S5p). Additionally, as in experiment, the image omission response does not change considerably between the familiar and novel-plus sessions (Fig. S6b, top right).^6^

###### Cortical circuit model’s cell subpopulations have diverse coding

Although the changes in mean response of our model’s cell populations is dependent upon the behavior of individual cells within these populations, we observe a significant variation in each neuron’s response over both stimulus features and experience-level. Indeed, a key finding of Ref. [24] is the emergence of functional cell subpopulations within the VIP, SST, and excitatory cell populations. Within each population, subpopulations are identified with similar changes in coding features over experience-level, as measured by a coding score metric that we briefly review. To determine the coding score, each cell’s response is fit using a kernel regression model [57–60]. Several input features that may influence a cell’s response, including image presentations, image omissions, and task-relevant triggers such as image changes, are convolved with fit kernels to reproduce the cell’s activity (Fig. 6a, top; Methods). To determine the importance of a given input feature category in explaining a cell’s activity, the regression model is refit while omitting each category and its corresponding kernel(s). The *coding score* of a cell to a given feature category is defined as the relative amount of variance explained that the regression model loses by removing the feature category (Fig. 6a, bottom; Methods). This procedure is repeated across the 3 distinct sessions/experience-levels for 4 feature categories, resulting in a 12-dimensional coding-score vector for each cell. Lastly, the resulting coding score vectors of a given cell type are collected across mice and run through an unsupervised clustering algorithm (Fig. 6b, Methods) [24].

**Figure 6:**
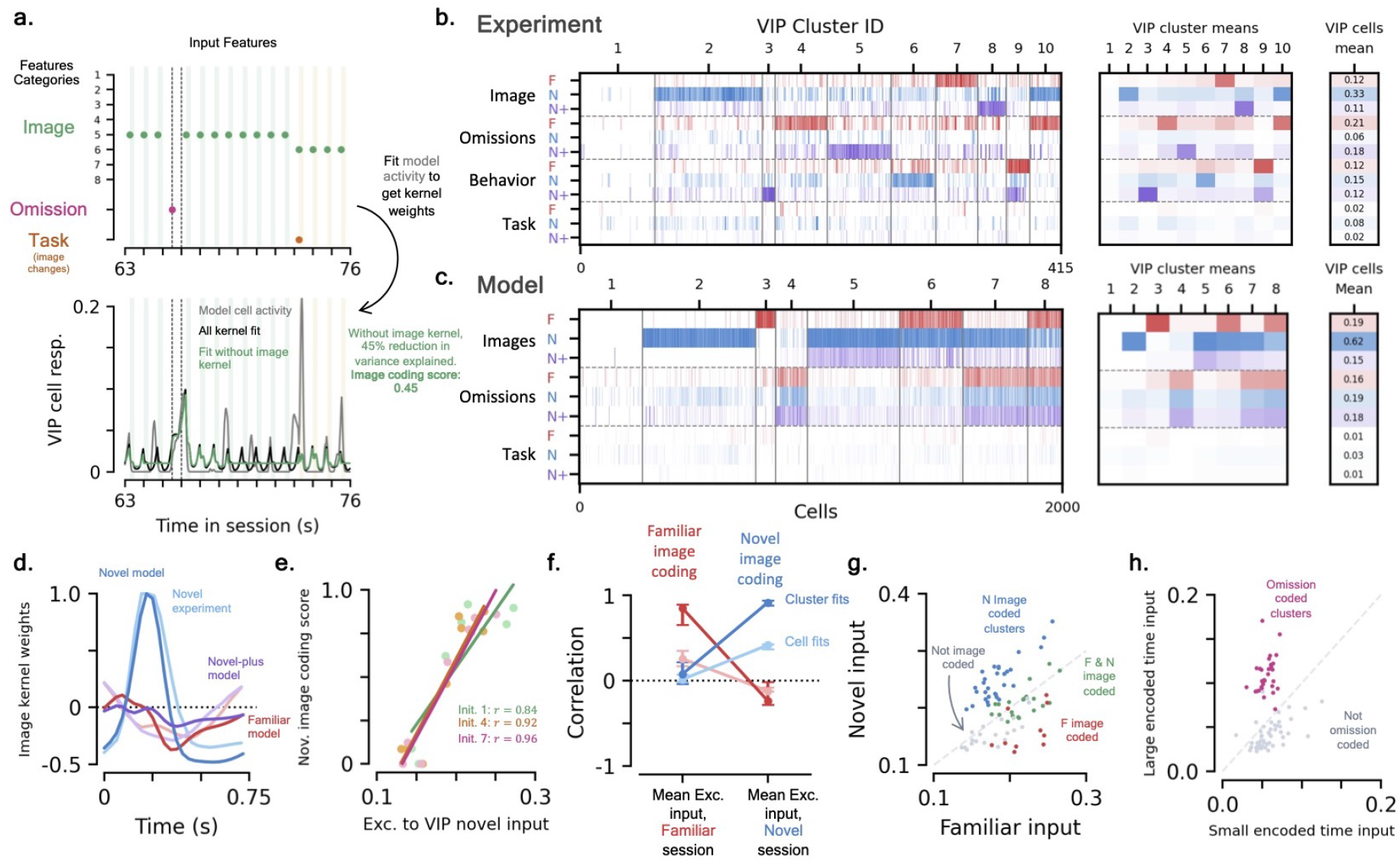
Coding diversity in the cortical circuit model. **(a)** Kernel regression model: input features are convolved with learned kernels and summed to predict individual cell activity in a kernel regression model (Methods). Top: the cortical circuit model is driven by three primary feature categories: image presentations, omissions, and task-relevant image changes. The experimental data shares these same feature categories as well as behavioral effects [24]. Bottom: feature categories are removed one at a time and the kernel regression model is refit to determine each feature’s contribution to a cell’s activity, summarized by the category’s coding score. **[b, c]** *VIP coding scores*. **(b)** Experimental results [24]. Left shows clustered VIP coding scores for all four feature categories across the familiar, novel, and novel-plus sessions; middle shows cluster means; and right shows mean over all cells (Methods). **(c)** Same as (b) for the cortical circuit model. **(d)** Normalized mean image feature kernels for the familiar (red), novel (blue), and novel-plus sessions (purple). Dark lines show fits to model data, light lines to experimental data [24]. **[e-h]** *Cell-specific cortical circuit properties influences VIP cell coding*. **(e)** Cluster-averaged novel image coding scores as a function of the mean input each cell receives from the excitatory population during the novel session. Colors show three different initializations, dots are cluster-averaged values, lines are linear regression fits. **(f)** Change in correlation of familiar (red) and novel (blue) image coding scores with network properties. Correlation over all VIP cells (light) and cluster-averaged values (dark). Dots are median, error bars are Q1 to Q3 over initializations. **(g)** Cluster-averaged network properties across 10 different initializations. Colored dots (red, blue, green) show clusters with distinct types of image coding, grey dots are clusters not coded to images. Grey dotted line shows equal familiar and novel input. **(h)** Network properties for omission-coded clusters (pink) versus omission-agnostic clusters (grey).

We use the same analysis pipeline to analyze cell subpopulation diversity in our cortical circuit model. We again focus on investigating the VIP population’s activity in particular (Fig. S8 for Exc. and SST). Since we have no explicit behavioral effects in the network, we only code for three input feature categories: image presentations, omissions, and changes (‘task’). Repeating the aforementioned fitting procedure to determine coding scores and then clustering the data across 10 network initializations, we again observe diverse coding across features and experience-levels in the VIP population (Fig. 6c). Notably, the resulting kernel fits of the cortical microcircuit qualitatively resemble the fits on the experimental data (Fig. 6d, S8). Membership of the clusters are shared across the different networks, demonstrating the diversity is not due to the different initialization parameters or training sequences (Fig. S8f).

Many of the same subpopulation motifs observed in experiment are also present in the microcircuit model [24]. Although our overall cell coding scores are larger, we observe subpopulations that have very little coding to any feature or can be coded to one or more features (see Sec. 4.4.3). Several clusters of cells have weak coding to images in the familiar session only to gain said coding in later sessions and vice versa. As in experiment, since the VIP population responds more strongly to image changes in the novel session, its average image coding score increases relative to that in the familiar and novel-plus sessions (Fig. S8c, right). Nevertheless, there are many features of the experimental VIP subpopulations that we do not observe in our model: solely novel-plus image coded clusters, a clear over representation of novel image coding, and significantly less diversity in omission coding.

The FMS mechanisms we have inserted into our model evidently affect cell diversity in addition to the mean responses. Since the familiar and novel stimulus trains are statistically identical, without the FMS modulations the coding scores would be distributed across the two sessions evenly. That is, the fact that the excitatory to VIP synapses are not as familiarity-suppressed by *FMS*_A_ causes many cells to become image-driven in the novel session. The effects of the FMS modulations can also be seen in the substantial difference in coding scores between the novel and novel-plus sessions. Since these sessions are driven by the same stimuli trains, there should be no statistical difference in coding scores without changes in connection properties due to modulations. There are several qualitative features of the experimental data we do not capture that are outside the scope of this work, e.g. clusters with experience-dependent omission coding. In the Discussion, we comment on how additional FMS mechanisms could be added to the model to produce such effects.

###### Cell-specific synapse properties strongly influence cell coding

Having observed a diversity in VIP cell coding in the cortical circuit model, we analyzed what cell-specific network properties may be responsible for the heterogeneous coding. The point-like neurons in our model primarily differ in their connectivity properties and how said connectivity is acted upon by the FMS mechanisms. To determine what differences individual VIP neurons may have that explains their diverse coding scores, we take many cell-specific network properties and see how well each of these correlates with either individual cell coding scores or cluster-averaged coding scores (Methods). For example, if we plot the cluster-averaged novel image coding scores as a function of the total synaptic input a VIP cell receives from the excitatory to VIP synapses during the image presentations of a novel session, we observe a statistically significant positive correlation (*p* = 0.002; Fig. 6e). Since the amount of input a given VIP cell receives is influenced by the heterogeneous synaptic connections and image encoding signals, this quantity is different for each VIP cell and is evidently reflected in the cell’s image coding scores. We repeat this procedure across 16 cell-specific network properties of our cortical circuit model and all 9 coding score values (Fig. S9). Additionally, we analyze how said network properties vary as a function of each VIP cluster (Fig. S10).

Across all the cell-specific properties we consider, the strength of the modulated excitatory to VIP synapses mentioned above has the largest correlation with the novel image coding scores (Figs. 6f, S9). Similarly, if a given VIP cell happens to have strong input from excitatory synapses during familiar image presentations, it tends to have a larger familiar image coding score (Figs. 6f). As might be expected, we see the amount of input a cell receives in familiar session has relatively little correlation with the novel image coding score and vice versa. These coding score correlations can be used to determine properties that cells in a given cluster may share. For example, the cluster-averaged excitatory input from the familiar and novel sets roughly separates clusters into those that are familiar and/or novel image coded across network initializations (Fig. 6g).

Interestingly, the image coding scores for the familiar and novel-plus sessions also have a strong correlation with how much stimulus history input the cells receive (Fig. S9). That is, in the absence of significant within-layer excitatory input, a result of *FMS*_A_’s suppression, the VIP cells that happen to have strong synapses from stimulus history sources are the most coded to images during the familiar and novel-plus sessions. An important distinction about the stimulus history input the VIP cells receive is that it is image-presentation-correlated but is uniform across all images, up to effects from noise. This lines up with the experimental observation that VIP cells increase image decodability during the novel session, but do not see a significant increase in decodability during presentations of the familiar and novel-plus sessions [24]. Note that since these inputs do not change across images or sessions significantly, this could alternatively be interpreted as cells coded to familiar/novel-plus images are those closer to firing thresholds during all image presentations.

Although the model’s omission coding exhibits far less experience-level diversity than what is observed in the data, we see how strongly connected a given VIP cell is to those cells excited during omission-encoded times, i.e. *>* 500 ms, strongly influences the omission coding (Fig. S9). Additionally, there is a slight negative correlation of the omission coding scores with the amount of input a VIP cell receives during non-omission encoded times, i.e. ≤ 500 ms. Any VIP cell that receives stimulus history input during the training session will have their presynaptic connections suppressed by *FMS*_AH_, meaning said inputs will not influence the omission response. We see the clusters that are omission-coded tend to have larger input during large encoded times compared to those that do not (Fig. 6h).

## 3 Discussion

In this work we have introduced familiarity modulated synapses (FMSs), a simple familiarity detection mechanism that relies solely on local, unsupervised synaptic modulations to encode exposure to past stimuli. The individual modulations of the FMS mechanisms evolve via well characterized dynamics: Hebbian or anti-Hebbian associative or presynaptic only dependence. We first investigated the basic properties of the FMS mechanism in a simple feedforward network, what we refer to as the FMSN. There we saw that, unlike several other familiarity-detection models, FMSs can detect novelty in a single forward pass, which is supported by evidence showing such stimulus distinctions can occurs rapidly in humans [61, 62]. We then demonstrated the generalizability of FMSs by modeling three distinct novelty novelty effects recently observed in a cortical disinhibitory circuit containing excitatory, VIP, and SST neurons. The connectivity of the cortical circuit model we develop is constrained by an aggregate of recent experimental results [17, 18, 20]. The three separate VIP novelty effects were reproduced in a continual learning setting with experimentally realistic stimulus sequences. Finally, due largely to the modulations that change the network’s response over time, we found significant cell subpopulation diversity in our model, reproducing results that have been recently highlighted in the cortical disinhibitory circuit [24].

In the cortical circuit model, we specifically focused on reproducing three distinct novelty effects seen in the VIP population responses. The goal of this modelling study was to demonstrate how several FMS mechanisms could be combined to produce various novelty responses within a single cell population. With this in mind, the specific choices of where we have added FMS mechanisms and their parameterization are not tested against other potential plasticity configurations that could also give rise to the novelty responses. With the addition of plasticity elsewhere in the cortical circuit and/or further neuronal contributions to the model that we have neglected, plasticity that weakens the synapses feeding into the VIP neurons may not be necessary to produce novelty effects [22]. We leave an extensive study of how various plasticity configurations could give rise to *all* the novelty effects observed within the cortical circuit for future work (see below) [24]. Although we do not explicitly model all the novelty effects observed in experiment, given the results we have seen here we can speculate on how the generalizability of FMSs would allow them to model such effects. The inhibition from the VIP population alone is not enough to produce the change in SST response seen across the familiar and novel sessions (Fig. S6). A slow strengthening FMS on the the excitatory to SST synapses, i.e. similar *FMS*_A_ with positive learning rate, would result in a larger SST population response in the familiar session, similar to what is observed in the experimental data [23, 24]. A fast FMS, analogous to *FMS*_P_, on synapses that carry the bottom-up signal into the excitatory population could drive the observed increased excitatory response to image changes, very similar to what we produced in the VIP cells. We also do not attempt to model the suppressed omission ramping that is observed in the VIP population during the novel session [24]. A population of VIP neurons that have increased mean activity in a novel session, via an FMS mechanism similar to *FMS*_A_, could act to gate the ramping signal. If this population inhibits the excitatory neurons that produce the stimulus history input signal it would lead to an overall smaller input and thus a smaller ramping only during the novel session. Furthermore, if this VIP population activity was different across experience-levels, like the VIP population in our model and experiment, the inhibition to the history input would change between sessions, potentially leading to experience-level-dependent omission coding that has been observed [24] (Fig. 6b). We also note that it may be possible to observe the omission ramping response using a fast, STSP-like FMS in place of the LTD-like *FMS*_AH_, i.e. the ≤ 500 ms encoded times become familiar on a timescale of seconds rather than hours/days.

We cannot rule out that part of the novelty responses observed in the cortical circuit may be driven by signals outside the visual cortex (though see Ref. [52]). Regardless, since it has been confirmed that it is not the specific stimuli in the familiar and novel image sets that evoke the novelty responses [24], they must be generated somewhere in the brain and we have demonstrated a plausible plasticity mechanism that could produce said responses in cell populations. We do not attempt to model the heterogeneous learning rates across the same synaptic population that has been observed in short-term plasticity [20]. However, since synapses continue to either be strengthening or weakening on average, we do not believe this will affect the mean population activity significantly. We also neglect effects of neuron scaling, e.g. adjusting those synapses not strengthened/weakened via, say, heterosynaptic LTP or LTD. The FMS’s similarity to Hopfield networks means modern extensions to said networks could be used in the FMSs to increase their effectiveness and/or capacity [63].

As mentioned above, the FMSs’ simplicity, generality, and effectiveness in producing novelty effects makes them an ideal candidate for studying plasticity in future work. For example, modelling projects could scan over various FMS configurations and parameterizations within the cortical circuit to match the experimentally observed novelty responses and see if the resulting fits match or may constrain experimentally observed plasticity [20, 39]. On the experimental end, understanding how neuronal responses change throughout all of training would allow us to further characterize the types of plasticity that modulate experience-dependent activity. From our cortical circuit model, predictions about how connectivity and plasticity influence the observed cell subpopulation diversity could be tested by pairing physiological and learning studies together. Altogether, the effectiveness of FMSs highlights the role simple modulations within large synapse populations may have in shaping neuronal responses to stimuli. We demonstrated the FMS relies on no specialized training and testing schedules and requires no carefully placed excitatory or inhibitory connections to operate. Our cortical circuit demonstrates two important features of the FMS mechanism: (1) its ability to operate with several distinct types of modulations on the *same* synapses, even at significantly different timescales, as well as (2) the ability to have multiple inputs with distinct synaptic dynamics influencing a single cell population. Crucially, these mechanisms allowed us to model the novelty effects that have been observed to occur over significantly different timescales (seconds to days) and from different sets of information on the same set of cells [24]. Other than a few parameters adjusted at initialization to ensure realistic input and firing rates (*<* 10), the cortical circuit’s response is driven by the FMS mechanisms, themselves only containing 1 free parameter a piece: their learning rate/size of modulations. It is surprising to the authors that what seem like complicated novelty-responses can be captured by such straightforward modulation mechanisms, yet speaks towards the influence that simple synaptic changes can have on our brains.

## 4 Methods

Many quantities we consider throughout this work are sequence time-dependent, and their time dependence is generally denoted by a subscript *t* (and sometimes *s*), i.e. **x**_*t*_. Time is uniformly discretized in our setup, so the quantities **x**_*t*_ and **x**_*t*+1_ are separated by Δ*t*. When unambiguous, we use *I, J* = 1, …, *d* to index the neurons of the presynaptic layer, and *i, j* = 1, …, *n* to index the neurons of the postsynaptic layer.

### Data availability

Supporting code for this paper can be found at: https://github.com/kaitken17/fms. Data from the two-photon image change detection task can be found at: https://portal.brain-map.org/explore/circuits/visual-behavior-2p.

### 4.1 Familiarity Modulated Synapses and Networks

In this section we give additional details about the FMSs and a brief introduction to the networks they’re used in. Additional details about the FMSN and the cortical microcircuit model can be found in Secs. 4.3 and 4.4, respectively.

#### 4.1.1 Familiarity modulated synapses (FMSs)

As in the main text, we take **x**_*t*_ ∈ ℝ^*d*^ to represent the presynaptic (i.e. input) neuron population firing rates at time *t* and **y**_*t*_ ∈ ℝ^*n*^ the postsynaptic (output) equivalent. The synaptic modulation matrix, **M**_*t*_∈ ℝ^*n×d*^, represents changes to the network’s connections induced by some general biological mechanism through unsupervised learning. To incorporate changes in synapses due to various modulation mechanisms, we allow the modulation matrix to change otherwise *fixed* synapses generally represented by some randomly initialized matrix **W**.^7^ We consider two distinct update rules for the modulations in this work. The *associative* mechanism, Eq. (2a), dependent upon both the pre- and postsynaptic firing rates, is given by

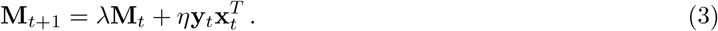

The simpler *pre-only* modulation update that is only dependent upon the presynaptic firing rates, Eq. (2b), is

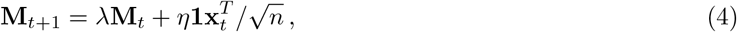

where **1** ∈ ℝ^*n*^ is the all 1s vector. In the above expressions, *η* ∈ ℝ is the learning rate of the modulations that controls the rate at which modulations are learned. Its sign determines the sign of the modulations and thus whether they *strengthen* or *weaken* the corresponding synapses.^8^ The other parameter, 0 *< λ <* 1, represents the gradual decay of changes to the weight matrix. Occasionally it will be useful to discuss the *decay timescale τ*_decay_, which is related to the decay rate via *λ* = 1 − Δ*t/τ*_decay_. Throughout this work, at the beginning of training, the modulations are initialized to be zero, i.e. **M**_0_ = **0**.

We distinguish neurons in the networks we consider between excitatory and inhibitory. A neuron that is excitatory is defined to have only positive weights leaving it so that it can only enhance the response of the postsynaptic neurons it feeds into. Similarly, an inhibitory neuron has all negative weights leaving it, ensuring it can only depress the postsynaptic neurons it feeds into. For this definition of excitation/inhibition to be meaningful, we also limit all firing rates in the network to be positive definite, including all input firing rates. Explicitly, this means the weights of our network are subject to the constraints

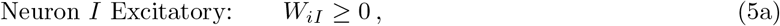

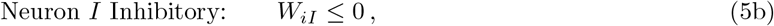

for all *i* = 1, …, *n*. Importantly, we do not allow for modulations to change the sign of weights leaving a given neuron. Per our definition above, this ensures that an excitatory neuron can never inhibit and vice-versa. Explicitly, this means that

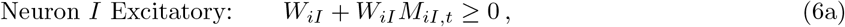

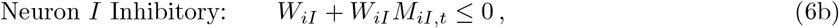

for all *i* = 1, …, *n* and all *t*.

Several experimental studies have investigated the amount of change a given neuron can undergo through mechanisms such as STSP or LTP/D. To bound the modulations to realistic values, we further restrict the modulations so they cannot enhance or depress weights beyond what has been observed in experimental settings. Since such changes can differ depending on the mechanism, we enforce different bounds for the associative and pre-only dependent modulations. Explicitly, we limit

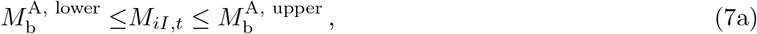

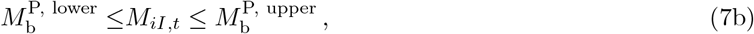

We take 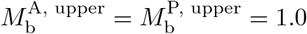, so that both types of modulations can at most double the strength of the corresponding synapses. Meanwhile, we take 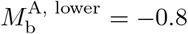, i.e. a synapse can at most be reduced to 20% of its original strength, similar to values observed in several long-term plasticity experiments [39, 64, 65] . STSP has been recently observed to almost completely suppress certain synapses [20], so we take 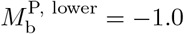. Note with these chosen values of *M*_b_, the bounds are stricter than those of Eq. (6), so in practice the enforcement of Eq. (7) implies the bounds of Eq. (6) are automatically met.

#### 4.1.2 Familiarity modulated synapse network (FMSN)

As mentioned in the main text, it is useful to first understand the properties of FMSs in a simple setting. To this end, we investigate FMS properties in a simple two layer neural network, what we call the familiarity modulated synapse network (FMSN). We take the input and output layers to have *d* and *n* neurons, respectively. The input and output layers are connected by weights, representing the synapses of a biological neural network. We will assume the synapses of the network have some underlying strength at initialization, that we denote by the weight matrix **W** ∈ ℝ^*n×d*^. We take **W** to be sparse such that its elements have magnitude

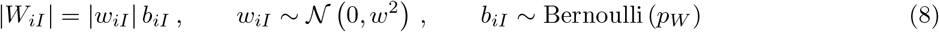

where the parameters *w* and *p*_*W*_ determine the magnitude of the nonzero elements and the sparsity, respectively. The sign of the nonzero elements are determined by the input neuron cell type (see Sec. 4.1.1 above).

Like the synapses in the brain, we allow the individual weights in our network to be modulated over time. We denote the modulations at time step *t* by **M**_*t*_ ∈ ℝ^*n×d*^, i.e. the same size as the weight matrix **W**. The combined weights and modulation matrix yield the output neuron preactiviation values,

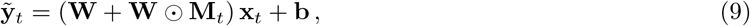

where **b** = *b***1** with **1** ∈ ℝ^*n*^ is a uniform bias term that can represent neuron firing rate thresholds as well as other network factors neglected in this simple model (see below). The parameter *b* is adjusted at initialization to ensure realistic response sparsity in the output population, but is otherwise fixed throughout training, see Sec. 4.3.1 below. Finally, the output preactiviations are passed through a nonlinear function, *ϕ*(·), representing the output neurons’ properties such as their firing rate threshold and maximum firing rate. Thus, the output population’s activity at time *t* is given by

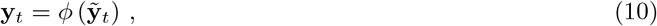

where *ϕ*(*·*) is applied piecewise. Throughout this work, we use a rectified tanh as the output neuron’s nonlinearity,

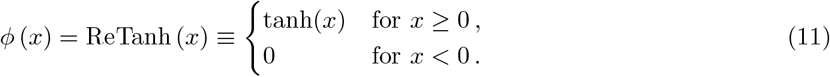

This activation was chosen since it has three desirable properties: (1) it is positive definite which was required for cell types in our model to make sense, (2) it is bounded, and (3) it is approximately linear for 0 *< x* ≪ 1, so small positive preactivation values become small activity.

The FMSN’s behavior can be changed considerably by modifying the distribution of excitatory/inhibitory neurons in the input neuron population. Note that since the neuron types only influence the sign of the weights/synapses leaving the neurons, the cell types of the output neurons do not affect the FMSN’s behavior. When we investigate the FMSN in the main text, for simplicity, we consider a setup that only has excitatory neurons in its input layer. The bias adjustment within the output population that we perform to ensure realistic firing rates helps balance excitation/inhibition in the output population that only explicitly receives excitatory input. We find that the bias term is always negative for the output population firing rates we consider, which can be thought of a uniform inhibitory input into the output neuron population that provides E-I balance.

In the supplement, to understand the effect of FMSs applied to inhibitory synapses, we also consider cases where input population of the FMSN has both excitatory and inhibitory neurons in it. In these cases, the unmodulated weight matrix, **W**, has some columns with all positive nonzero elements and some columns with all negative nonzero elements. We study a simple case where the FMS mechanism applies to synapses belonging to either the excitatory or inhibitory input neuron populations. Of course, it is also possible to have modulations acting on both populations, but we leave such investigations to future work. All other properties of the FMSN, including the firing rate adjustment, remain unchanged.

#### 4.1.3 Cortical microcircuit network

A rough schematic of the cortical microcircuit network is shown in Fig. 2c. The three primary populations we consider in the network are the excitatory, SST, and VIP neuron populations. We will index variables belonging to these three populations using *p* = E, S, V, respectively. Lastly, we also include an additional population of excitatory neurons that drive the stimulus history inputs into the VIP population and represent a subset of the top-down input into the cortical layer we explicitly model. We denote these additional excitatory neurons by the superscript ‘hist’. We do not make any attempt to model behavioral effects related to the image change task, including the licking response to the task [42].

We begin by introducing the cortical microcircuit without any FMS mechanisms added to its synapses. The preactivation response of the excitatory, VIP, and SST populations are respectively given by

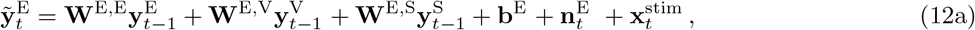

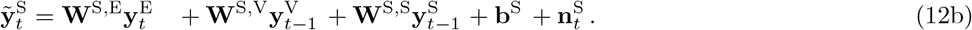

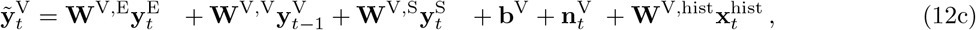

where 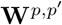 represents the synapses connecting presynaptic population *p*^*′*^ to postsynaptic population *p*, **b**^*p*^ is the bias vector of population *p*, and 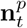 represents additional noise injection (see Sec. 4.2.2 below). Note the three populations do not update in sync: at time *t* the excitatory population’s activity is updated first, followed by the SST population, and then VIP. Asynchronous updates were found to help numerical stability. This order is also biologically motivated since the canonical input to layer 2/3 from layer 4 pyramidal neurons is much weaker to VIP and SST than pyramidal neurons.

All the preactivation responses pass through a non-linearity,

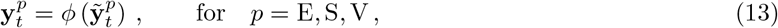

where *ϕ* (*·*) = ReTanh (*·*), ensuring the rate remains positive definite, see Eq. (11).

The excitatory and VIP populations both receive additional external input related to the stimulus change task. Specifically, 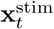 and 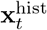 represent the present stimulus input and activity of the stimulus history excitatory neurons, respectively (see Sec. 4.2.2 below for details). Note the activity of the excitatory neurons that drive the stimulus history signal is an input to the network, so we have denoted it by 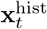 rather than 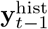 for clarity. Both stimulus inputs, 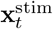 and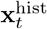, are fed directly into an excitatory population that in turn drives the rest of the circuit through sparse synapses. The primary difference between them is that the former drives an excitatory population inside the microcircuit while the latter drives an excitatory population that is assumed to reside in a higher cortical area that we do not explicitly model. The stimulus history excitatory neurons represents a subset of top-down information fed into the microcircuit and were chosen to feed into the VIP population since they are known to receive feedback input [20].

The 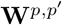 are sparse matrices whose elements are also drawn according to Eq. (8), i.e. in the same way as the weight matrix of the FMSN. Once again, the cell type of the presynaptic population *p* determines the sign of the nonzero 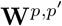 elements, and thus **W**^*p*,E^ ≥ 0, **W**^*p*,S^ ≤ 0, and **W**^*p*,V^ ≤ 0 for all *p*. For a given 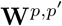, the sparsity of the synapses (i.e. *p*_W_), magnitude of the nonzero elements (i.e. *w*), and relative number of cells in each population are all set by experimental literature [17, 18, 20], see Sec. 4.4.1 below. In short, for all 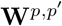, we take *p*_W_ to be the corresponding entry in Fig. S4d and *w* to be the corresponding entry in Fig. S4b, up to a multiplicative constant *c*. Importantly, *c is the same for all* 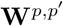, and thus the relative connection strengths between populations is completely fixed by experimental results, up to changes from FMSs.

Since **W**^V,hist^ represents an unknown subset of top-down excitatory to VIP connections, we simply sets its sparsity equal to the within-layer excitatory to VIP connections. Its synaptic strength is set to a value comparable to the within-layer excitatory connections so that the omission ramping response has a comparable magnitude to image responses.

All three biases are parameterized similarly to the FMSN, i.e. **b**^*p*^ = *b*^*p*^**1** where **1** is the all 1’s vector in the corresponding space. Once again, the *b*^*p*^ are adjusted at initialization to ensure realistic response sparsities in all three neuron populations, see Sec. 4.4.2 below.

##### Adding FMSs

So far, the network above has no FMS and thus has no way of adapting to the stimuli over time. We add the following three FMS mechanisms onto synapses feeding into the VIP cells,

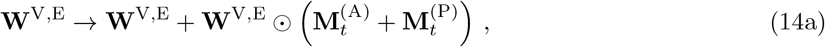

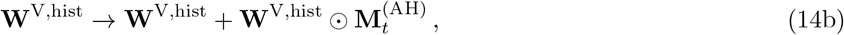

where the superscripts in (*·*) refer to distinct FMS mechanisms. Specifically, (A), (P), and (AH) respectively correspond to what we refer to as the *FMS*_A_, *FMS*_P_, and *FMS*_AH_ mechanisms. Note we have added *two* FMS mechanisms to the same set of synapses, those going from the excitatory to the VIP population. When multiple sets of FMSs are present on the same synapses, we still enforce the cell-type bounds of Eq. (6). The three distinct modulation correspond to the three novelty responses we aim to model. They are respectively subject to the following update expressions,

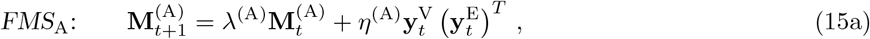

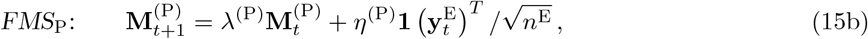

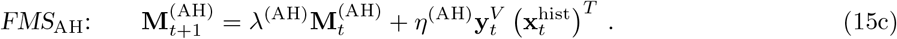

Note that the updates are distinct, but are all of the same fundamental form we have used throughout this work, see Eq. (2). That is, the associative updates are dependent upon both the pre- and postsynaptic firing rates of the populations they connect, while the pre-only is only dependent on the presynaptic firing rates since we want it to represent STSP-like modulations that occur at timescales on the order of seconds. The three FMS mechanisms are subject to the corresponding bounds motivated from experiment discussed below Eq. (7). In practice, during training, the *FMS*_A_ and *FMS*_P_ modulations rarely come close to saturating the bounds imposed by experiment, while the modulations of *FMS*_AH_ come close to their bounds at a much higher rate. For the exemplar network shown in Fig. 5, for the modulation matrix terms corresponding to nonzero synapses of **M**^(A)^, **M**^(P)^, and **M**^(AH)^, only 1.5%, 0.09%, and 42% come within 50% of their bound and 0%, 0%, and 23% come within 10% of their bound, respectively. Note for the slower modulation mechanisms, these rates were only calculated during roughly the last quarter of training time.

To adjust the parameters of the three FMSs shown above, we scan over learning rates of all three FMS mechanisms and determine which of these yields the best mean response fits, see Sec. 4.4.6 for additional details. We take 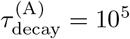 seconds, 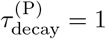 second, and 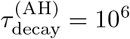 seconds based on the timescale of the corresponding biological mechanisms we wish to match onto and experimental observations (though see Sec. 4.4.7 for how these may change to expedite training).

### 4.2 Tasks

#### 4.2.1 Familiarity-novelty task

The familiarity-novelty task is used to train the FMSN discussed in Sec. 2.1. The neuronal encoding of different stimuli are represented by distinct random binary vectors, **x**^*α*^ ∈ ℝ^*d*^, where *α* indexes the distinct stimuli. The random binary vectors are chosen to be sparse, i.e. the elements of stimuli *α* are given by

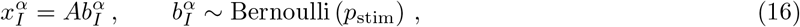

where *A* is some normalization factor. Since we generally use small *p*_stim_, we also ensure that there are a minimum number of nonzero elements for each **x**^*α*^.

Prior to training, *N*_F_ stimuli are generated and defined to be the *familiar set*,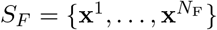. The unsupervised training consists of passing the network a sequence whose elements are randomly drawn from the set *S*_*F*_ (with replacement). After each sequence step, the network’s modulations are updated according to Eq. (2). Random Gaussian noise (iid to each element/sequence step)^9^, ***ϵ***∼𝒩 (**0**, *σ*_*ϵ*_**1**), is added to the inputs before each element is passed through a ReLU function to ensure all elements are positive,

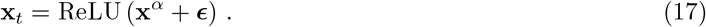

As mentioned in the main text, the sequence of familiar stimuli is ordered such that each element of the familiar set is seen every *N*_*F*_ sequence steps. The order of the familiar set is shuffled within every *N*_*F*_ window. Implicit in this training is that the time difference between successive stimuli is constant, a feature we relax in stimulus change detection task. The parameter *λ* that controls the decay can be thought of as corresponding to a decay length relative to number of examples.

During and after training, we test the network’s response to both the familiar set of stimuli as well as a novel set of stimuli, 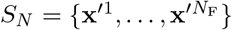, where **x**^*α*^ ≠ **x**^*′β*^ for all *α* and *β*. For these test responses, the network’s modulations are not updated after being passed through the network, so they do not affect the network’s response to future inputs. Measuring the network’s response at these steps is simply done for the sake of comparison and is not a necessary step in training.

#### 4.2.2 Stimulus change task

The task we train our cortical circuit model upon is meant to imitate the image change task used in the experimental data we are modelling [23, 24]. At any given time, the input to the network consists of three distinct parts:

1. **Present stimulus:** A stimulus input vector, 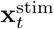, representing a neuronal encoding of the current visual input at time *t* (see Figs. S5a,b for examples).
2. **Stimulus history information:** Information about the recent history of the stimulus sequence, specifically an encoding of the amount of time that has elapsed since the last stimulus presentation, 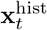(see Fig. S5c for example).
3. **Time-correlated noise:** Additional noise input into each population representing contributions to the neuronal activity from factors neglected in our model (e.g. behavioral effects), 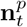 for *p* = E, S, V (Fig. S5d).

All input sequences are discretized to a time length of Δ*t* = 1*/*32 s, or 31.25 ms. This time difference is chosen to match the experimental sampling rate.

Before discussing the details of each of these inputs, we briefly review the experimental task, see Ref. [24] for significantly more details.

##### Experimental image change detection task details

The image change detection stimulus sequence consists of image presentations in quick, regular succession. Stimuli are presented for 250 ms and then followed by a grey screen for 500 ms. The same image is presented several times in a row before a new image is chosen and the process repeats (Fig. 4b). In the experiment, mice are tasked with licking in response to an image change. The number of times an image is presented in a row is between 4 and 11, with the count being drawn from a truncated exponential distribution so that 4 image presentations in a row is the most likely. Additionally, there is a 3 second grace period after an image presentation before the trial restarts. Note the image after an image change is drawn from the entire set of possible images and as such there is a 1*/*8 chance that the *same* image is drawn again. These cases are not included when measuring the network response to image changes.

The mice are first exposed to static grating and trained on a grating change detection task, which was found to improve the training time on the subsequent image change detection task. Mice are trained on the image change detection task from a set of 8 images that gradually become the *familiar set* (Fig. 4c). During these *training sessions* mice learn to perform the task. Once mice reach a particular performance threshold on the image change detection task using the familiar set, their neuronal responses are recorded over several *familiar imaging sessions* that are separated by at least one night of rest. Generally, the second of these sessions is a “passive” imaging session where they do not need to perform the task to obtain rewards (see Ref. [24] for exact training sequences). Afterwards, a novel set of 8 images are introduced into the same image change task. Without any additional training, the mices’ neuronal responses while performing the task are recorded over several *novel imaging sessions*. Once again, generally the second of these is a “passive” session and is omitted from this analysis. After at least one session of exposure to the novel imaging set, the mice’s responses while performing the same task on the novel imaging set are gathered in what is called a *novel-plus imaging session*.

During only the imaging sessions, i.e. not included in the training sessions, each image has a 5% chance of being omitted. For an omissions, the grey screen continues to be displayed for the 250 ms where the image would have been presented (Fig. 4a, middle). There is no limit on the number of omissions that can occur in a row, though longer chains become increasingly rare. Omissions cannot occur for the image presentation that would be a change or the pre-change image. This means that all omissions, including sequences of multiple omissions, are surrounded on either side by the same image.

We now discuss the three distinct parts of the input to the cortical microcircuit network.

##### Present stimulus

The present stimulus sequence is constructed to represent a neuronal encoding of the equivalent visual stimulus of the experiment. It is constructed to closely match the statistics of the image change detection task the mice are trained upon. Since image presentations last 250 ms and are separated by 500 ms of grey screen (ignoring the possibility of omissions for the moment), there are 250 ms*/*Δ*t* = 8 time steps of the neuron encoding of the image followed by 500 ms*/*Δ*t* = 16 time steps of the neuronal encoding of grey screen (though see below for additional details). This sequence then repeats, with the image identity of each 750 ms window being chosen so that image changes and omissions occur at frequencies described above. Different images encodings are represented by distinct random binary vectors drawn in an identical manner to that described in Sec. 4.2.1 above. Similar to experiment, the familiar and novel sets are chosen to have 8 distinct stimuli in them. All inputs have random Gaussian noise added to them. As observed in the experimental data, neuronal activity is low during stimulus times where the grey screen is displayed, so the present stimulus inputs representing grey screen encodings only consist of the added Gaussian noise discussed above. When an image omissions occurs, the image input is simply replaced by additional grey screen input. We allow for at most two omissions to occur in a row.

During the time steps representing an image presentation, there are three additional contributions to the stimulus sequence used to mimic the responses of experimental studies. First, to best match mice response data, we delay the mean onset of the image presentation stimulus by two time steps, corresponding to 2Δ*t* ≈ 62 ms, relative to when we consider the image stimulus has begun display. This has the net effect of shifting the neuronal responses later in time relative to the image presentation time period (for example, see Fig. 5) and approximately matches known delays of the visual cortex to visual stimuli [66] as well as the experimental data [23, 24, 66]. Second, we also smooth the input stimulus with a smoothing kernel to represent the ramping and decay of the image response to the input sequence [66]. The smoothing kernel is the normalized experimental mean excitatory response, deconvolved with the experimental stimulus smoothing function (see Sec. 4.4.5).^10^ The resulting smoothing kernel from this process is shown in Fig. S5f. Third, for each cell we allow the onset of the image stimulus to vary by *±* Δ*t* so that not all cells receive input representing the image presentation at the same time step. This incorporates effects of lag times of stimulus responses across a population and is also useful for numerical stability so that all cells do not respond in unison. We incorporate all three of the above effects on a cell-by-cell basis into a stimulus kernel 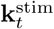. If **x**_*t*_ is the random binary vector representing the current image being presented, then the full raw stimulus stimulus input is given by

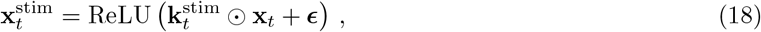

where ReLU(*·*) = max(*·*, 0) and ***ϵ*** ∼ **𝒩** (**0**, *σ*_*ϵ*_**1**). See Figs. S5a,b for exemplar present stimulus input for an image change and image omission.

Unless otherwise stated, the present stimulus inputs are taken to have average sparsity *p* = 0.05 with a minimum sparsity of 0.025 for any given stimulus. The input normalization was chosen so that nonzero elements had size 0.2. Since time-correlated noise is already injected into each population (see below), the present stimulus noise was taken to only be *σ*_*ϵ*_ = 0.03*×* 0.02.

Since mice can lick mid-sequence and this can reset the task mid-trial, the experimental distribution of number of presentations between a given image is thus fairly heavy tailed. Across the entire experimental dataset, we find there to be on average 20.4 image presentations between changes. To ensure shorter trial times, we truncate the maximum number of image presentations between a change to 75. This only omits the 2.3% of image changes on the tail, and shifts the average number of images between a change to 18.4. See Figs. S5g,h for a plot of the true number of image presentations until the next presentation. We do not find using the experimental image-change-distribution versus the idealized one that assumes no licks affects the mean response results significantly. However, in the cell subpopulation analysis the fitting metrics are dependent upon the global distribution of change occurrences and so the truncated experimental distribution was used in said analysis.

##### Stimulus history information

As mentioned above, we also pass the network information about the recent history of the stimulus, in particular the time that has elapsed since the last image presentation. This information is assumed to be encoded in a subset of the top-down input to the cortical circuit. This additional input into the network is necessary to observe responses that are dependent on the relative time between image presentations, e.g. the omission ramps. The top-down inputs could be produced from the present stimulus sequence described above using, say, a simple recurrent network that counts the time steps since the last stimulus and encodes said information in output neuron responses that match known stimulus tuning properties. As our goal for this study is the effects for FMS in the local circuit, we avoid an explicit implementation of such history encoding and simply input it directly into the network.

In this section, we denote the time that has elapsed since the last image presentation at time *t* by *s*, which is measured in seconds. For example, with no omissions, *s* = 500 ms immediately before the onset of the next image presentation. For times when the stimulus is currently being presented, *s* = 0. The time since the last image presentation is maximized after omissions, and since at most we allow for two successive omissions, 0 ≤ *s* ≤ 2 seconds.

We denote the neuronal encoding of the time *s* by **r**(*s*). We *assume that encodings of times that are close together are more similar than times further apart, as measured the dot product between the two representations*. That is, if |*s*− *s*^*′*^ |*<* |*s* − *s*|^*′′*^, then **r**(*s*) **r**(*s*^*′*^) *>* **r**(*s*) **r**(*s*^*′′*^).

The temporal encoding input is generated by creating a population of neurons that are each tuned to a particular *s*. For simplicity, we take the neuronal tuning curves to take the shape of a Gaussian, though our results easily generalize to other tuning curves such as cosine bumps. To match experimental results, we assume the population of neurons’ tuning curves are centered at times that are logarithmically distributed and that the width of each tuning curves is proportional to their center [67–69]. Specifically, we take the neuron tuning centers to be logarithmically distributed between 10^−2^ to 10^1^ seconds. For neuron *I* centered at time *s*_*I*_, its width is directly proportional to the size of the center of its tuning,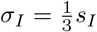. Altogether, the tuning curve of neuron *I* is

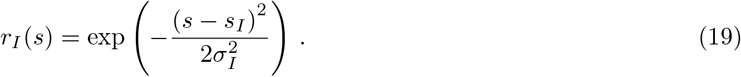

See Fig. S5i for exmplar raw turning curves. In practice, the longest delay time between images is at most two seconds, so those neurons tuned to higher times are almost always silent in our setup. The resulting neural population responses are then each individual neuron’s response to the corresponding *s* (Fig. S5j). Due to the higher density of neurons centered to small *s*, the relative magnitude of the population response vectors decreases gradually for larger times (Fig. S5k). Notably the trend in magnitude is the opposite to that of the ramping response, i.e. smaller *s* have the largest magnitude. A verification of the decrease in similarity for times further apart is shown in Fig. S5l. Since we assume this history stimulus represents some unknown subset of the total top-down input into the VIP population, we simply set the magnitude of the cell activity to be comparable to the present stimulus input so that the omission ramping has a similar response to image presentations.

Similar to the present stimulus input, noise is added to the stimulus history stimuli and thresholded to be positive definite,

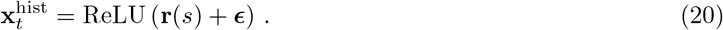

We note that this encoding of the history of what the mouse viewed is purposely simplistic and likely misses other effects that could be observed experimentally. For instance, an image that lasts longer or a shorter delay would not elicit a large response from the VIP cells despite these being outside of the the normal rhythm of the task. A more thorough encoding of the history of the task would allow the model to react to additional disruptions to the regular task flow, but we leave such exploration for future work.

##### Time-correlated noise

To account for contributions from neurons not included in the circuit, as well as contributions from other task-relevant effects (e.g. behavior), the excitatory, SST, and VIP populations are injected with additional time-correlated noise (in addition to the noise added to the present stimulus and stimulus history inputs described above).

Specifically, the noise is generated by convolving white noise with a Gaussian smoothing kernel over time, and then weighting the noise injected into each neuron to account for any variance the population may have from such effects. The continuous Gaussian smoothing kernel is given by *k*(*t*) = 1*/* (*πσ*^2^)^1*/*4^ exp (−*t*^2^*/*2*σ*^2^) with *σ* = 125 ms. The discrete smoothing kernel, *k*_*t*_, is found by evaluating the above at *k*_*t*_ = *k* (*t*Δ*t*) for *t* = −(*σ/*Δ*t*)^2^, …, (*σ/*Δ*t*)^2^. Note the normalization of *k*(*t*) is chosen such that the convolution does not change the variance of the uncorrelated noise, e.g. 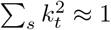 . The weight accounting for how much noise is injected into neuron *i* is *w*_*i*_ ∼ *U* (0, 1). Thus we have

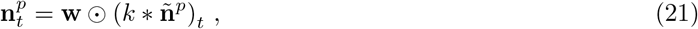

where 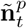 is the uncorrelated noise with 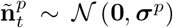. As shown above, this noise is added to the preactivations of all neurons for each population. The variance of the noise for each population, ***σ***^*p*^ = *σ*^*p*^**1**, is adjusted to match experimental baselines, see Sec. 4.4.4 below.

### 4.3 Familiarity modulated synapse network details

#### 4.3.1 Response sparsity adjustment

With only excitatory synapses in the input later of the FMSN, all inputs to the output neuron population are positive and so, without any threshold/bias term, the output population would have a response rate of close to 100% for every possible stimulus. Of course, such a response is not realistic over an entire population of neurons in the visual cortex and such excitation should be balanced by inhibition to achieve realistic response sparsities. This could be achieved by introducing inhibitory neurons with appropriate synaptic strengths into the input layer, in which case output neurons would receive somewhat balanced levels of excitation and inhibition. Indeed, we study such a network when we want to consider modulations on inhibitory neurons. However, for the FMSN investigated in the main text we choose a simpler solution that we discuss here that generalizes to the method we use to balance the cortical microcircuit below.

The output neuron population’s response rate can also be adjusted by changing the population’s firing threshold/bias, **b** in Eq. (9). In this work, we only consider **b** = *b***1** with **1** ∈ ℝ^*n*^ so that the point neurons we study only differ in connectivity and noise injection. For a given neuron, a negative *b* effectively acts as a uniform inhibition across all possible inputs. Across the entire output population, a negative bias allows the neurons to have more realistic response sparsities despite only receiving excitatory inputs.

To adjust *b* to get the desired response rate, we first draw a validation set of size 100 from the same distribution that generates the familiar and novel sets. This entire validation set is passed through the network at initialization with *b* = 0, with no adjustment to the modulation matrix **M**. Given the known activation function, Eq. (11), from the validation set’s preactivation values the bias needed to have the desired response sparsity across the validation set can be exactly computed. In short, all preactivation values (across the stimuli of the validation set and output neurons) are sorted by value, and the bias is chosen such that the desired percentage of these values are above 0. Since the familiar and novel sets are drawn from the same distribution as the validation set, this yields a similar response rate over said sets without being directly fit to them. Note this procedure means that the network has the desired response sparsities at initialization, but the induced modulations during training can change the response rate of the novel and familiar sets (Fig. S2a).

#### 4.3.2 Decoding accuracy and dimensionality

For the associative weakening FMSN example considering in the main text, the change in output activity of the familiar set significantly affects the decodability of the output signal. Post-training, decodability of stimulus identity within the familiar set is significantly lower (0.46 *±* 0.05) while that of the novel set is perfect (1.00 *±* 0.0, mean *±* std). The difficultly in decodability is reflected in the effective dimensionality of each set’s output activity: the novel outputs occupy a low-dimensional space (*D* = 6.3 *±* 1.5) while the familiar outputs are small enough that their signal is hard to distinguish from noise and thus the space they occupy is significantly higher dimensional (*D* = 48.5 *±* 7.1, mean *±* std, Figs. S2b,c). Both the above properties are a function of the amount of modulation within the network, so the decoability/dimensionality of the familiar and novel sets can vary significantly by, say, changing the modulation learning rate (Figs. S2m,n).

To compute the decoding accuracy of the familiar and novel sets, we use the same input noise that is used during training (see above) to create 1000 noisy versions of each stimulus. Each noisy stimulus is then passed through the trained network, resulting in a total of 8000 output responses across the entire familiar or novel set. Said responses are then labeled by their index within their set and a linear SVC is used to decode them using 10-fold cross validation to compute test accuracies. Specifically, we use sklearn.svm.LinearSVC with default parameters other than max iterations=1e5. The approximate dimensionality of the representations reported in the main text are computed using the participation ratio of the ratios of variance explained of the resulting PCA fits.

### 4.4 Cortical microcircuit details

#### 4.4.1 Microcircuit structure

In our model, the total strength of the collection of synapses from a given presynaptic population into a given postsynaptic neuron depends on three major factors:

1. **Connection probability:** The probability for a given synapse to exist between any two cells of the given pre- and postsynaptic type (Fig. S4d).
2. **Relative cell counts:** Along with the mean connection probability, the number of presynaptic cells in a given population affects the mean number of inputs a given postsynaptic neuron receives. Since our network does not explicitly model true cell counts found in the visual cortex, this enters as a relative value that we can then normalize by some baseline number (Fig. S4e).
3. **Synapse strength:** The strength *per synapse*, which we measure by the mean time-integrated postsynaptic potential (PSP), see below (Figs. S4b,c).

When layer-specific experimental observations are available, we take the values given for L2/3 of the visual cortex. The three factors above directly affect both the population count in our microcircuit model as well as the explicit form of the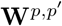. It can also be helpful to track the mean population strength, a product of the three factors above

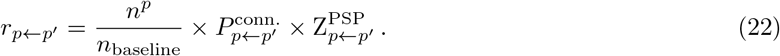

In practice, we take *n*_baseline_ = *n*^S^, which gives us the inter-population connection strengths shown in Table 1 and Fig. S4a. Although they are not explicitly included in our microcircuit model, we include data for PV neurons here as well (see Discussion for further details).

**Table 1:** Relative cell counts and inter-population connection strengths of the various cell populations. Second column shows relative cell counts for Layer 2/3, and it is assumed that 70% of Htr3a cells are VIP [18]. Third through sixth column shows population connection strengths for various pre- and postsynaptic combinations. See Sec. 4.4.1 for details on how relative population connection strengths are computed. Note, although PV neurons are not included in our microcircuit model, they are included in this table for completeness. The ‘N/A’ entry corresponds to synapses for which there is no data in Ref. [20].

|  | Rel. Count | Post Exc. | Post SST | Post VIP | Post PV |
| --- | --- | --- | --- | --- | --- |
| Pre Exc. (L2/3) | 27.35 | 0.105 | 0.750 | 1.000 | 0.908 |
| Pre SST | 1.00 | -0.081 | -0.024 | -0.356 | -0.060 |
| Pre VIP | 1.67 | -0.008 | -0.227 | -0.020 | -0.004 |
| Pre PV | 1.38 | -0.262 | -0.107 | N/A | -0.353 |

1. **Connection probability:** The connection probabilities between pre- and postysnaptic populations are computed from the fully adjusted connection propabilities of Ref. [20] (Fig. S4d). In said work, it was found that connection strength was independent of connection probability. Additionally, excitatory connections most strongly distinguished by postsynaptic connection, while inhibitory by presynaptic connection [20]. The adjusted connection probabilities of Ref. [20] are reported as fits that are dependent upon the distance between cells in addition to the dependence on pre- and postsynaptic neuron type. Since we do not explicitly simulate the spatial distribution of cells in our model, we use length scales from the experimental measurements to set distance-dependent connection probabilities. Specifically, since the imaging field of the two-photon experiment was 400 *μ*m *×* 400 *μ*m [24], we randomly generated cells locations within a two-dimensional box of this size and then computed the average connection probability between all possible pairs. The connection probability decay lengths were taken to be 100 *μ*m for E→I or I→E and 125 *μ*m for E→E or I→I [20]. From the randomly generated cell locations, this resulted in a reduction of *p*_max_, i.e. the connection probability if the cells were right on top of one another [20], by 0.25 for E→ I or I →E and 0.34 for E→ E or I→ I. Taking into account this distance-dependent reduction yields the connection probabilities shown in Fig. S4d.
2. **Relative cell counts:** We assume the microcircuit has a ratio of cell counts of Excitatory : VIP : SST found in the investigation of L2/3 of the visual cortex of mice from Ref. [18] (reproduced in Table 1, Fig. S4e). However, in order to maintain a reasonable cell counts for numerical simulation, we instead use the ratio *n*^E^ : *n*^S^ : *n*^V^ = 2 : 1 : 1, and adjust each population’s outgoing synapses to account for any discrepancy in their simulated cell count relative to their experimental cell count. For example, since in simulation there are only twice as many excitatory to SST cells, but from experimental data their ratio is closer to 27.35 : 1, we strengthen each excitatory synapse by a factor of 27.35*/*2 = 13.675 to account for the missing simulated cells. From Fig. 4g, we see this scaling of synaptic strengths to account for the differences in cell counts in our microcircuit maintains the relative population strengths.
3. **Synapse strength:** We take the time-integrated voltage over a typical postsynpatic potential (PSP) pulse fit as a measure of the synaptic strength, where

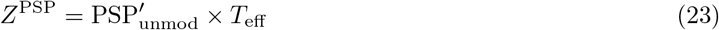

where 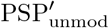 is the adjusted PSP amplitude when the neuron is not being facilitated or depressed from STSP effects [20] and *T*_eff_ is the effective time of the PSP pulse.

We compute an adjusted PSP amplitude that accounts for potential differences of the *in vitro* measurements versus what we assume to be a cell’s *in vivo* operating potential [20]. These differences are distinct across cell types, and thus can affect the relative strengths of excitation and inhibition within the cortical circuit.

To arrive at the adjustment factor, we assume the experimental current is proportional to the difference in the experimental reversal potential and the resting potential. Furthermore, we assume the *in vivo* current is proportional to the difference in reversal potential and the potential where we presume neurons are generally close to operating, which we take to be the threshold potential. The constant of proportionality in both cases is taken to be the neuron’s conductance, and thus the ratio of these differences gives the adjustment factor of the PSP value, namely

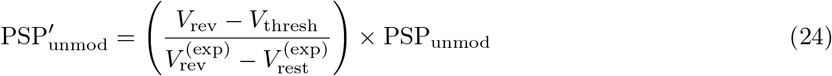

where *V*_rev_ is the estimated reversal potential of relevant channels in the presynaptic neuron, *V*_thresh_ is the estimated threshold potential of the postsynpatic neuron, 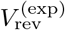 is the experimentally measured reversal potential that is dependent on neurotransmitters of the presynaptic neuron, and 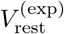 is the experimentally targeted resting potential that is presynaptic dependent.

From the literature, we use *V*_rev_ = 0 mV for excitatory [70] and *V*_rev_ = −82 mV for inhibitory presynaptic populations [71, 72]. We determine *V*_thresh_ from electrophysiology data from the Allen Cell Types Database, found at https://celltypes.brain-map.org/data [19]. Specifically, the *V*_thresh_ for each sweep and averaging over all sweeps for a given specimen identification, then averaging these values across the Cre-line. We only vary *V*_thresh_ by postsynaptic cell identity. The Cre-lines, total cell count, and computed *V*_thresh_ are shown in Table 2.

**Table 2:** Threshold voltage estimates. Values are computed across an entire session sweep and then averaged across a given specimen ID and then Cre-line. From the Allen Cell Types Database, found at https://celltypes.brain-map.org/data [19].

| Cell Type | Cre line | Cell count | $V_{\text{threshold}}$ (mV, mean±std) |
| --- | --- | --- | --- |
| Exc. | Cux2-CreERT2 | 81 | $-47.4 \pm 6.0$ |
| SST | Sst-IRES-Cre | 123 | $-41.0 \pm 7.6$ |
| VIP | Vip-IRES-Cre | 97 | $-47.2 \pm 8.7$ |
| PV | Pvalb-IRES-Cre | 217 | $-35.0 \pm 8.1$ |

Since only a small subset of synapses have experimentally measured 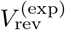, we take the median value across synapses of a given pre- and postsynaptic neuron type and use this across all cells. We do not find this impacts the resulting 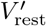 significantly. Finally, for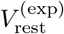, we use the targeted holding potential from experiment, which are −70 mV for excitatory presynaptic cells and −55 mV for inhibitory presynaptic cells. Junction potential corrections of −14 mV are accounted for at all steps of this calculation [19]. Altogether, these above computation yields the 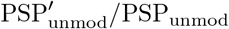 ratios shown in Fig. S4h.

The effective time of the PSP pulse is computed by integrating the PSP fits over time [20]. Up to an amplitude correction, synapse PSPs were fit using the following function

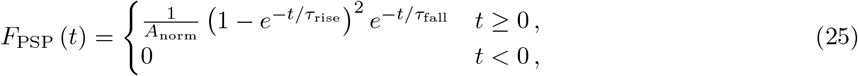

where *A*_norm_ = *F*_PSP_ (*T*_max_) with *T*_max_ = *τ*_rise_ ln (1 + *τ*_fall_*/τ*_rise_) is a normalization factor to ensure the maximum of *F*_PSP_ is equal to 1.0 [20]. Integrating this expression over time, we find the effective time of the PSP fit,

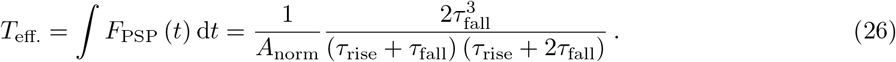

This procedure yields the values shown in Fig. S4i.

We computed *Z*^PSP^ for each synapse and then averaged across all synapses of the given pre- and postsynaptic cell type (see Fig. S4f for count). For certain pre- and postsynaptic populations, we found the fits of *τ*_fall_ were exceedingly high, and so any synapse with a *τ*_fall_ *>* 300 ms was omitted. In practice, this only resulted in a small decrease in the number of synapses for each pre- and postsynaptic cell combination (Fig. S4f).

#### 4.4.2 Response sparsity adjustment

Similar to the FMSN, the biases/threshold of the various populations are adjusted in order to set baseline response sparsity at initialization. Since we consider 3 primary populations of neurons in this work, this procedure amounts to the fitting the 3 parameters of the network at initialization, namely *b*^E^, *b*^S^, and *b*^V^ that determine the biases in Eq. (12). Again like the FMSN, since our model neglects many influences that affect the firing rates of the various populations, e.g. from inputs from other layers or from PV neurons, we assume that this bias adjustment partially accounts for the mean activity of other possible inputs. In particular, since some populations receive fairly unbalanced inputs from excitatory or inhibitory populations (i.e. the VIP population), this threshold adjustment is assumed to at least partially account for excitatory-inhibitory balance. Unlike the FMSN, our microcircuit model has recurrent connections, and so any adjustment to response sparsity at one time step affects the inputs and thus the response sparsity of subsequent time steps. This leads us to a different bias fitting procedure to account for this additional complication. Finally, note this entire procedure is performed at the network’s initialization, prior to any unsupervised training, and is thus insensitive to FMS mechanism placement or parameters.

The neuron population thresholds (*b*^E^, *b*^S^, and *b*^V^) are adjusted using supervised training to reach a certain population response sparsity over a validation set prior to training. The validation set consists of the 8 familiar input vectors as well as 504 additional vectors (for a total of 512) drawn from the same distribution. In this work, all neurons of the same population share the same firing threshold parameter, meaning particular neurons within a given population may fire more/less over the given validation set.

In particular, the response rate is adjusted with respect to the loss function

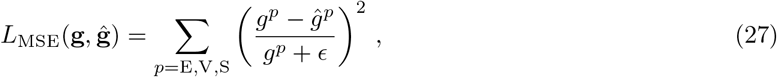

where **g** = (*g*^*E*^, *g*^*V*^, *g*^*S*^) are the experimental response rates to be matched, 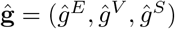 are the network’s approximate response rates, and *ϵ* = 10^−4^ provides numerical stability. Note here we use a weighted mean squared error loss so that populations with smaller response rates are treated on even footing. See Fig. S5m for exemplar fit results. For a given network population *p*, the approximate response sparsity is computed as

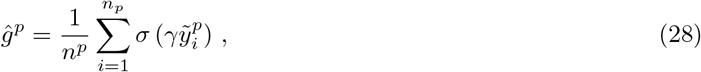

where *σ* (·) is the sigmoid function and serves as smoothed version of the step-function to enable backpropagation, *γ* = 100 controls the rate of smoothing, and 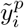 is the preactiviation of neuron *i*, see Sec. 4.1.3.

As mentioned above, since the microcircuit we investigate is recurrently connected, adjusting the response rate of one population influences the response rate of the other populations, so a self-consistent solution across all neurons must be met. To do so, the validation set is repeatedly passed to the network and the thresholds of all neuron populations are adjusted simultaneously until a self-consistent solution is found. To best resemble the stimulus sequence the network will be exposed to during the stimulus change detection task, the validation sequence is smoothed with a ramping function that matches the deconvolved signal. Response rates are computed for the populations at the peak of the ramping function, and the thresholds of the various populations are adjusted using backpropagation through time. Backpropagation is truncated to 10 time steps backwards. We used ADAM with default parameters, a batch size of 128, and shuffle the validation set every 10 network steps. Note we neglect the effect of the time-correlated noise that is injected into all neuron populations in adjusting the firing rates. Additionally, we assume the stimulus history information during images changes will be strongly suppressed, so within the validation set the corresponding part of the input is just noise.

We use **g** = (0.05, 0.4, 0.2) throughout this work to represent target firing rates during an image response in the novel session. We do not observe a large difference of the parameters *b*^E^, *b*^V^, and *b*^S^ values across different microcircuit initializations.

#### 4.4.3 Response rates and variance explained

We note that we do not aim to exactly reproduce response rates or variance explained across cells that are observed in experiment due to the significantly smaller neuron populations used throughout this work. As mentioned above, for the parameters used in our cortical circuit model, individual cells receive synaptic input from on order 10 to 100 other cells. Simulating larger populations of neurons would allow significantly more noise to be injected into each individual cell since each cell, on average, has a smaller effect on output behavior of other neurons.

#### 4.4.4 Noise injection and matching baseline responses

From the experimental data, we see that all neuron populations exhibit a nonzero baseline mean response between image stimuli (Fig. S6). Said baseline responses carry almost no information about image identity or task information except within a small window after the image stimulus turns off, indicating they may represent neuronal activity unrelated to the image change detection task [24]. To model the effects of these baseline responses, we inject time-correlated noise directly into each neuron population. We adjust the variance of this time-correlated noise to match the baseline response values observed in each population using supervised training. Similar to the firing rate adjustment considered above, this again corresponds to only one number per population, so this procedure fits a total of 3 parameters at the network’s initialization (and occurs after the firing rate adjustment). Once again, since this procedure occurs at network initialization, it is completely independent of FMS placement or parameterization.

We define the experimental baseline responses to be the mean population response halfway between the pre-change image and the change image. With this definition, for a given population, the mean population response doesn’t change much between the familiar and novel sessions, so we average across the two sessions. Taking the novel session values yields baseline targets of 1.5 *×* 10^−3^, 6.1*×* 10^−3^, and 3.4 *×* 10^−3^ for the excitatory, VIP, and SST populations, respectively. Once again, we weight how well our model fits the experimental data using a weighted MSE loss,

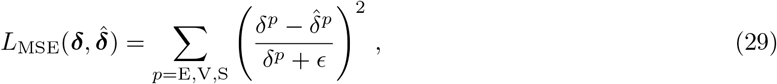

where now ***δ*** = (*δ*^*E*^, *δ*^*V*^, *δ*^*S*^) are the experimental baseline mean responses to be matched, 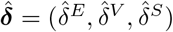 are the network’s baseline mean responses. The network’s baseline responses are simply the mean over each population response,

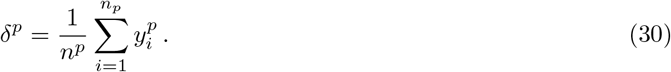

For simplicity, to initially fit the amount of noise injected into each population, we inject uncorrelated noise with standard deviation *σ*^*p*^. In practice, we find the addition of time-correlation causes a negligible change in the networks’ baseline responses (and the kernel used to generate uncorrelated noise is chosen such that it has approximately the same variance as its uncorrelated counterpart). See Fig. S5n for exemplar fit results. Similar to the above firing rate adjustment, we pass uncorrelated to the network as the input stimulus repeatedly until it reaches a self-consistent solution. We once again use ADAM with default parameters and truncate backpropagation through time to 3 network passes backward. Both the present stimulus and stimulus history information parts of the input are assumed to be just noise for the validation set. Due to additional contributions from the stimulus history input during grey screen times, the VIP baseline was found to overestimate the noise needed to fit the data, so the noise injection was reduced after fitting by a fixed percentage across all initializations.

#### 4.4.5 Response smoothing

To get the raw cell responses, we convolve the output function with the same half-normal function to match the responses of Ref. [24]. Explicitly,

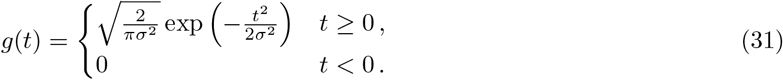

with *σ* = 60 ms (Fig. S5a). We discretize the kernel along time steps separated by Δ*t* and normalize such that the summed amplitude is equal to 1.0.

#### 4.4.6 Fitting FMS learning rates

To determine the learning rates of the three FMS mechanisms we add to the microcircuit network, we perform a brute-force scan over grids of learning rates and evaluate how well the resulting modulation effects fit the novelty data. Note that procedure still amounts to fitting only three numbers: *η*^(*A*)^, *η*^(*P*)^, and *η*^(*AH*)^. We evaluate the parameters by seeing how well their mean image change and omission responses match the experimental data. Specifically, we compute the MSE loss of the mean fit to the experimental data over all three cell populations,

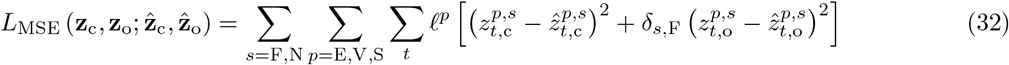

where 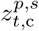 is the experimental mean image change response of population *p* in session *s* at time *t* relative to the image change, 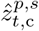 is the cortical circuit model equivalent, 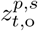 is the omission equivalent, ℓ^*p*^ is a population-dependent weight, and *δ*_*s*,F_ is a Kronecker delta function ensuring omission loss is only computed during the familiar session (since we do not try to model the suppressed omission of novel sessions). We take ℓ^V^ = 5 and ℓ^E^ = ℓ^S^ = 1, so that fitting the VIP response is more important than the excitatory or SST populations. The sum over *s* represents a sum over the familiar and novel sessions. The sum over *t* represents a sum over the relative time to the change/omission for a given mean response. We take the relative time window to be 25 time steps before and after the corresponding image change/omission event, which is roughly *±*800 ms.

#### 4.4.7 Training schedule details

As discussed in the main text, the mices’ training schedule consists of many sessions that each last on the order of one to two hours [24]. Several sessions are required for the mice to learn the task completely, meaning often they have been exposed to on the order of ten hours of the image change task to achieve the task performance threshold needed to progress onto imaging. Since neuronal responses are only collected after this performance threshold is achieved, it is not yet known how many sessions are required for the neuronal responses to the familiar image change task to stabilize.

For numerical tractability, we do not explicitly simulate the full tens of hours of the training sequences for the microcircuit model. Instead, we expose the model cortical circuit to a shorter version of the task and increase its learning rates so that it achieves stabilized responses to the familiar data over a shorter simulated time. As we saw in the FMSN, higher learning rates are capable of becoming familiarized with responses at a quicker rate, at the cost of fitting the noise to a greater degree. Thus, explicitly simulating full training/imaging times at equivalent lower learning rates should only improve the results we have shown throughout this work. Additionally, as we mentioned above, we did not find that using that exact distribution of image change times affected any results outside of the cell subpopulation analysis, and thus to further expedite training we reduced the number of repeated presentations that are between each image change to between 4 and 9.

An explicit demonstration of the training time equivalence is explored in the FMSN in Figs. S2o and S2p. There, the FMSN is trained on sequences that vary in length over two orders of magnitude and it is shown that, by correctly adjusting *λ* and *η*, the networks essentially develop almost identical responses despite the large differences in training time. For two training sequence lengths *T* and *T* ^*′*^, the equivalence is given by

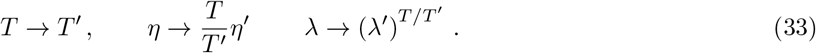

Thus, if *T* ^*′*^ *> T*, this corresponds to a reduction in the learning rate and decay rate for the longer train sequence. Note the last relation is equivalent to 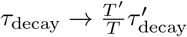.

Specifically, we train our cortical circuit models on 2000 seconds of change detection, which consists of approximately 2660 total image presentations or 330 presentations of each familiar image. With the expedited image change time, the training session consists of roughly 400 image change events.

To gather responses in cortical circuit model’s ‘imaging’ sessions, we monitor the network’s responses while it continues to train. We measure the network’s responses on 250 seconds of change detection, but we take advantage of batch training to allow us to gather responses to several distinct input streams simultaneously. However, since the cell coding analysis is dependent on statistics over an entire session, we use the actual image change distribution when collecting data for said analysis.

Since we would like to simulate the gradual familiarization of the novel set during the novel imaging session, but we have used a larger learning rate to expedite the training procedure, we reduce the learning rate of both *FMS*_A_ and *FMS*_AH_ during imaging sessions. There is evidence the mices’ response changes between sessions even without explicit exposure to the stimuli [24]. This may be due to replay. To simulate this additional familiarization that occurs between sessions, as well as additional stimulus exposure during the passive session, we train the networks on the novel images for an additional session equal in length to the imaging sessions, but at a higher learning rate, similar to training.

### 4.5 Cell subpopulation analysis

We reproduce the functional cell subtype analysis pipeline of Ref. [24] to compare our model to experimental results on equal footing. Here we give a summary of said pipeline for completeness, additional details and justification for certain parameters we match to the experimental analysis can be found in Ref. [24]. Throughout this section, we suppress indices that indicate the population and session of a given cell unless needed.

#### 4.5.1 Experimental data

We take the computed coding scores directly from Ref. [24]. The codings scores are for cells collected across several different brain areas and layers. Although our cortical circuit model specifically takes cell counts and connection data for L2/3, we note that the vast majority of VIP cells were found in upper cortical layers and there does not appear to be a significant difference in coding scores with brain area [24].

The experimental coding score analysis focuses on four primary input feature categories (also called ‘components’): images, omissions, behavioral, and task. These feature categories are further subdivided into various features that each have their own kernel and input data. For example, the image feature category contains one feature for each of the eight possible images in the corresponding image set. When a feature category is removed to compute its coding score (see details below), all feature kernels within that category are removed. See Ref. [24] for additional details.

#### 4.5.2 Model fitting

To understand how the various features coded in the task explain individual cell activity across the VIP, SST, and excitatory populations, we fit each cell’s activity using a linear regression model with time-dependent kernels. The feature categories we consider are image presentations, omitted images, and image changes. With the exception of behavioral feature category, these are the same categories considered in Ref. [24]. Also note that the image change feature category can no longer be divided into behavior-dependent features representing hits and misses.

We thus define the ten time-dependent features vectors; 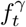 for *γ* = image1, image2, …, image8, omission, change; to have value 1 at the onset of a given feature and to be 0 otherwise (Fig. 6a, top). These features each belong to one of three feature categories; *α* = image, omission, change; with the eight image features belonging to the ‘image’ feature category, and the omission and change features belonging to the category of the same name.

For each cell *i* in each ‘imaging’ session, we fit its full session response (post smoothing, see Sec. 4.4.5), *y*_*i,t*_, using time-dependent feature kernels, 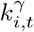, such that an estimate of its response is given by the convolution

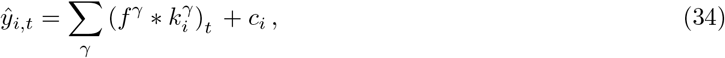

where *c*_*i*_ is bias term. Each kernel’s width in time is matched to that used in Ref. [24]: the image, omission, and change kernels persist for 0.75, 3.0, and 2.25 seconds after the corresponding feature onset, respectively.

The kernels 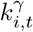 and bias terms *c*_*i*_ are fit using ordinary least square regression with an L2 penalty (i.e. ridge regression, see Ref. [24] for additional details). We evaluate the fit of the models by computing their *variance explained* on a test set,

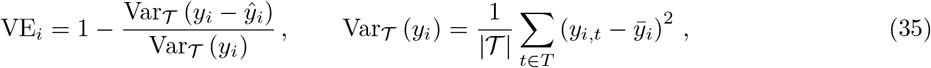

where 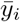 is the cell’s mean activity over the entire imaging session. Here, the 𝒯 subscript on Var_𝒯_ indicates the subset of sequence times over which the variance is computed |𝒯|and represents the number of time steps (see below). To find the optimal L2 regularization, we scan over regularization coefficients, evaluate said fits, and choose the regularization that yielded the highest mean variance explained across the entire cell population. Train/test splits are computed over distinct batches.

Since certain feature categories are quite sparse across the full input sequence (e.g. omissions and changes), their corresponding feature kernels influence only a small subset of sequence time steps. To account for the different possible kernel coverage over the entire sequence, below it will be useful to compute the variance explained over only the subset of sequence time steps where a given feature category’s kernel(s) could have possibly had an influence. Let 𝒯^*α*^ be the set of time steps a feature category’s kernel(s) could have possibly influenced the response given the sequence’s feature vectors 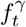 and the kernel widths in time. We define the *adjusted variance explained* as

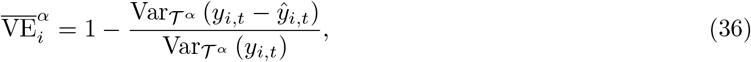

where the variance is now only computed on the subset of sequence times 𝒯^*α*^. Since |𝒯|^*α*^ is the number of time steps in 𝒯^*α*^ and |𝒯|is the total number of sequence time steps in the session, |𝒯^*α*^ |*<*|𝒯|, for all three feature categories we consider. Specifically, ^*α*^ *<* 0.95, 0.19, and 0.16 for the image, omission, and change categories, respectively. Lastly, note that from our definition in Eq. (35), the adjusted variance is always computed relative to the mean cell activity over the entire session.

#### 4.5.3 Coding scores

For each cell in each session, we compute its *coding score* with respect to each of the three feature categories we introduced above. Intuitively, a category’s coding score represents how important its feature(s) are for fitting the cell’s response. To compute coding score, we compare the cell’s adjusted variance explained of a model fit *without* a given feature category’s kernel(s) to the model fit with all kernels. Explicitly, the raw coding score is defined as

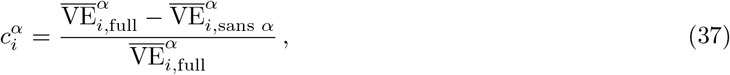

where 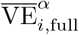 and 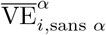 are the adjusted variance explained of the models fit with all kernels and all kernels except those belonging to category *α*, respectively.

Finally, it will sometimes be useful to compare coding scores across sessions, in which case we want to normalize all coding scores on equal footing. The *across-session coding score* of session *s* is defined as

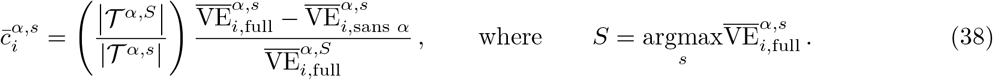

Since we have three feature categories *α* and three sessions *s*, each cell will have a 9-dimensional across-session coding score vector.

In experiment, a minimum variance explained is required for a nonzero coding score. Specifically, 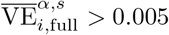 . Relative to the full fit variance explained, approximately 54.5%, 34.7%, and 52.1% of VIP cell fits fall under this threshold in the familiar, novel, and novel-plus sessions, respectively. Since the cortical microcircuit model has overall higher variance explained for all cell populations, we adjust this minimum coding score threshold to compensate for the different distribution. Setting a threshold of 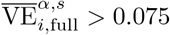 results in a similar rates as experiment, namely 56%, 20%, and 54% of VIP cells across initializations fall under this value.

#### 4.5.4 Cell clustering

We use spectral clustering to cluster the set of 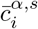 for each population. In this subsection, we use **c**_*i*_ to denote the 9-dimensional coding score vector of cell *i*.

To compute the ideal number of clusters for cell population, we use two measures: the gap statistic and the eigengap. For the gap statistic, we scan over cluster sizes from *k* = 2 to 15. We use the `SpectralClustering` method from `scikit-learn` with default parameters and a given *k* to fit the data and compute the pairwise Euclidian distance within each cluster. Let the *n*th cluster contain the set of cells indexed by *i*_*n*_. Then,

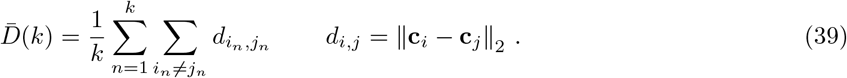

This metric is computed for the actual clusters and compared to a baseline of shuffled data. The shuffled data is the across-session coding scores shuffled across experience-level and feature categories. For the metric over the shuffled data 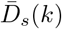, the *gap statistic* is then 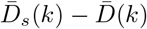, and the optimal *k* is the one that maximizes this metric (Fig. S8e).

To compute the *eigengap*, we compute differences in consecutive (ordered) eigenvalues of the Laplacian of the coding score’s affinity matrix. Specifically, the affinity matrix has elements 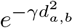, with *d*_*a,b*_ the Euclidean distance computed above. The eigengap is then the difference in eigenvalues of the Laplacian, where large gaps are associated with sudden changes of the amount of similarity explained by additional cluster partitions (Fig. S8d).

Once the optimal number of clusters is computed, we perform spectral clustering on the set of coding score vectors of a given population for 150 different initial seeds. Across all these fits, we compute the symmetric matrix of co-cluster probabilities for all cell pairs. This co-clustering matrix is then passed through `scikit-learn’s` `AgglomerativeClustering` method, again with the optimal number of clusters as determined above, with default parameters except for affinity=‘euclidean’ and linkage=‘average’. Finally, cell clustered are ordered by mean across-session coding scores, with the clusters with the smallest mean being ordered first.

### 4.6 Figure details

#### Figure 1 details

The equivalent plot for the pre-only dependent update rules is given in Fig. S1a.

#### Figure 2 details

The equivalent plot of 2c for strengthened modulations is shown in Fig. S1b.

For Figs. 2[d-g], an exemplar FMSN was trained using a set of 8 familiar stimuli. To ensure an equal distribution of the familiar stimuli over the training time, a training schedule where each of the 8 familiar stimuli is shown every 8 inputs is used. Note the order of the 8 examples is shuffled within every 8 inputs. The FMSN has a population size of 300 input and 500 output neurons. As mentioned in the main text, we take all input neurons to be excitatory. The distribution of **W** elements is taken to have *w* = 1*/*300 and *p*_*W*_ = 0.2, see Eq. (8). For the stimuli, the sparse random binary vector population has *p*_stim_ = 0.05, with a minimum number of nonzero inputs of 1, and nonzero elements of size 0.15.^11^ The standard deviation of the Gaussian noise added to the inputs is taken to be 0.1 times the size of the nonzero elements. We threshold is adjusted at initialization such that the output population has a firing rate of 30%. The associative modulations obey the bounds discussed below Eq. (7). We take *η* = −5*×* 10^2^ and *τ*_decay_*/*Δ*t* = 2*×* 10^4^ so that the modulations undergo practically no decay during the stimulus learning period. These parameters are used in FMSNs throughout this work unless otherwise stated.

Note in order to track both the familiar and novel output activity throughout training, we treat them as “test sets” when we pass them to the network, which distinguishes them from the sequential training set we use to change the modulations of a network. For any input that belongs to the test set, *we do not update the synapses*. In this way, we can understand what would be the network’s response to these various stimuli without actually updating the network’s modulations as if it truly “saw” the stimuli during training. The full familiar and novel sets were treated as test sets in order to track their output activity over training shown in Fig. 2e.

In Fig. 2g we introduce the idea of an *important synapse*. An important synapse is stimulus dependent, and as such a given synapse can be important for multiple stimuli. Important synapses are also defined before any modulations in the network occur, and are thus independent of the FMS mechanism. We define important synapses in our model as those synapses that satisfy two requirements: (1) there must be a synapse there and (2) the synapse’s pre- and postsynaptic neurons both fire when the stimulus is input into the network (without modulations). Formally, for a given stimulus input **x**, let **y** be the corresponding activity of the output layer *without any synapse modulations from FMSs*. For example, for the FMSN, **y** = *ϕ* (**Wx** + **b**), but this generalizes to other possible postsynaptic expressions. The mask **S** that defines the important synapses contained within **W** for stimulus **x** is given by

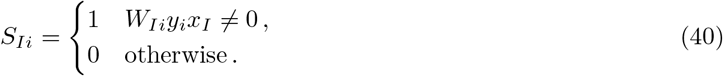

Intuitively, whether or not a synapse is important tells us whether or not said synapse would be modulated from an associative FMS mechanism. In practice, since we add noise to all stimuli being passed through the FMSN, many synapses that are not important are also modulated.

#### Figure 3 details

Figure 3a quantifies how a vector’s distance from the familiar subspace influences its output magnitude. The familiar subspace is defined as the subspace spanned by the familiar set of vectors. To measure the distance of any random vector **v** ∈ ℝ^*d*^ to this subspace, we orthonormalized the familiar set to obtain the matrix 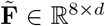 and then formed the projection matrix onto the familiar subspace via, 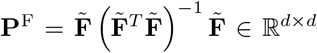. A given vector’s distance from the familiar subspace is then measured by calculating the cosine similarity of the vector and its projection onto the familiar subspace

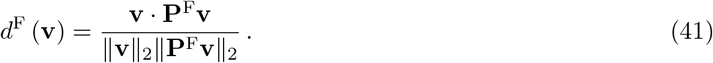

By this definition, any vector from the familiar set or any linear combination thereof has *d*^F^ = 1.0. Note that noise is added to all vectors before being passed through the network, and this noisy vector was used to compute *d*^F^ in Fig. 3a. Hence, even the noisy familiar vectors do not lie exactly in the familiar subspace.

Since simply drawing from the same distribution that generated the familiar and novel sets almost always generates stimuli that are far from the familiar subspace, we generated inputs as follows. We first drew a vector **v**^*′*^ from said distribution and also for each vector drew *f* ∼ *U* (0, 1) and a random vector from the familiar set **f** . Then, each element of the final vector **v** has probability *f* to be the same as **f**, and is otherwise equal to the corresponding element of **v**^*′*^. In this way, as *f* varies from 0 to 1, we interpolate between vectors that are drawn from the original distribution (i.e. the novel set) to those in the familiar set. Finally, to generate random vectors in the familiar subspace, 8 binary weights were drawn, their sum was normalized to 1, and these weights determined the linear combination of familiar vectors that formed the new vector within its subspace.

Figure 3b and 3c show how the evolution of the modulation matrix can change with different learning rates, *η*, and decay rates, *λ*. Both setups use a more specialized learning schedule than those in Fig. 2. Figure 3b consists of a single input vector passed over and over again to observe how quickly modulations can grow and when they saturate. Figure 3c consists of a single input vector passed on the first time step and then only noise afterwards. The goal of this plot is to observe how quickly the modulations created by a single input can shrink over time.

Figure 3d shows the average result of several KS tests on a network as we scan over number of exposures and the learning rate, *η*. With the exception of the parameters scanned over, the network and training parameters used in this setup are identical to that in Figs. 2[d-g]. KS test results are averaged of log_10_(*p*) over 10 distinct network/stimuli initializations. Figures 3[e-f] are the same as Fig. 3d, but now scan over the FMS’s decay time constant and learning rate.

#### Figure 4 details

Additional details of the experimental training procedure can be found in Ref. [24]. Details of the model network (Fig. 2c) and task (Fig. 2d) can be found in Secs. 4.1.3 and 4.2.2, respectively.

In Fig. 4g, for synaptic matrix 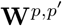 between presynaptic population *p* and postsynaptic *p*^*′*^, the model connection strength values were computed via

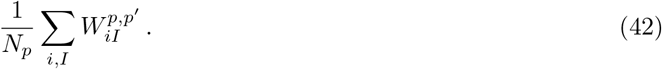

Note here *p* and *p*^*′*^ are treated on unequal footing to account for the fact that, for a given postsynaptic cell, the relative count of presynaptic inputs contributes to its total activity. The theoretical values were computed via Eq. (22).

#### Figure 5 details

Unless otherwise stated, we take the cortical circuit model to have 400 excitatory, 200 VIP, and 200 SST neurons, though see Sec. 4.4.1 for how synaptic strength is adjusted to compensate for deviations from realistic cell counts. Weights are initialized as described in Sec. 4.1.3, with multiplicative constant *c* = 0.18. The three FMS learning rates are scanned over to determine the best fit to experimental responses, see Sec. 4.4.6.

Figures 5a and 5h show comparisons of the mean responses of our cortical microcircuit model and responses measured in experiment. We match event traces that are smoothed by a half-normal filter, see Sec. 4.4.5. See Ref. [24] for significantly more details on the experimental details including details about the event extraction. To extract the mean responses of the model, let the set of all times of interest (e.g. for image changes or omissions) be denoted by 𝒯 . We denote the mean response as

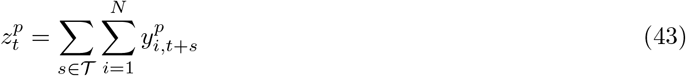

The full familiar and novel set responses of Fig. 5b are gathered similarly to the test sets of the FMSN. That is, a test set consisting of the stimuli from the familiar and novel sets representing the image changes is passed to the network at particular times during training. No temporal history or time-correlated noise input is passed to the network in these test sets so that the VIP’s change in response to the present input from *FMS*_A_ can be isolated (the temporal history response also changes during training from *FMS*_AH_). Once again, we do not update the network’s modulations during these passes, and so the microcircuit has no memory of viewing these stimuli that are solely used to monitor training progress. The test set is evaluated at every image change during training, specifically at the step corresponding to the peak of the smoothing kernel (see Sec. 4.2.2 above). The modulation magnitudes of Fig. 5c are computed analogously to Fig. 2f and also measured at each image change during training.

Figs. 5d, 5g, and 5k all show the mean responses as a particular FMS learning rate is varied. Networks and tasks are initialized identically to those used to produced the analogous figures in Figs. 5a and 5h, the only thing that changes is the corresponding FMS’s learning rate and thus its asymptotic modulations.

The responses in Fig. 5e show the VIP response of a test set, evaluated in an identical manner to that described for Fig. 5b above. The test set now consists of all novel stimuli with the temporal history part of the input corresponding to the zero-time encoding and no noise input (since the *FMS*_AH_ modulations have stabilized, they no longer significantly affect the change in VIP responses for the relevant timescales shown here). Test responses are again gathered at the step corresponding to the peak of the smoothing kernel. The responses shown are averaged over 50 image changes measured during a novel session. Additionally, the responses are normalized by the largest mean response. Fig. 5f shows the modulations of *FMS*_P_ during an exemplar image change of the novel imaging period. The modulation magnitudes shown are gathered at every time step.

Fig. 5i shows an exemplar mean VIP response to all the encodings of time-since-last image from the stimulus history input. These responses are once again gathered as a test set (see above), where now the present stimulus and time-correlated noise inputs are taken to be zero so that the VIP’s change in response from *FMS*_AH_ can be isolated. Responses are gathered before and after training on the familiar image set. The modulation magnitudes shown in Fig. 5j are gathered analogously to those in Fig. 5c.

#### Figure 6 details

Fig. 6a shows exemplar input features and fits over time. The top subfigure dots correspond to when the corresponding input features is on, see Sec. 4.5 for additional details. The bottom subfigure shows exemplar raw cell data, full kernel fit, and fit without the image kernels. Fig. 6b shows the clustered VIP *across-session* coding scores from experiment [24]. The middle plot shows cluster-average coding scores and the right plot shows average coding scores across all VIP cells. Fig. 6c shows the clustered VIP coding score from the model, see Sec. 4.5 for details of how these are computed. Clustered are ordered by smallest mean coding score to largest. Fig. 6d shows image kernel fits from the full kernel regression model. Image kernel fits are averaged across all 8 image kernels, all VIP cells, and all initializations. Fig. 6e shows fits and raw data of the cluster-aveeraged novel image (across-session) coding score as a function of a cluster-averaged network property, see details of Figs. S9 and S10 for full details of this network property and all others. Fits are done using linear regression and we use the resulting correlation to measure how much a network property influences the value of a given across-session coding score. This plot shows only one exemplar network property and one coding score, the resulting correlations across all 16 network properties we investigate and all 9 coding scores can be found in Fig. S9. In Fig. 6f we show the median correlation for the familiar and novel image coding scores as a function of two network properties for both the cluster-averaged values and the raw cell data. Fig. 6g shows the amount of familiar and novel input cells belonging to a particular cluster receive. Each point represents a single cluster for a given initialization. Points are colored by whether they are familiar-coded (cluster 3), novel coded (clusters 2, 5, 7), both familiar and novel coded (clusters 6, 8), or not image coded (clusters 1, 4). Fig. 6h is generated similarly, with clusters colored by whether they are omission coded (clusters 4, 7, 8) or not (all other clusters).

#### Figure S1 details

Fig. S1a shows the equivalent of Fig. 1b for the pre-only modulation mechanism.

Figs. S1[c-f] use identical parameters to the FMSN in Fig. 2d, but with *η* = −5 *×* 10^0^ and *λ/*Δ*t* = 60. Figs. S1[g-j] also use identical parameters to the FMSN in Fig. 2d, but with *η* = 5 *×* 10^0^. Figs. S1[k-n] also use identical parameters to the FMSN in Fig. 2d, but with *η* = 1 *×* 10^1^ and *λ/*Δ*t* = 180.

#### Figure S2 details

Figs. S2[a-d] show the additional properties of the example network shown in Figs. 2[d-g]. The output sparsity in Fig. S2a is thresholded above a minimum value to remove contributions from negligibly small activity. In particular, we defined an output to be active if it is greater than 0.01 times the mean nonzero activity at initialization across the entire validation set. For the example shown, this gives a threshold of roughly 10^−5^. Figs. S2b,c show PCA projections of the output activity of the familiar and novel sets, respectively. PCA is fit to the novel and familiar set independently in these plots. The percentages of modulations shown in Fig. S2d are all relative to the number of nonzero synapses. The “modulation” curve simply counts the number of synapses that have a nonzero modulation. Since this value approaches 1.0, the majority of synapses that can undergo modulation have been modified, but many of these modulations are quiet small and simply due to noisy activity. To compute the number of modulations that have been modified by 50% and saturated the bounds of Eq. (7), we require the modulation to be above 50% of the weights original magnitude and within 1% of the bounded value, respectively. Many modulations are quite close to the bounded value but do not exactly saturate it because of the gradual decay from the *λ* term in Eqs. (3) and (4).

As with the main text figure, to generate the rest of the figures in this supplemental figure we considered the FMSN with: *n* = 500, *d* = 300, a training length of 80 (so that each familiar example is shown 10 times), and all excitatory neurons in the input layer. Gaussian noise with a standard deviation of 0.1 times the maximum input was added to all inputs. The weights are generated from a sparse half-Gaussian distribution, with *p* = 0.2 of any connection being present. Bias is adjusted so there is a 30% firing rate at initialization. Modulations decay with a timescale of *τ*_decay_*/*Δ*t* = 2 *×* 10^4^ steps.

Figures S2[e-j] were generating by scanning of over various FMSN network/task parameters and performing a KS test on the L1 magnitudes of the familiar and novel sets post-training. All were averaged over 10 distinct initializations of the network and the task. Figs. S2e,f are scans over the lower and upper modulation bounds with grey vertical lines representing the values that have been chosen based on LTP/D changes found from experiments (see Eq. (7)). Fig. S2g scans over the noise scale, *σ*_*ϵ*_, see above Eq. (17), and the length of the training sequence. Fig. S2h scans over the number of input and output cells, *n* and *d*, respectively. Fig. S2i scans over FMSN initialization parameters, namely the weight sparsity, *p*_*W*_, see Eq. (8), and the target response rate of the response sparsity adjustment, see Sec. 4.3.1. Note that for low weight sparisities, the target response rates are often not met. Fig. S2j scans over the random binary vector parameters, namely the sparsity, *p*_stim_ and the magnitude, i.e. the equivalent of varying *A*, see Eq. (16).

Figs. S2k,l were generated by training the FMSN on different sized familiar sets. Note that we keep the number of exposures to each familiar stimulus constant, namely 10 exposures to each familiar stimuli, so as the size of the familiar set increases so does the training time. Since training is independent of the novel set, the size of the novel set is always taken to be 128. Additionally, to put the network in a more optimal parameter setting to generate distinct inputs, *η* = − 10^4^ was used. In Fig. S2k we use a different measure than the usual KS-test because its *p*-values are heavily influenced by the size of the distributions being compared. The fact that the mean responses of the familiar and novel sets begin to overlap for larger familiar set sizes indicates that members of the two distributions become harder to distinguish from one another.

Figs. S2o and S2p investigate how networks trained on a different number of familiar examples evolves approximately the same when their learning and decay rates are appropriately varied. We train identical FMSNs on the same set of familiar images for 240, 2400, and 24000 time steps. To compensate for the different number of examples each of the networks see, we lower the learning and decay rate appropriately. Let *η*_0_ and *λ*_0_ be the learning and decay rates of the network with the fewest examples, the number of which we denote by *N*_0_ = 240. Then, a network with *N* total examples has

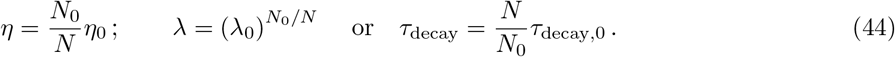

That is, networks trained on more examples have their learning and decay rates appropriate decreased or, equivalently, their decay timescale increased. Figs. S2o and S2p show the evolution of a single network on a single set of familiar examples.

#### Figure S3 details

Fig. S3b tracks the magnitude of the **M**_*t*_ in identical networks that only differ in the sign of *η*. The networks are identical to that used in Fig. 2, with *η* = 10^4^ and *η* = − 10^4^. Figs. S3c, d show results form the same networks, with the output magnitudes of the familiar and novel sets as a function of training time. Since all input neurons are excitatory, strengthening corresponds to *η >* 0 and weakening corresponds to *η <* 0. Fig. S3e, f show results from several networks with varying *η* magnitude. The dotted unbounded magnitudes are computed by training identical networks with the modulation bounds removed. Fig. S3h shows schematic of FMSN networks with both excitatory and inhibitory neurons in the input layer. For the results shown in Figs. S3i,j we specifically consider a network with an equal chance of each neuron in the input layer to be excitatory or inhibitory. Additionally, we increase the number of neurons in the input and output layers to be 1000, and increase the percent of synapses present to be 40%. Note this still means that each output neuron has only 400 synapses on average, well below what is found biologically.

#### Figure S4 details

Details of the connection strength computations shown in this figure are discussed in Sec. 4.4.1 above. Figs. S4a,b show relative values that are normalized by the maximum magnitude across the 16 possible pre- and postsynaptic combinations.

#### Figure S5 details

Figures S5[a-d] show exemplar plots of the three types of input into our cortical circuit model, see Sec. 4.2.2. Figures S5[a-c] show the L1 magnitude of the corresponding input. Similarity in Fig. S5l is simply the normalized dot product of **r**(*s*) ·**r**(0.5).

#### Figure S6 details

The mean responses are gathered analogously to those in Figs. 5a, h.

#### Figure S7 details

Mean responses as a function of learning rate are gathered analogously to those in Figs. 5d, g, k.

#### Figure S8 details

Note since the task feature category we use in our model is equivalent to the combined hit and miss features used in the experimental analysis, we plot a comparison of its learned kernel to both the experimental hit and miss kernels in Figs. S8b, c. Figs. S8d and S8e show the eigengap value and gap statistic used to determine the number of clusters for each neuron population, see Sec. 4.5.4 for details. Since there is no well-defined way of combining these two measures to choose an optimal number of clusters, an optimal number of clusters was chosen by inspection of both metrics, as in experiment [24].

Figs. S8[f-g] show the cluster membership across the 10 different network intializations.

#### Figures S9 and S10 details

For a given VIP cell, the metrics that are plotted are as follows:

- Raw excitatory input: Mean of **W**^V,E^ over presynaptic cells.
- Excitatory input (familiar): Mean of 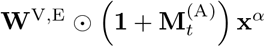 over the familiar set, where 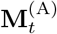 is measured at the beginning of the familiar imaging session. Note 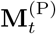 dependence is neglected here because if varies over the entire familiar session. Including 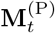 does not noticeably change correlation results.
- Excitatory input (novel): Mean of 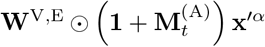 over the novel set, where 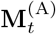 is measured at the beginning of the novel imaging session.
- Excitatory input (novel plus): Same as above, but 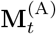 is measured at the beginning of the novel plus imaging session.
- Raw history input: Mean of **W**^V,hist^ over presynaptic cells.
- History input familiar: Mean of 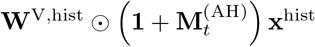 over all encoded times ≤ 1.25 seconds (i.e. those that would be passed to the network during a single omission), where 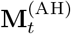 is measured at the beginning of the familiar imaging session.
- Raw history input, no omissions: Mean of **W**^V,hist^**x**^hist^ over encoded times when omissions are not present, i.e. ≤ 0.5 seconds. Correlation results do not differ significantly for any modulated version of this metric.
- Raw history input, omissions: Mean of **W**^V,hist^**x**^hist^ over encoded times that only occur during omissions, i.e. 0.5 *< s* ≤ 1.25 seconds.
- Raw Exc. + history input: The sum of “raw excitatory input” and “raw history input” above.
- Variance explained (familiar): Variance explained of all kernel fit during the familiar session.
- Variance explained (novel): Same as above for the novel session.
- Variance explained (novel plus): Same as above for the novel-plus session.
- Noise inject: Amount of time-correlated noise injected into cell, i.e. **w** in Eq. (21).
- Raw SST input: Mean of **W**^V,S^ over presynaptic cells.
- Raw VIP input: Mean of **W**^V,V^ over presynaptic cells.
- Bias: Value of **b**^V^. Since biases are the same for all cells of a given population, this cannot correlate with any coding scores but is provided for completeness.

## Acknowledgements

We thank Anton Arkhipov, Stefan Berteau, Darrell Haufler, Shinya Ito, Lukasz Kusmierz, Zhixin Lu, Alex Piet, Iryna Yavorska for feedback on this paper. SM has been in part supported by NSF 2223725, NIH R01EB029813, and RF1DA055669 grants. We also wish to thank the Allen Institute for Brain Science founder, Paul G. Allen, for his vision, encouragement, and support.

## Author Contributions

Conceptualization, K.A., M.G., and S.M.; methodology, K.A., L.C., and S.M.; software, K.A.; formal analysis, K.A.; investigation, K.A.; writing – original draft, K.A.; writing – review & editing, K.A., L.C., M.G., S.O., and S.M.; visualization, K.A.; supervision, M.G., S.O., and S.M.; funding acquisition, S.M..

## A Additive modulations

An alternative form of modulations that has also recently been considered has the modulations directly added to the fixed weights rather than the multiplicative form we consider throughout this work [35, 36]. Explicitly, rather than the transformation of the form of Eq. (1), these modulations affect the fixed weights via

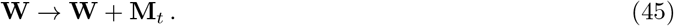

The same modulation update expressions continue to be used: for the associative case, Eq. (2a), and pre-only, Eq. (2b). Equivalent modulation bounds to those in Eq. (7) are enforced such that the associative and pre-only modulations do not exceed biological bounds of LTP/D and STSP, respectively. Many equivalent FMSN results to those in the main text and SM using this additive modulation are shown in Fig. S11.

## B Theoretical intuition

Here we discuss some analytic properties of FMSs that help us understand them better. Throughout this section, we will investigate both the multiplicative modulations considered in the main text as well as the additive modulations that were introduced in SM Sec. A. The analytical approximations for the additive modulations and their connection to Hopfield networks is more straightforward to understand, but similar qualitative results hold for the multiplicative modulations. Like the analysis here, feedforward version of Hopfield networks as familiarity discriminators were explored in Ref. [32], though their setup requires specialized weights that take on the value of the input neurons to compute the energy function used for familiarity discrimination.

Like the main text, let **M** be the modulation matrix and **W** be the fixed matrix of synaptic connecting the input and the output. Let **x**_1_, …, **x**_*m*_ be the set of familiar inputs we would like the network to memorize. Additionally, let 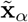 for *α* = 1, …, *m* be novel inputs, that also obey 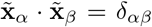 and 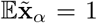 for all *α, β* = 1, …, *m*, but also 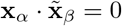 for all *α* and *β*. Here we assume the novel and familiar sets are of the same size, but it is straightforward to generalize what we show here for different size sets. Note since we consider inputs that are positive definite, in practice the dot product between any two inputs is finite, but it can approach zero as the size of the input space gets large and the sparsity is small.

To begin with, we establish some properties of the element-wise product that will be useful for the multiplicative form of the modulations. We will often make use of the identity that shows how an outer product of vectors (e.g. the modulations) act through matrix multiplication,

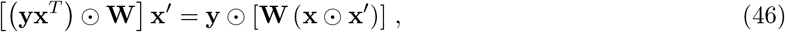

from Ref. [36]. Additionally, it will be useful to compare the (L2) magnitudes of the element-wise product between two stimulus vectors. Let nonzero values of the vectors be *A* with probability *p*. If the two vectors are different we have

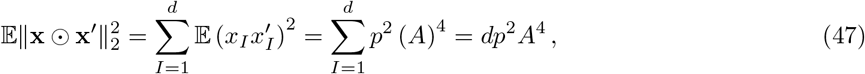

where we have used the fact that the only nonzero element of the element-wise product occurs when the Meanwhile, if the two stimulus vectors are the same, instead we have

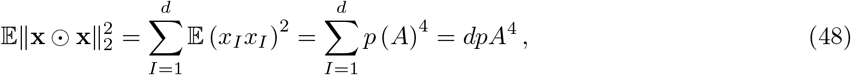

which is larger by a factor of *p*. Note a similar property holds for the dot product between two vectors, where 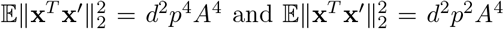, but now the same vector result is larger by a factor of *p*^2^.

Thus, in the limit that *p* → 0 and proper normalization of *A*, we make the analogous approximations

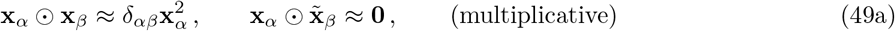

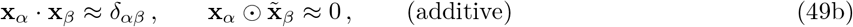

where we use the shorthand 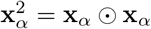.

Let us start with an approximate setting that will serve as a rough representation for the function of the FMSN. We assume that the modulations are *not* involved in the the feedforward pass of our network but are still updated as we have described in the main text. That is, the output activity is given by **y** = *ϕ* (**Wx** + **b**) but we still update **M**_*t*_ via Eq. (2a) even though it has no effect on the network’s behavior. We also presume the modulations do not decay, i.e. *λ* = 1, and that modulation are small enough such that they do not encounter the biological bounds or violate the bounds of Eq. (5).

Over a training time where the familiar inputs are each shown *N* times, the associative update will lead to an expected **M** given by a sum of *m* outer products

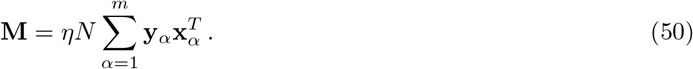

Notably, if **W** = **I** and **y**_*α*_ = **x**_*α*_, i.e. if *ϕ*(*x*) = *x*, this would be the same form of the updates to the lateral connections in a Hopfield network with an associative learning rule. Now consider how this modulation matrix acts on a given familiar input (for next three equations, top: multiplicative, bottom: additive),

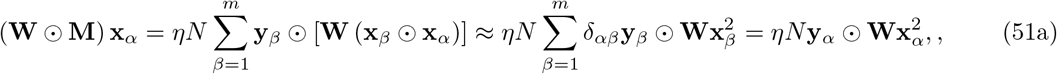

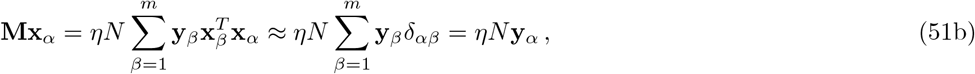

where for the multiplicative case (top) we have used Eq. (46) for the first equality and in both lines we have used the approximation of Eq. (49). Now compare this to a novel input

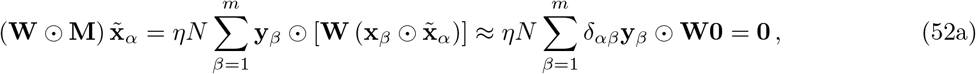

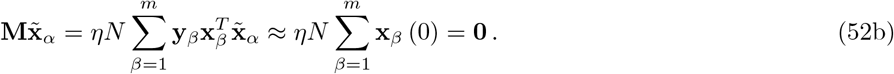

Thus a familiar input yields a non-zero modulation but a novel input simply yields zero. From the above results, we have

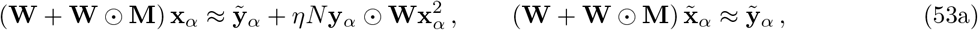

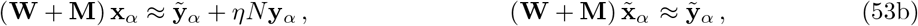

Thus we see that if *η >* 0 (or *η <* 0) the familiar preactivations grow (shrink) in size from the effects of the modulations, while the novel preactivations are left approximately unchanged.

Let the *familiar subspace* be the subspace of the input stimulus space spanned by the familiar stimuli. Then, since any stimulus can be decomposed into parts that lie within the familiar subspace and perpendicular to it, a generalization of the above arguments shows that the modulation matrix **M** will yield a nonzero result for any vector that has components in the familiar subspace. Since for a large input stimulus space the novel inputs are close to perpendicular to the familiar subspace (so long as *m* ≪ *d*), they yield approximately zero output.

For the additive case, we can see Eq. (53) is similar to checking the energy function of a Hopfield network (up to the vector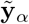), which has been used previously as a method of familiarity detection [32]. Indeed, since the activity in our network is positive-definite, taking the L1 normalization of this output is equivalent to taking the dot product with **1**, so it is similar to a Hopfield energy measurement with one occurrence of the stimulus replaced by **1**.

Now in practice, the modulations are involved in the forward pass, so as the modulations get updated during training they affect the output. For the FMSN, **y** = *ϕ* [(**M**⊙ **W**) **x** + **Wx** + **b**] and thus the modulations affect its own update. Notably, it is only the *output* activity that is affected by our approximation above, and so what will change are the **y**_*α*_ dependence of Eqs. (50) to (53). However, what causes the significantly different behavior between Eqs. (51) and (52) is the *input* activity dependence of **M**, and this is unchanged when we include modulations in the forward pass.

## C Strengthening versus weakening

Here we briefly discuss differences in the effectiveness of developing distinct responses to the familiar and novel sets in the FMSN from strengthening and weakening modulation mechanisms. We first study the FMSN in the setting we investigated in the main text, namely networks with only excitatory synapses in the input layer. As a result, a strengthening (or weakening) of the synapses results in a larger (or smaller) output neuron response.

There is a significant difference in the evolution of the **M**_*t*_ as a function of training time in identical FMSNs that simply differ in the sign of *η* (Fig. S3b). Notably, **M**_*t*_ at roughly the same rate for the first 8 training steps, but for the weakening mechanism the growth of **M**_*t*_ drops significantly after these first few times steps. This is a result of the output activity being smaller, which means the updates to **M**_*t*_ are smaller (Fig. S3c,d).

We can also see the effect the biological bounds have on the modulation growth. In particular, for the strengthening mechanism, the evolution of **M**_*t*_ differs significantly with and without the biological bounds (Fig. S3e,f). Notably, the weakening mechanism isn’t as strongly affected by said bounds. The fact that weakening has been observed down to 20% for associative and only 200% for strengthening makes the former much more effective. This is a simple comparison of ratios of novel to familiar: for weakening, 20% leads to a ratio of 1*/*0.2 = 5 whereas strengthening to 200% leads to a ratio of only 2*/*1 = 2.

Another major difference between the strengthening and weakening mechanisms is the way the biologically-motivated non-linearity, Eq. (11), acts on preactivation values. Since strengthening excitatory connections can only increase a neuron’s output, but said output is bounded by the non-linearity, eventually the strengthening yields diminishing returns in terms of how much a given output can change. Meanwhile, weakening excitatory connections can push a neuron below its firing threshold, completely cutting off a neuron’s response. We see that the evolution of preactivation values is fairly comparable between the two mechanisms (Fig. S3g).

Thus we have seen two major factors that cause the weakening and strengthening of excitatory synapses to differ: (1) a difference in the bounds of said changes from experiment and (2) an asymmetry in the FMSN of how larger/smaller output activity is handled through the neuron’s non-linearity as well as the the modulation updates. Of course, we have only considered plasticity mechanisms on synapses belonging to excitatory neurons thus far. For inhibitory synapses, a strengthening (or weakening) of the synapses results in smaller (or larger) output neurons response, the opposite effect of the excitatory synapses. Thus we can investigate if the FMSN has different behavior when introduce inhibitory neurons in the input population and then make the inhibitory synapses FMSs.

For direct comparison to the FMSN with only excitatory synapses, we assume inhibitory plasticity obeys similar bounds to what we use for STSP and LTP/D effects [20, 39]. We compare the behavior of an FMSN with both excitatory and inhibitory neurons in its input when either the excitatory or inhibitory neurons have a strengthening FMS mechanism on them (Fig. S3h). Consistent with our findings above, we find that strengthening of the inhibitory neurons is more effects of at separating the novel and familiar distributions than a strengthening of excitatory neurons (Figs. S3i,j). Since the bounds of the two FMS mechanisms are identical, this difference is caused by the neuron’s non-linear behavior discussed in point (2) above.

## D Additional figures and tables

**Figure S1:**
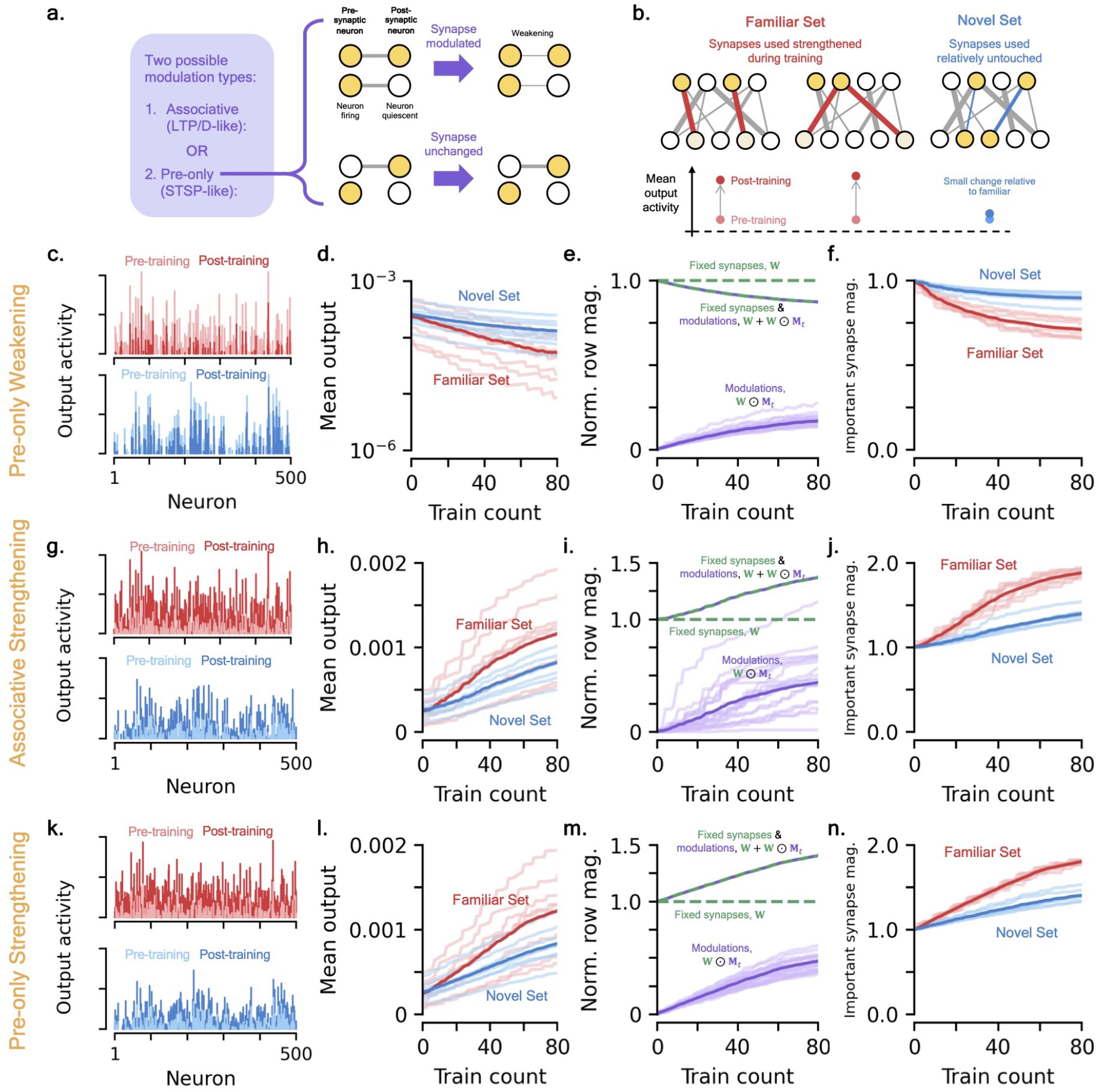
Pre-only weakening and associative strengthening FMSs and their behavior in the FMSN. **(a)** Schematic of pre-only FMS mechanism. **(b)** Strengthening FMS equivalent of Fig. 2c. **[c-f]** *FMSN with pre-only weakening modulations*. Equivalent associative plots in Figs. 2[d-g]. **(c)** Example raw output response activity for a familiar (red) and novel (blue) stimulus pre- and post-training. **(d)** Change in mean output activity of the familiar and novel sets over training. The dark lines show the mean output activity across each stimulus set, the light lines show individual stimuli. **(e)** Relative mean row magnitude of the modulations, **W** ⊙ **M**_*t*_ (purple), the unmodulated weight matrix, **W** (green), and total synaptic strength (green and purple) over training. Dark lines show means while light lines shown individual rows. **(f)** Change in important synapse magnitude for familiar and novel inputs as a function of training time (Methods). **[g-j]** *FMSN with associative strenthening modulations*. Note many trends compared to Figs. 2[d-g] are reversed because of strengthening rather than weakening. Figures are equivalent to [c-f]. **[k-n]** *FMSN with pre-only strenthening modulations*. Figures are equivalent to [c-f].

**Figure S2:**
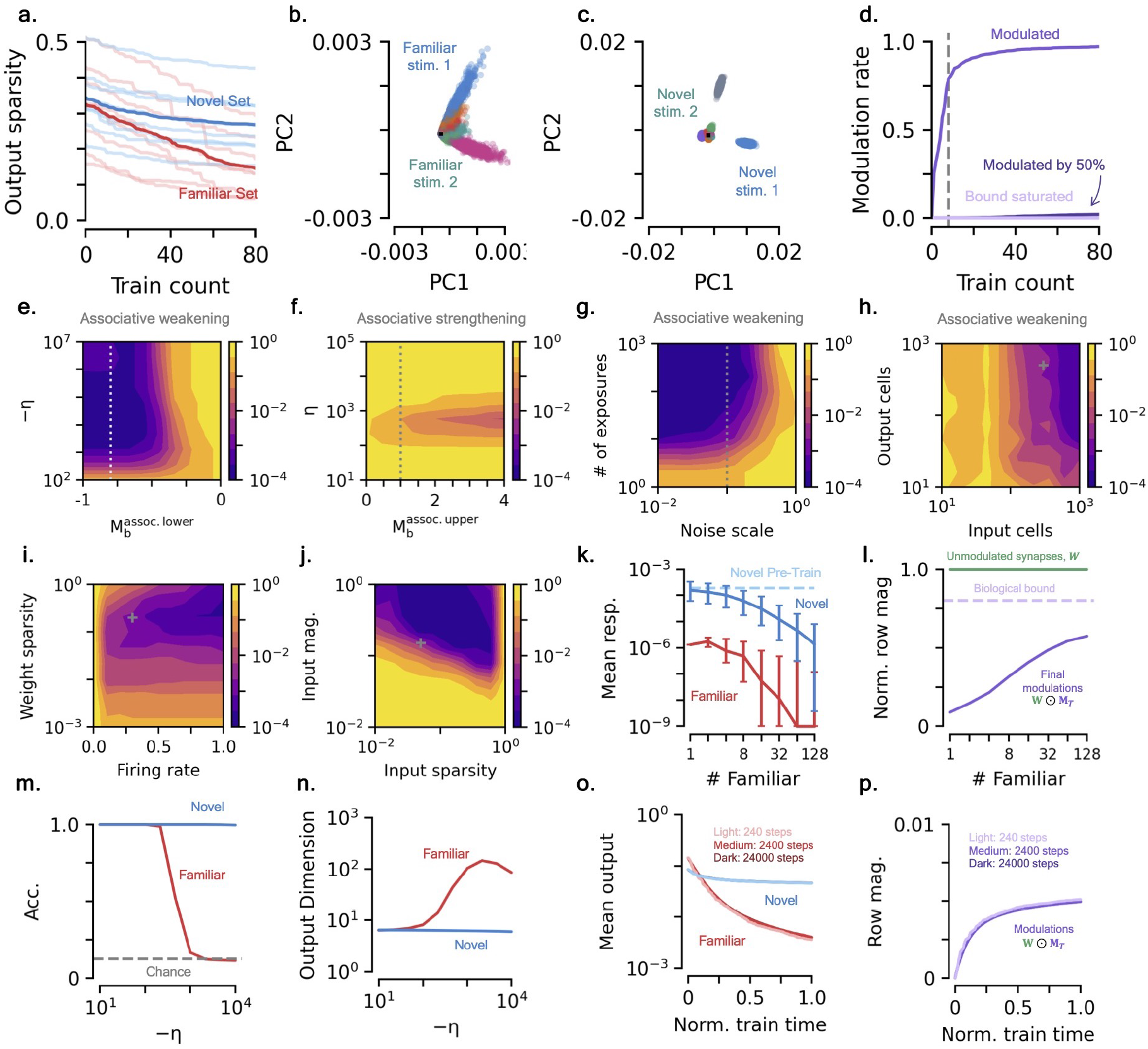
Additional properties of the FMSN. **(a)** Output sparsity of the novel (blue) and familiar (red) sets as a function of training count (Methods). Dark lines show mean across entire sets, light lines show individual members of sets. **(b)** PCA projection of output activity of the familiar set (Methods). Distinct colors show distinct members of set. **(c)** Same as (b) for the novel set. Note the different axis scale. **(d)** Modulation rate of synapses of the FMSN as a function of training time. Medium purple curve shows rate of synapses modulated at all, dark purple shows synapses that have been weakened by 50% of their initial value, and light purple shows synapses that have their biological bounds saturated (Methods). **[e-j]** *Contour plots of p-values from KS test as a function of various FMSN parameters, averaged over* 10 *initializations. Grey lines/marks represent values used in main text*. **(e)** Modulation bounds and learning rate. Specifically we consider associative weakening so we vary the lower bound that controls how weak a synapse can be made through modulations. **(f)** Same as (i) for associative strengthening and thus we vary the upper modulation bound. **(g)** Noise scale and number of exposures to each familiar stimulus. **(h)** Number of input and output cells. **(i)** Synapse sparsity and firing rate at initialization. **(j)** Random binary vector sparsity and magnitude. **(k)** Post-training median response (bars: 10th and 90th percentiles) of the FMSN’s response to familiar (red) and novel (dark blue) sets as a function of the size of the familiar set. Pre-training novel response shown for reference (light blue). Each familiar stimulus is shown 10 times, so training times increase for larger familiar sets. Responses thresholded to 10^−9^ for visibility on plot. **(l)** Size of post-training modulations (dark purple) as a function of familiar set size, with unmodulated synapses (green) and biological bounds (light purple) shown for reference. **(m)** Accuracy of decoding the familiar and novel sets as a function of learning rate (Methods). **(n)** Output dimensionality of familiar and novel sets as a function of learning rate (Methods). **[o, p]** *Equivalence of FMSN training for different sequence lengths and learning/decay rates*. See Sec. 4.4.7. **(o)** Mean output magnitude of familiar and novel sets as a function of normalized train time. Different colors vary the training time, learning rate, and decay rate, see Eq. (33). **(p)** Same as (o), but modulation magnitudes.

**Figure S3:**
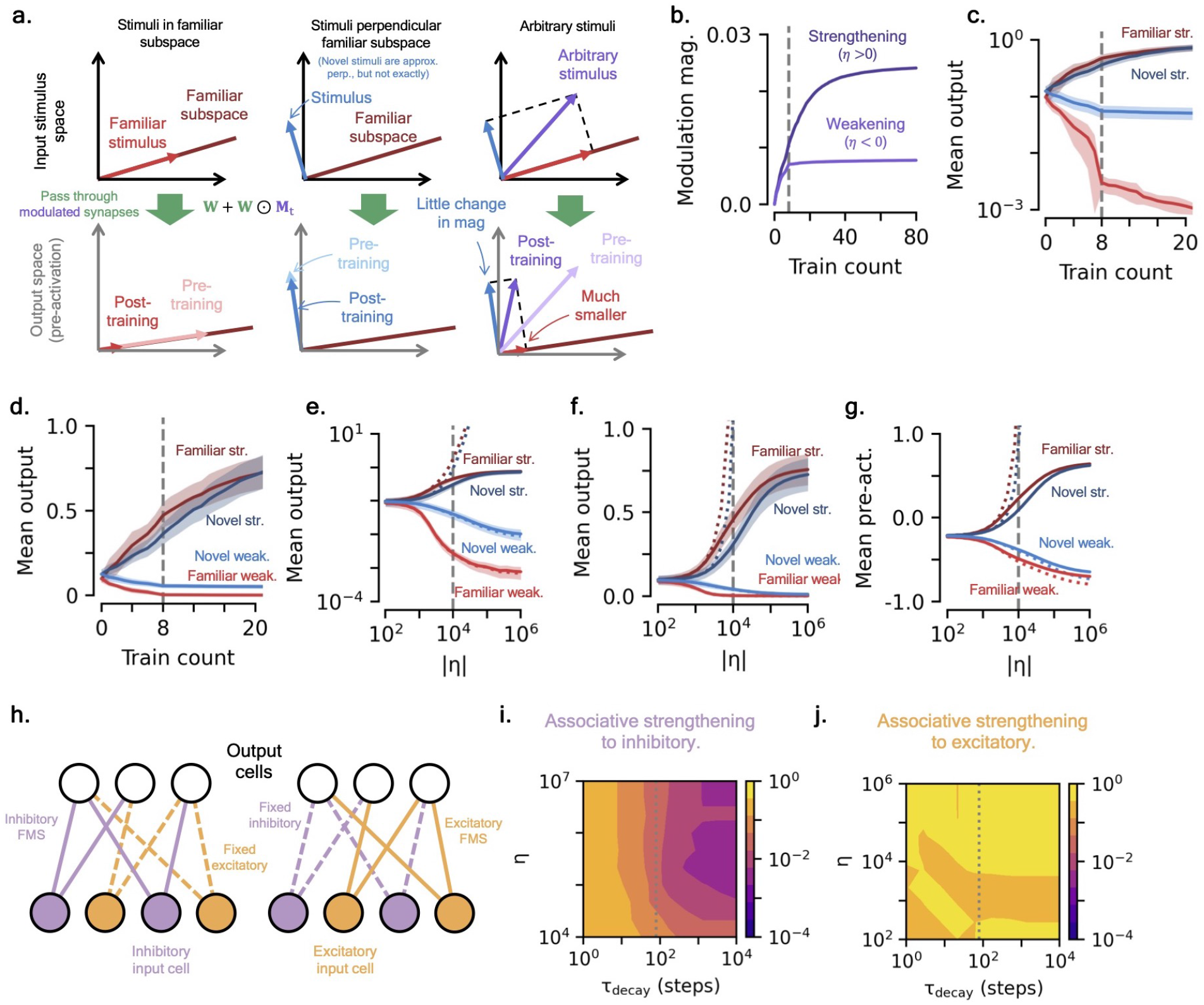
Familiar subspace and FMS strengthening versus weakening. **(a)** Visualization of familiar subspace and how it affects the output magnitudes of different stimuli pre- and post-training. **(b)** The evolution of mean modulation row magnitude for identical networks, one whose modulations are strengthened (dark line, *η >* 0) and another whose modulations are weakened (light line, *η <* 0), as a function of training count. **(c)** Mean output magnitude of the familiar (red) and novel (blue) sets as a function of training count for an FMS mechanisms that strengthens (dark lines) and weakens (light lines) the synapses. Vertical grey line shows time where all members of familiar set have been shown once. **(d)** Same as (c), but *y*-axis is now a linear scale. **(e)** Same as (c), but mean output magnitudes after all members of the familiar set have been shown once (i.e. 8 training steps) as a function of learning rate size. Dotted lines show results without modulation bounds of Eq. (7). Vertical grey line shows learning rate used in (b) and (c). **(f)** Same as (e), but *y*-axis is now a linear scale. **(g)** Same as (e), but the pre-activation values of output. **(h)** Schematic of two FMSN setups with both excitatory and inhibitory in the input neuron population. The network on the left has FMSs on its inhibitory synapses, while the right one has FMSs on its excitatory synapses. **(i)** Contour plot of *p*-values from KS test between post-training familiar and novel mean output activity as a function of learning rate, *η*, and modulation decay rate, *τ*_decay_ for the network shown in (e) with inhibitory FMSs. The grey vertical line shows the timescale of the task, 80 steps. **(j)** Same as (f), but for the network with excitatory FMSs.

**Figure S4:**
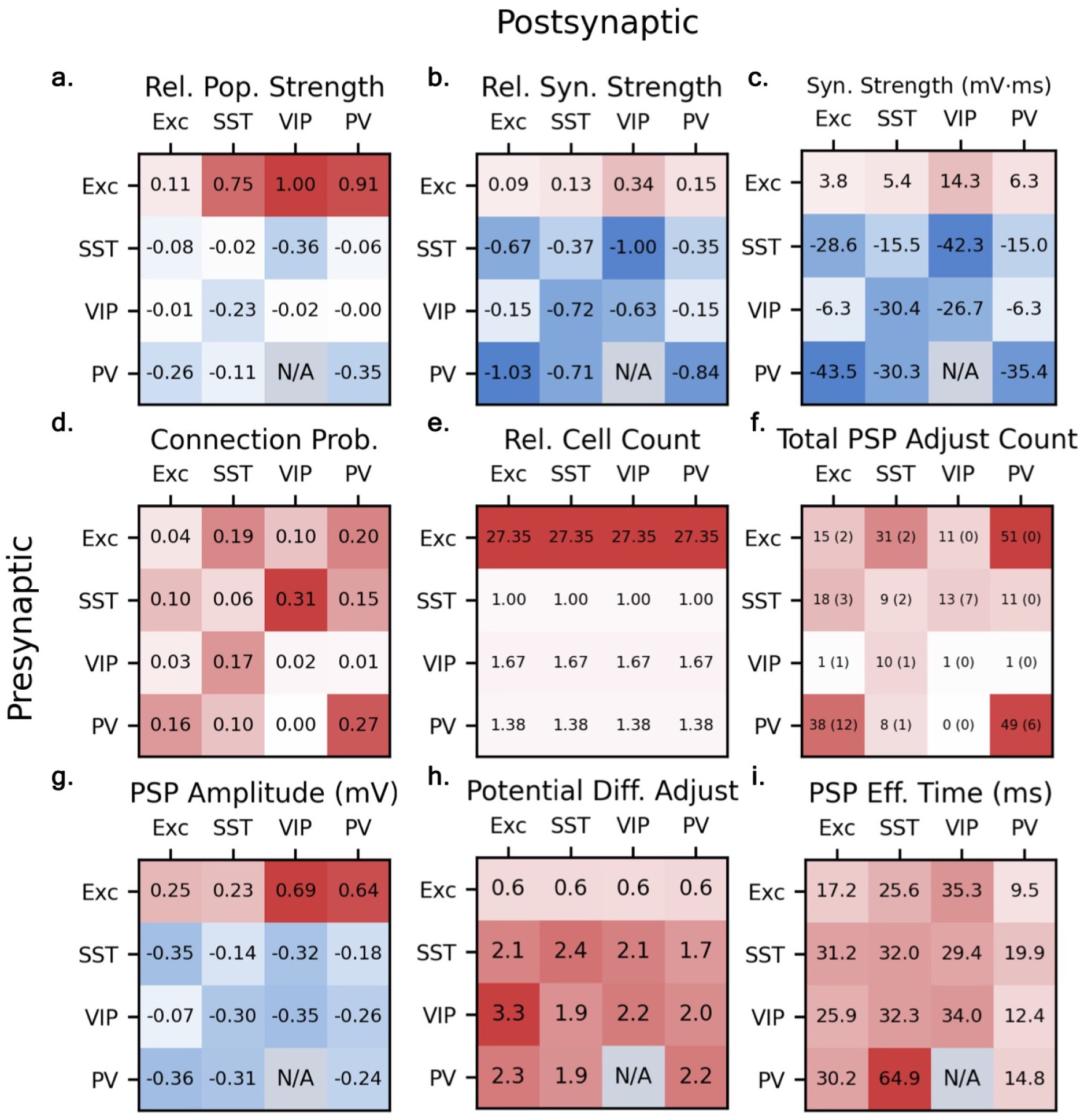
Microcircuit population strength contributions. Many metrics computed as a function of pre- and postynaptic cell type. All values are specific for layer 2/3 of the primary visual cortex. Although PV cells are not explicitly included in the cortical microcircuit model, we have included them here for completeness. “N/A” indicates synapses where no data was available, which correspond to cell-type connections that were exceedingly sparse in experimental analyses [20]. **(a)** Relative between-population connection strengths, computed via Eq. (22) and normalized by maximum magnitude. **(b)** Relative individual synaptic strength, as measured by time-integrated postsynaptic potential (PSP), see (c). **[c-e]** Three contributions to values shown in (a), see Eq. (22). **(c)** Mean time-integrated PSP, see Eq. (23). **(d)** Connection probabilities, from Ref.[20] (Methods). **(e)** Relative presynaptic cell counts (to SST cell count) from Ref. [18]. **(f)** Number of synapses used to compute time-integrated PSP. Number of omitted synapses from too large a *τ*_decay_ is shown in (*·*). Note counts reflect total number of synapses found in computing (d), so those connections with smaller probabilities of existing have less data. **[g-i]** Three contributions to *Z*^PSP^, see (c). **(g)** Amplitude of PSP pulse, from Ref.[20]. **(h)** Adjustment factor of PSP amplitude to account for potential differences, see Eq. (24). **(i)** Effective PSP time, computed from experimental PSP fits, see Eq. (26).

**Figure S5:**
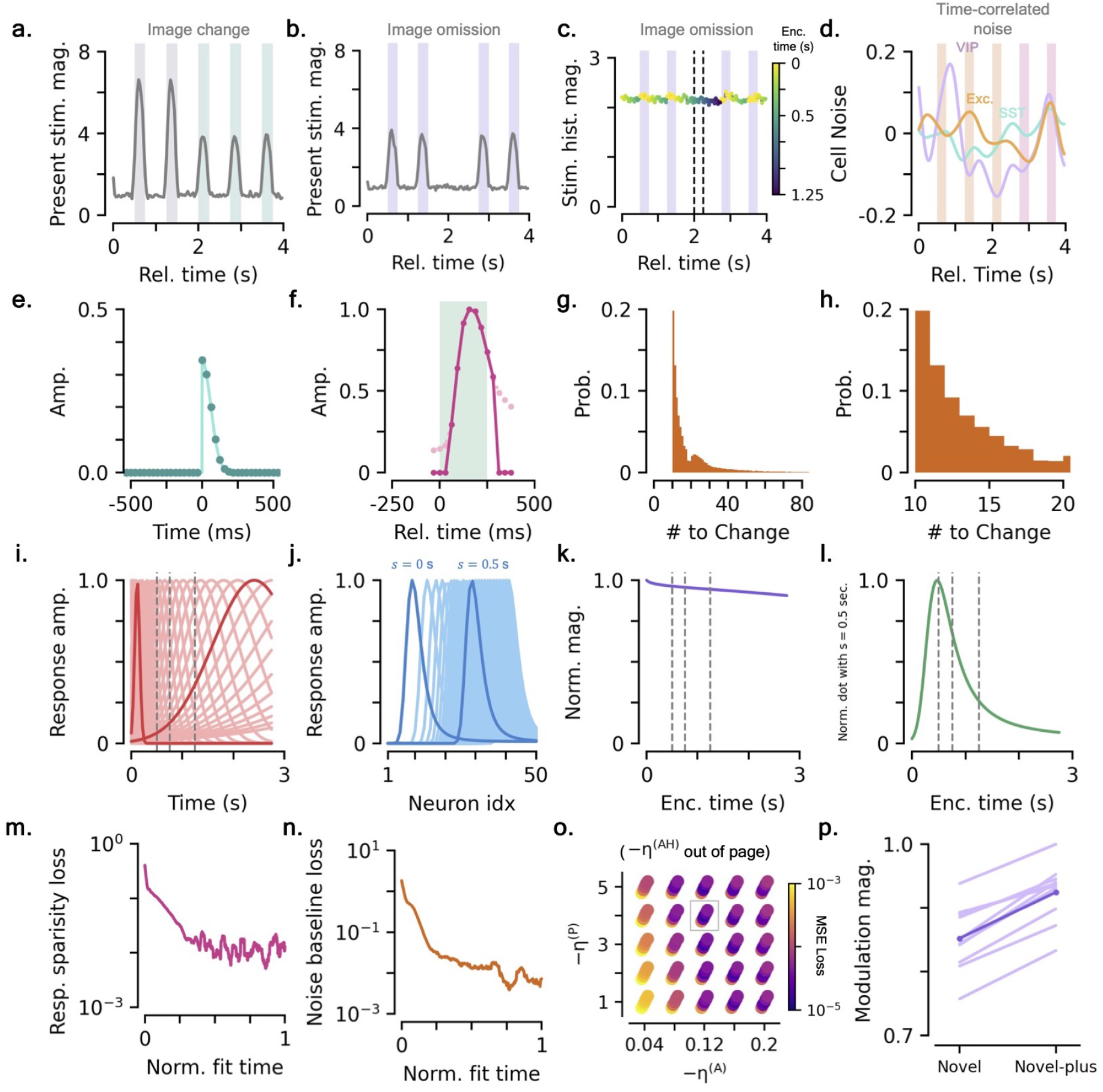
Additional microcircuit details. **[a-d]** *Microcircuit input details*. See Sec. 4.2.2 for details. **(a)** Magnitude (L1) of raw stimulus inputs during image change, 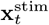, as a function of time. Shaded colored backgrounds correspond to distinct image presentations. **(b)** Same as (a), for an image omission. **(c)** Magnitude of stimulus history input, 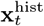, as a function of time. Discrete points are colored by the time since last image that they encode. **(d)** Exemplar time-correlated noise injections for one cell of each population. **(e)** Half-gaussian smoothing kernel applied to model and experiment cell responses, see Sec. 4.4.5 [24]. Discrete points show smoothing kernel values on discrete times separated by Δ*t* = 1*/*32 s. **(f)** Smoothing function (dark pink) used on model stimuli to gradually ramp up/down stimulus response, relative to the image presentation (green background). The smoothing function is the deconvolved mean excitatory trace, truncated to be the same number of time steps as the image width, with an additional offset, see Eq. 18. Light pink shows the non-truncated signal. **(g)** Experimental distribution of number of images from one change to the next. **(h)** Zoomed in version of (g). **[i-l]** *Stimulus history input details*. **(i)** Individual neuron tuning functions as a function of time since last image presentation (light) with particular neurons highlighted (dark). Grey vertical lines show times corresponding to onset of omission (0.5 s), end of omission (0.75 s), and onset of image after a single omission (1.25 s). **(j)** Neural population responses corresponding to various times since last image presentation, *s*, two times highlighted. **(k)** Magnitude (L1) of population response functions as a function of the time since last image presentation they encode. Grey lines same as (i). **(l)** Similarity of various population response functions to *s* = 0.5 seconds as a function of the time they encode. Grey lines same as (i). **(m)** Exemplar fit of response sparsity at initialization, see Sec. 4.4.2. **(n)** Exemplar fit of noise baseline at initialization, see Sec. 4.4.4. **(o)** Exemplar loss values from grid search over various FMS learning rates, box shows optimal values. **(p)** Change in modulation magnitude from start of novel to novel-plus imaging sessions.

**Figure S6:**
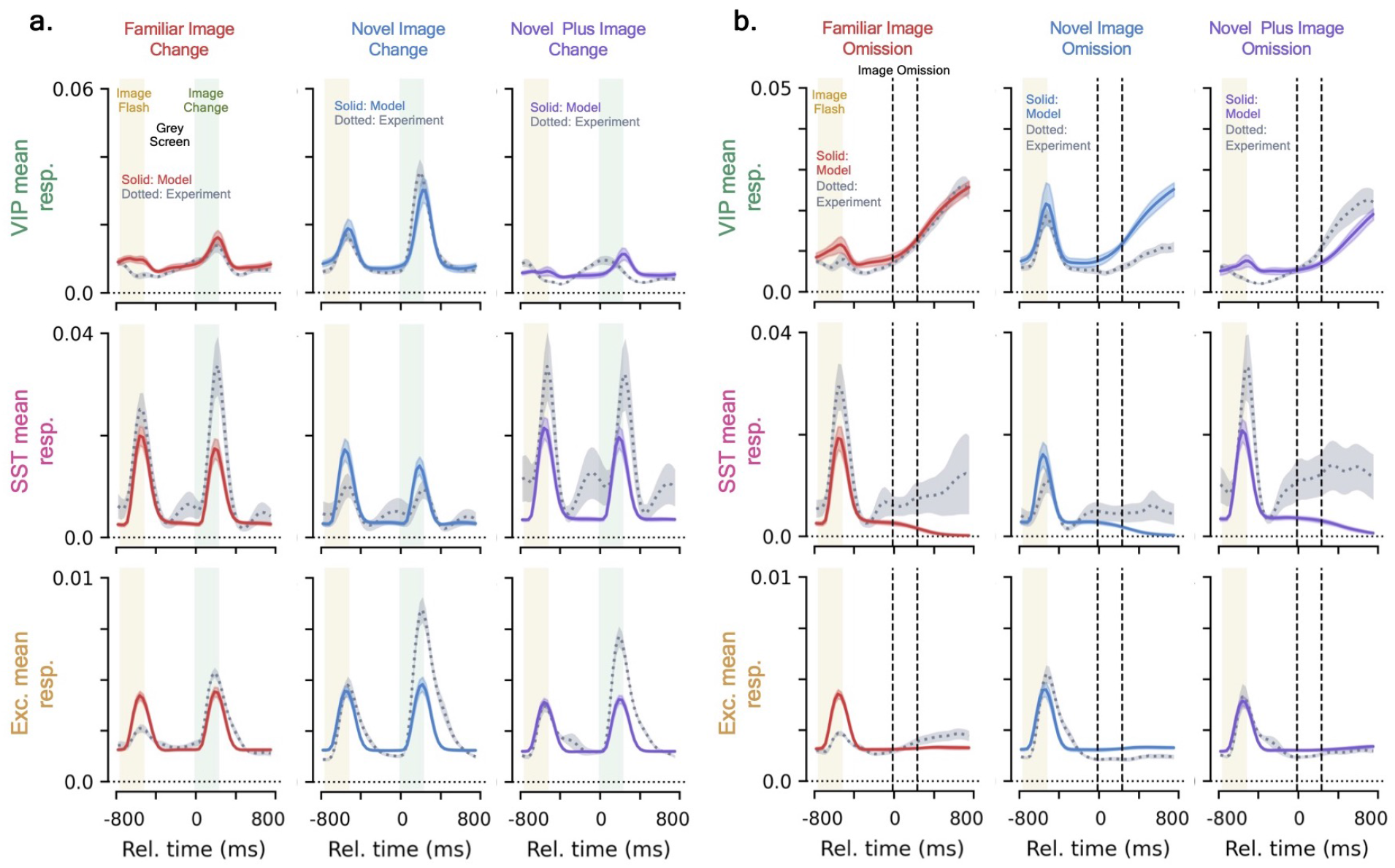
Microcircuit image change and omission mean responses. **(a)** Mean image change responses. Experimental results are shown as dotted grey line, model results are solid colored lines. Green background represents changed image being displayed, yellow background is pre-change image. Columns show different experience-levels (familiar, novel, novel-plus), rows show different cell populations (VIP, SST, Exc.). **(b)** Same as (a), but for mean image omission responses. Area between vertical dashed black lines represents time period where an image would usually be presented, but instead an image was omitted.

**Figure S7:**
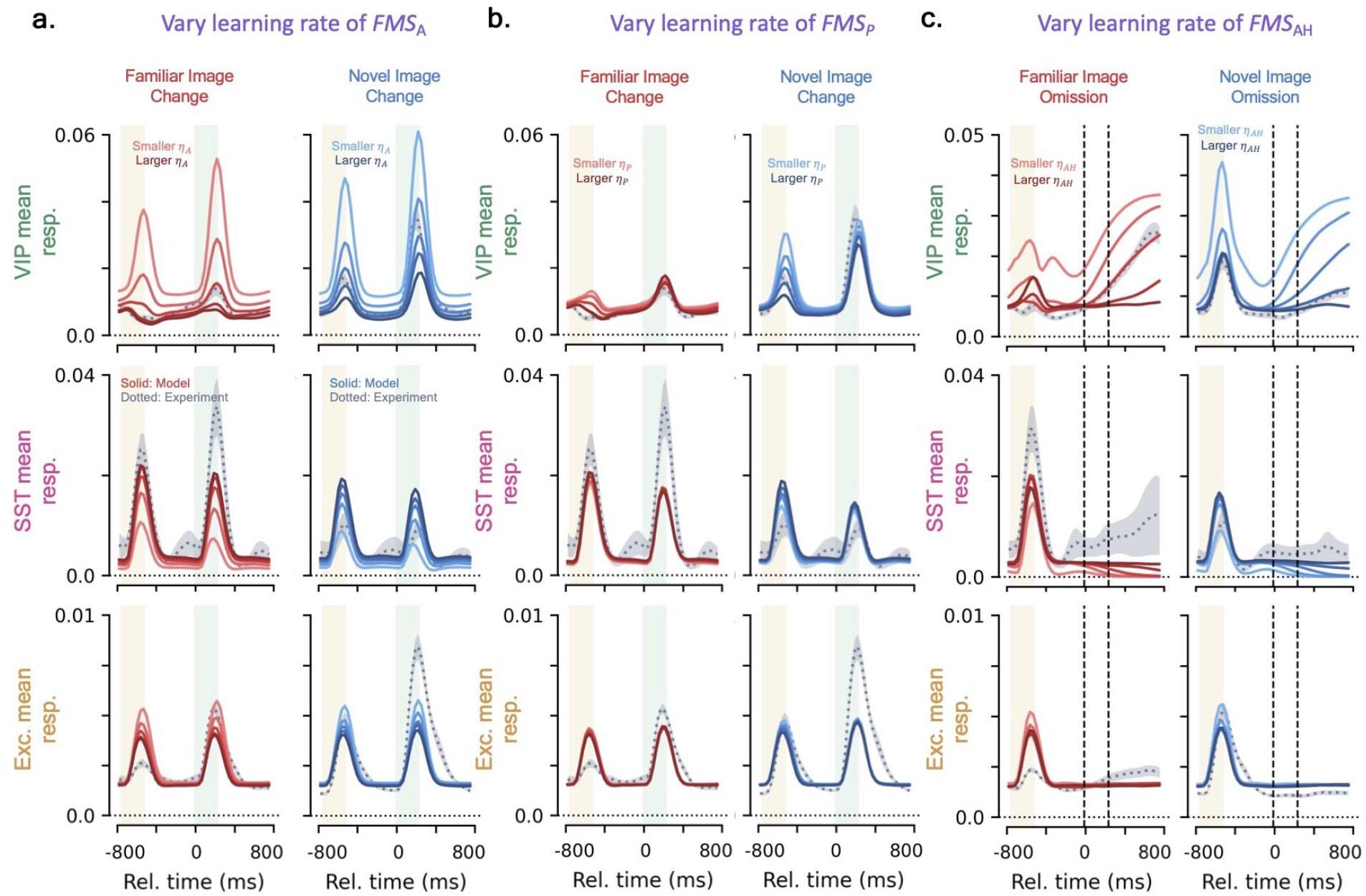
Microcircuit mean responses as FMS learning rates are varied. Darker lines show larger learning rates (i.e. stronger modulation) and lighter colored lines show smaller learning rates (i.e. weaker modulation). Notation is otherwise identical to that of Fig. S6. **(a)** Mean image change response as *FMS*_A_ learning rate is varied. Learning rates correspond to *η*_A_ = 0.04, 0.08, 0.12, 0.16, 0.20. **(b)** Mean image change response as *FMS*_P_ learning rate is varied. Learning rates correspond to *η*_P_ = 0.04, 0.08, 0.12, 0.16, 0.20. **(c)** Mean omission response as *FMS*_AH_ learning rate is varied.

**Figure S8:**
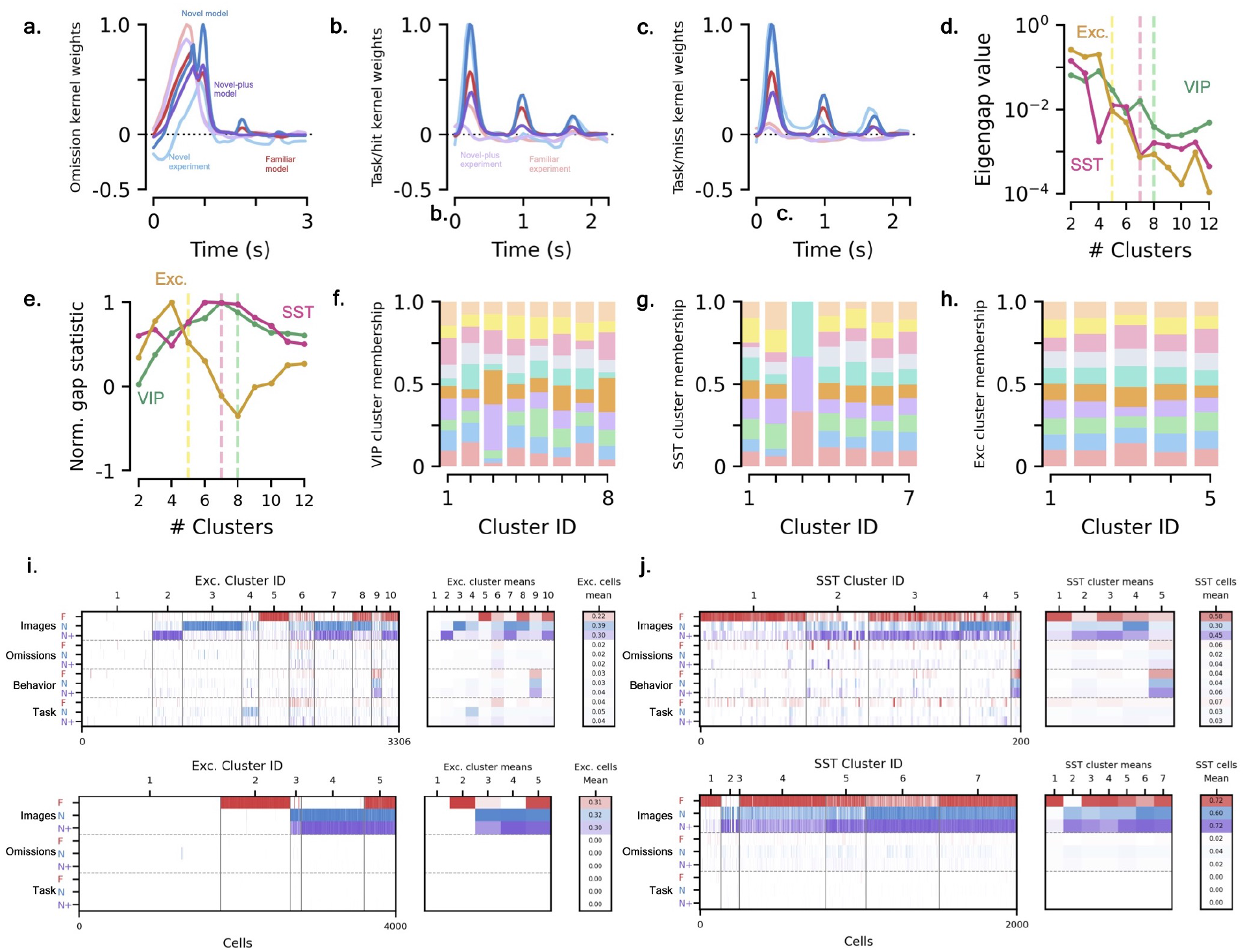
Additional cluster results. **[a-c]** *Feature kernel fits for VIP cells*. See Fig. 6d for image kernel results. **(a)** Omission kernel. **(b)** Task kernel, with experiment hit kernel. **(c)** Task kernel, with experiment miss kernel. **[d-e]** *Metrics to determine number of clusters*. Coding scores were clustered across 10 different intializations for each cell type. VIP (green), SST (pink), and excitatory (yellow) are shown. Vertical dotted line shows chosen cluster count. **(d)** Eigengap value as a function of the number of clusters, see Sec. 4.5.4 for details. **(e)** Gap statistic, normalized by maximum magnitude, as a function of the number of clusters. **[f-h]** *Cluster membership percentages across different network initializations*. Clustered results are gathered over 10 different network initializations and the percent of cells that of a given clustered that belong to a particular initialization are shown by distinct colors. Similar to experimental data, distinct initialization do not explain the distinct clusters. **(f)** VIP population clusters. **(g)** SST population clusters. **(h)** Excitatory population clusters. **[i-j]** *Clustered SST and VIP population coding scores*. **(i)** Excitatory population. **(j)** SST population.

**Figure S9:**
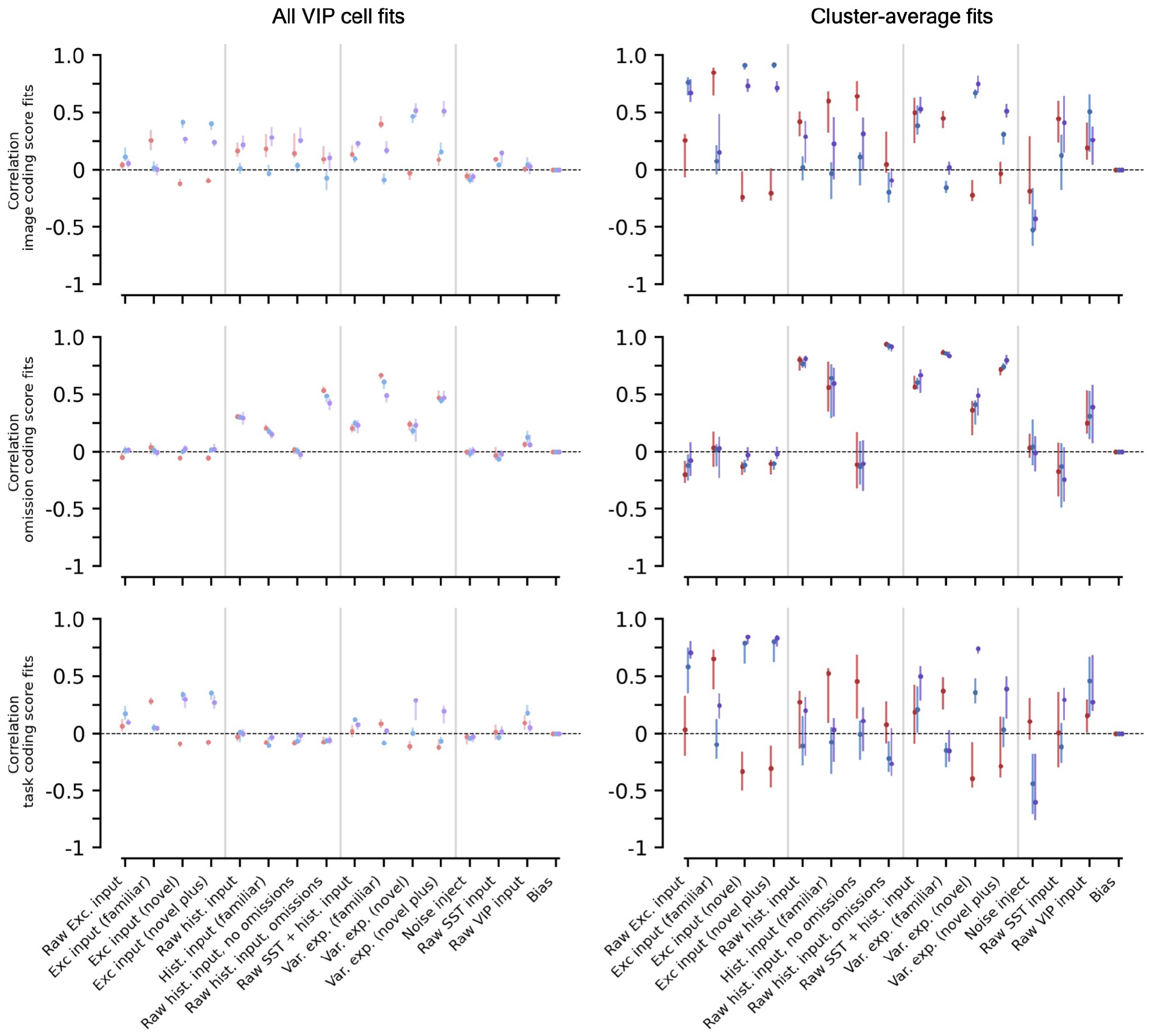
How well network metrics explain VIP cell coding scores and coding score clusters. Left column is fits over all VIP cell coding scores, right column is fits to cluster-averaged coding scores. Red, blue, and purple corresponding to the coding scores in the familiar, novel, and novel plus sessions, respectively. Top, middle, and bottom row are the image, omission, and task coding scores, respectively. Dot is median value across initializations, error bar shows Q1 and Q3 values.

**Figure S10:**
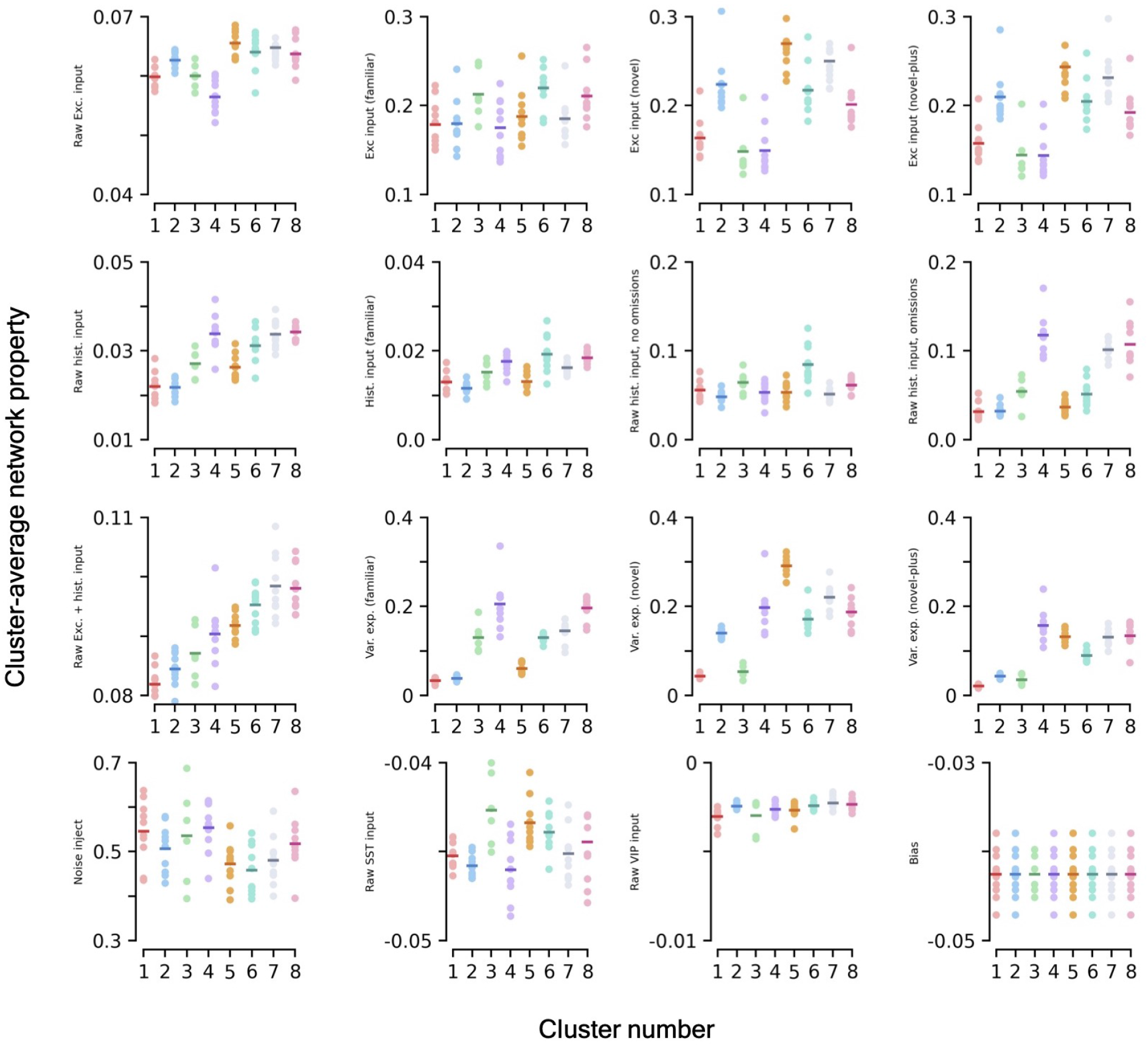
Network metrics as a function of VIP cell cluster. Various network metrics (see Methods for details) as a function of the VIP cluster number. Light dots show different networks, dark lines are mean across trials. Note clusters are sorted by mean coding score vector magnitude (smallest to largest). Note that if a given network initialization has less than 5 cells in a given cluster, its data is omitted.

**Figure S11:**
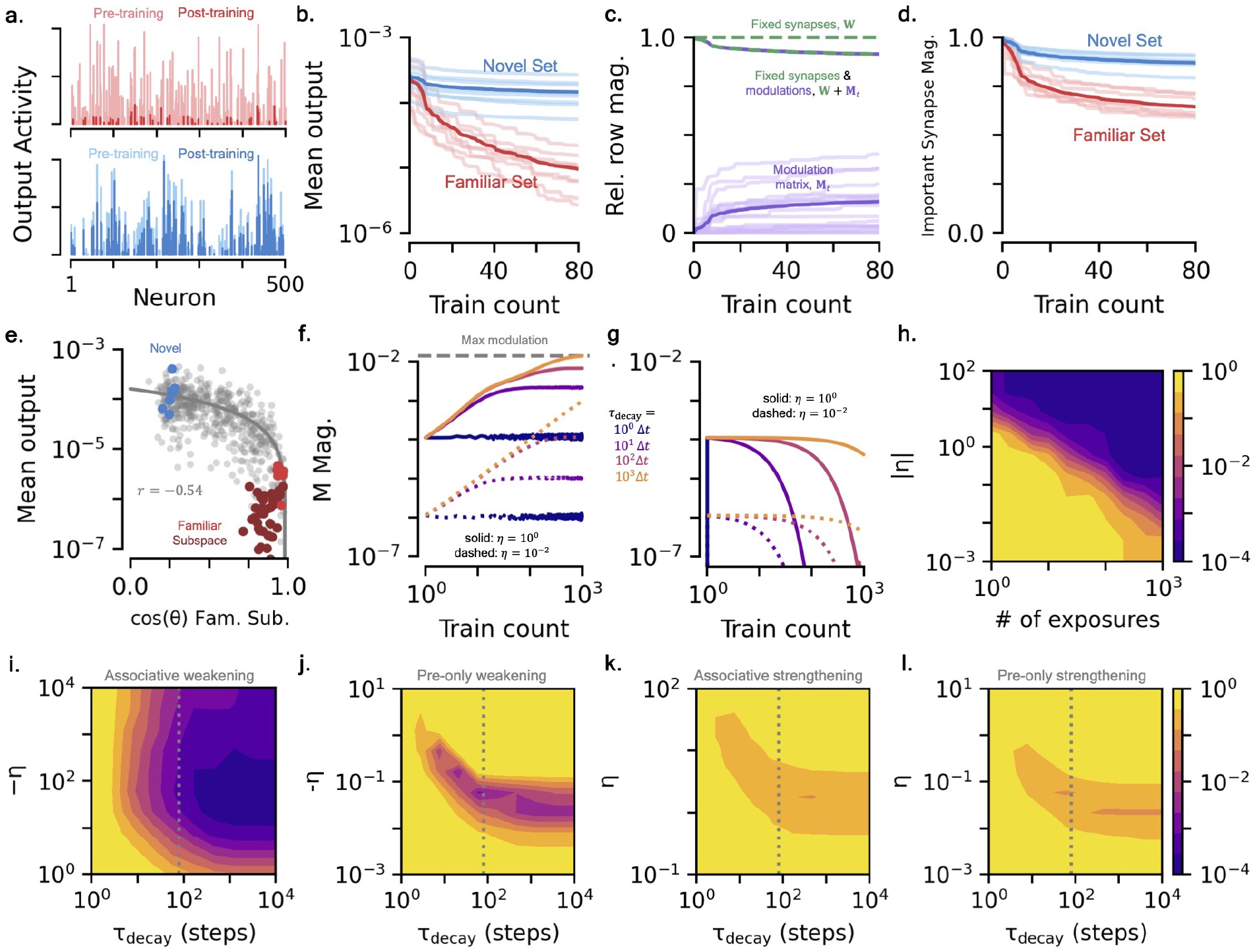
FMSN with additive modulations. **[a-d]** *Equivalent plots to Figs. 2[d-g]* **(a)** Exemplar raw output response activity for familiar (red) and novel (blue) pre- and post-training. **(b)** Change in mean output of novel set (blue) and familiar set (red) over training. **(c)** Change in synapse and modulation magnitudes over training. **(d)** Important synapse magnitude for familiar and novel sets over training. **[e-l]** *Equivalent plots to Figs. 3[a-h]* **(e)** How the distance of a stimulus from familiar subspace influences the FMSN’s mean output. **(f)** Change in **M** magnitude over training as learning rate and decay rate are varied. **(g)** Same as (f), but shows decay of **M** magnitude. **(h)** Distinguishability of familiar and novel sets post training (KS-test *p*-value) as a function of learning rate and number of familiar exposures. **(i)** Same as (h), scan now over FMS parameters *η* and *τ*_decay_, for associative weakening FMS. **(j)** Same as (i), for pre-only weakening FMS. **(k)** Same as (i), for associative strengthening FMS. **(l)** Same as (i), for pre-only strengthening FMS.

## Footnotes

1 In this setup, cell-type (excitatory versus inhibitory) only influences the sign of weights leaving a population. Since the activity of the output neuron population is directly measured, results here hold for either excitatory or inhibitory output neurons. An excitatory input population was chosen for simplicity, see the SM for the equivalent setup with inhibitory neurons as well.

2 To evaluate the novel activity pre-training without it becoming “familiar” to the network, we treat as we would a test set and do not modulate the synapses from its activity via Eq. (2a). The FMSN then has no memory of being exposed to it. We emphasize this is done solely for the sake of comparison to the familiar set and is not a necessary step in training.

3 For this approximation, we have assumed that all familiar inputs are presented roughly the same number of times in a randomized order, as is done for the FMSN training. For cases where familiar stimuli are presented in an uneven manner, the network will respond most weakly to inputs it has been exposed to the most and those most recently presented, see the SM.

4 Parvalbumin (PV) expressing inhibitory neurons are not included in our cortical circuit model directly, though the inhibition they provide to the other populations is partially accounted for from the threshold adjustments at the model’s initialization (Methods). This important simplification is driven by the desire to build a minimal model of the data from [24], where excitatory, VIP, and SST neurons were recorded and furthermore from the fact that VIP cells do not receive strong input from the PV population (Fig. S4a). The blanket inhibition in our model is in part supported by the general lack of specificity of PV to excitatory connections [55], though more recent evidence points to some levels of specificity [56].

5 The primary purpose of this input is to give the microcircuit information about the recent stimulus history. A simple neuronal circuit that counts the time steps since the last stimulus presentation, e.g. an RNN, could represent the higher cortical areas that may produce this additional input directly from the bottom-up present stimulus input.

6 We do not attempt to model the suppressed omission ramping that is observed in the VIP population during the novel session that gradually returns to familiar levels in the novel-plus session [24]. See Discussion for potential mechanisms which can model this effect.

7 An alternative form of the modulations considered in Refs. [35, 36] uses an additive modulation, rather than the multiplicative one we consider here. Essentially all results used for the FMSN generalize to this form of modulations as well (Fig. S11). See SM for additional discussion.

8 Since we use the FMSs to model several distinct types of synapse modulations that have their own vocabulary for synapse changes (e.g. depression and facilitation for STSP versus depression and potentiation for more long-term effects), we use “strengthening” and “weakening” as a general terminology that applies across the individual mechanisms the FMSs may model.

9 Results do not differ significantly from bit-flipped noise, both methods increase the dot product between two randomly drawn stimuli, making the familiar stimuli harder to distinguish from the novel stimuli.

10 In practice, smoothing the present stimulus signal from the L4 excitatory response would have been more realistic. However, the depth differences between L2/3 and L4 did not change the excitatory response significantly, so we have just used L2/3 for simplicity.

11 Any two vectors drawn from the sparse random binary vector distribution we consider in this example have cosine similarity of 0.14. Cosine similarity of stimuli decreases with increased sparsity and larger input dimension.

## Notes

### Competing Interest Statement

The authors have declared no competing interest.

### Summary of Updates

Title change; author affiliations updated; abstract updated; minor clarifying points throughout.

